# Twist-Dependent Elastic Limits and Unwinding of Tightly Bent DNA

**DOI:** 10.1101/2024.02.14.579968

**Authors:** Soumya Chandrasekhar, Emma Meyer, Fatemeh Fadaei, Rachel Bricker, Thomas P. Swope, Morgan Schreck, Mark Macri, Draven Houser, Daniel R. Hollis, John J. Portman, Thorsten L. Schmidt

## Abstract

DNA is under high bending and torsional strain in many biological contexts ^1,2^, but elastic limits of tightly bent DNA under simultaneous torsional strain are not fully understood ^3–22^. We synthesized DNA circles with all possible radii of curvature between *r* ≈ 2.7 – 5.7 *nm* in increments of *Δr* = 0.05 *nm* and defined helical repeats ranging from *h* ≈ 10 – 13 base pairs per helical turn. Nuclease digest reveals that DNA can be bent to *r* ≈ 3.0 *nm* without kinking, but only when *h* < ~10.9 bp/turn, while underwound DNA kinks irrespective of curvature. Histone proteins overwind DNA and thereby mechanically stabilize it. The natural helical repeat *h*_0_ increases from 10.45 to >11 bp/turn due to twist-bend coupling, which is not caused by kinking before ligation. These findings require reassessing our models and the energetics of molecular mechanisms involving DNA under mechanical stress.

## Introduction

In all cells and viruses, DNA is twisted and bent in a highly dynamic way. For example, two meters of human DNA are tightly wrapped around histone proteins, compacting it into a nucleus measuring mere micrometers across. In eukaryotes and archaea, DNA is wrapped around histone proteins to radii of curvature of just ~4.5 nm to compact it. During DNA replication and transcription, DNA unbinds from the nucleosome and relaxes its mechanical stress. DNA is also tightly bent in nanometer-sized virus capsids, bacteria, or complexes with many transcription factors that regulate gene expression.

A structurally and mechanically accurate physical model of DNA is essential to understand the molecular mechanism and energetics of some of the most fundamental molecular mechanisms in biology including DNA packaging, replication, transcription and gene regulation ^1,2^.

The twistable wormlike chain (TWLC) is the standard elastic model for DNA and treats DNA as an inextensible rod with elastic moduli controlling twist and bend deformations ^23^. It is generally believed to accurately describe the mechanical behavior of DNA that is longer than the persistence length (*l*_*p*_~50 nm or 150 bp). Some studies postulated extreme bendability in small circles ^5,7,14^ and even short, relaxed DNA fragments ^9^, but others found alternative explanations ^8,10,15,16,18,20^ and that DNA remains elastic as described by the TWLC model. Motivated by this important biological question, many experimental ^4,7,8,11–13,17–19,21,21,22^ and theoretical and computational studies ^20,24–27^ explored at what radius of curvature DNA loses its elastic properties and develops sharp kinks. Most reports find DNA with a radius of curvature of nucleosomal DNA (r = 4.5 nm) or above to be stable, however, the influence of twist is not clear.

In addition to bending stress, torsional strain is another biologically important mechanical stress factor occurring during DNA packing, replication and transcription. In plasmid DNA, negatively supercoiled (unwound) DNA develops sharp kinks while positive supercoiling (overwinding) prevents kinking ^28^. Current assays cannot precisely and simultaneously control the radius of curvature *r* and twist. The twist in DNA can be quantified by the helical repeat *h* which is the average number of base pairs (bp) that are required to complete a full helical turn. In the absence of torsional stress, the “natural” helical repeat *h*_0_ of long, random sequences *h*_0_ = 10.45 (±0.05) bp/turn ^29–35^.

A second open question associated with mechanical stress is if *h*_0_ depends on curvature or not. While twisting and bending deformations are treated as independent by the standard TWLC model ^23^, Marko and Siggia predicted twist-bend coupling (TBC) ^36^, which would lead to an unwinding, or increase of *h*_0_ at small *r*. Until now, there has been no direct experimental quantification of *h*_0_ at small *r*. The torsional persistence length, *l*_*p*_, and *h*_0_ of DNA can be determined with circularization assays ^29–33^, but the usefulness of the assay is limited to fragments that are longer than *l*_*p*_ as transition states contain sharp kinks at nicks, which release bending and torsional stress (Supplementary Figure S1, Supplementary Note 1, ref. ^20,19,21^). Magnetic torque tweezer experiments for example require fragments where *L* ≫ *l*_*p*_ and cannot produce tight curvatures in a controllable way.

Here we describe a new ligation assay on nicked minicircles to answer both questions: 1) What combinations of bending (*r*) and torsional stress (*h*) lead to kinking, and 2) test if *h*_0_ changes at tight curvatures or not.

## Results and Discussion

### Nicked Minicircle Assay

To create a nicked minicircle, we first circularize a linear template strand by enzymatic splint ligation (Figure 1A-B). In the next step, a complementary strand (blue) is added (Figure 1C-D). The point where the 5’ and 3’ ends of the linear strand meet is called a nick and mechanically the weakest point in the structure. Nicked circles are also the last of many transition states in circularization assays (Supplementary Figure S1). Unlike linear fragments with sticky ends, this complex can neither unloop nor dimerize which reduces the impact of bending stress on complex formation (Supplementary Note 1).

**Figure 1.**
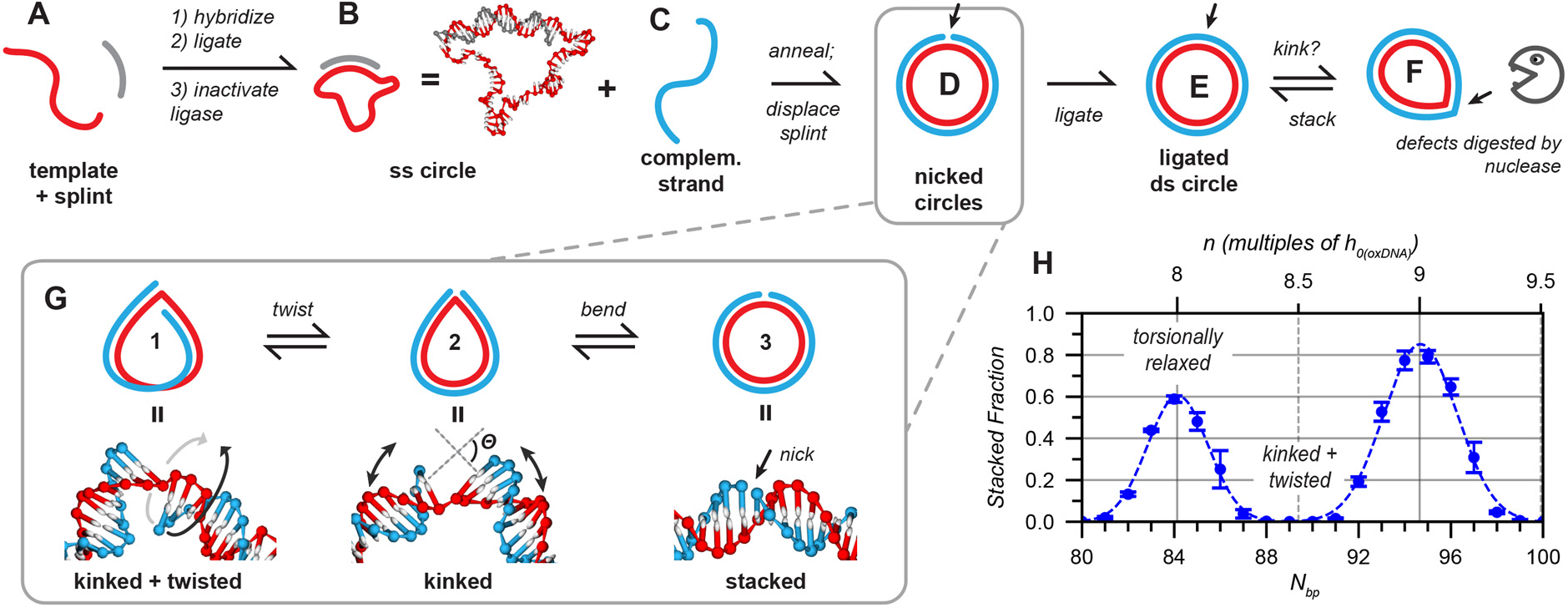
Nicked minicircle ligation assay. (**A-B**) Splint ligation of oligonucleotides of different lengths produces ss minicircles (**B**). The ligase is heat inactivated. oxDNA simulation snapshot illustrates flexibility of ss region. (**C**) Next, complementary strands are annealed to yield nicked circles (**D**), which can then be enzymatically ligated into double-stranded (ds) circles (**E**). (**F**) Circles with internal defects in circles can be enzymatically digested by nucleases. (**G**) Before the ligation, nicked circles can dynamically fluctuate between three different states: a kinked + twisted (**1**), kinked (**2**), or stacked conformation (**3**). Only the stacked conformation (**3**) can be experimentally ligated. Bottom: representative snapshots from oxDNA simulations, rendered with oxView (27, 37). (**H**) Plot of the fraction of stacked configurations from 21 different oxDNA simulations of nicked circles with circumferences between 80-100 bp. Dotted line: Gaussian fits.

The two ends of the blue strand are then enzymatically ligated by the addition of fresh ligase (Figure 1E). In the pictograms, the helical nature of the strands is omitted for simplicity but are twisted around one another with *Tw* turns. After ligation of the nicked strand, *Tw* must be an integer the same as the linking number (*Lk*), which is not formally defined for nicked strands. After ligation, *Lk* and any associated torsional strain cannot change anymore. Ligated circles containing structural defects such as kinks can be digested with certain nucleases.

### OxDNA Simulations

To get an understanding of the different states and dynamics of nicked circles, we simulated them with the coarse-grained model oxDNA, which is an excellent model for the mechanical behavior of DNA (38). The simulations reveal three distinct conformations: a “stacked” state; a “kinked” state; and a “kinked + twisted” state in which the ends are twisted out of alignment (Figure 1G). Stacked states are approximately circular or oval, whereas kinked configurations are sharply bent at the nick, which reduces the average curvature of DNA and therefore bending stress. The transition from a kinked to a stacked state is stabilized by additional *π*-stacking and hydrophobic effects between nucleobases but destabilized by increased bending stress. Nonetheless, the energy difference between these states is small compared to the energy required for looping a linear duplex of the same length.

We performed 21 different simulations with nicked circles with circumferences ranging from *N*_*bp*_ = 80 − 100 *bp*. The fraction of stacked configurations in each simulation was determined through a geometrical analysis of the bases around the nick (Figure 1G; Figure S3; Figure S4). Two maxima around *N*_*bp*_ = 84.1 *bp* and 94.6 bp appear roughly at 8 and 9 multiples of the natural helical repeat of oxDNA, *h*_0(oxDNA)_ = 10.55 *bp*/*turn* (Figure 1H) and *h*_0_ (averaged over multiple helical turns) is not significantly changed compared to a linear, relaxed dsDNA model. The minicircles are generally kinked when *N*_*bp*_ is a half-integer multiple of 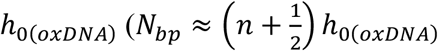 for integer *n*). These simulations show that the probability of finding a nicked circle in a stacked, ligatable conformation is strongly dependent on the twist of the DNA. Transitions between stacked and kinked conformations are in the nanosecond to microsecond range (Figure S5, Figure S6 Supplementary Movies S1-4), whereas lifetimes of both looped and linear configurations in classical circularization experiments with sticky ends can be on the order of seconds to minutes (ref. ^14,21^, Figure S1; Supplementary Note 1).

### Ligation and Nuclease Assay

Next, we experimentally assembled nicked minicircles with different *N*_*bp*_ and ligated them into ds circles with the strategy shown above (Figure 1A-E). We designed all template strands with the same sequences at the ends, so that one common 26 nt splint can be used for all strands (Figure 2A-B) and all circles share the same sequence around the nick. The sequences in between the common splint-binding domains are unique, randomized sequences, such that each ss circle can only hybridize to the correct counterstrand. Moreover, eventual sequence-dependent effects can be excluded this way.

**Figure 2.**
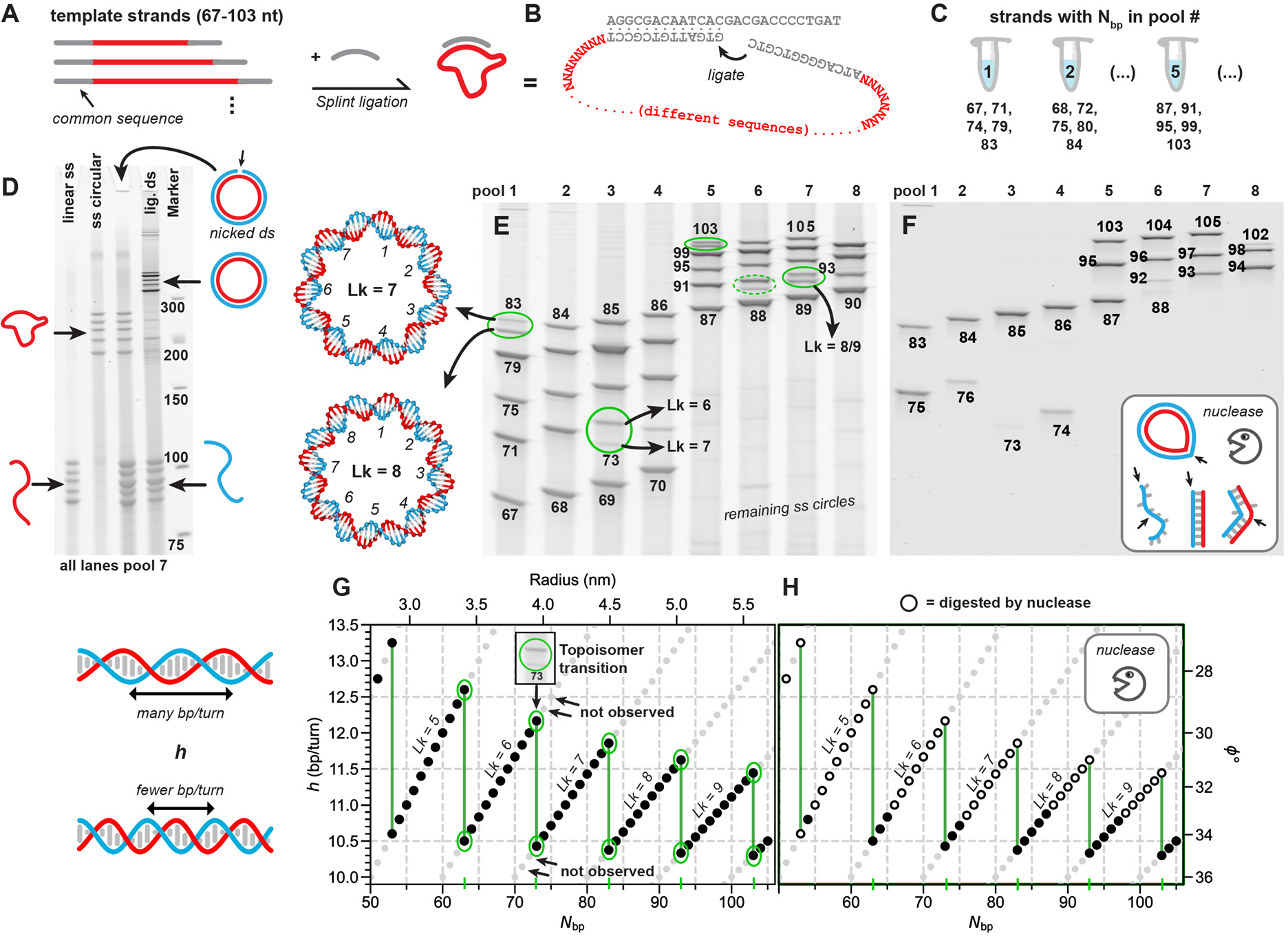
Experimental design and results of assay. (**A-B**) Template strands of different length (N_bp_ = 67-105 nt) were designed with a common terminal sequence that binds to one common splint sequence for splint ligation (**B**). The sequences between the splint binding sequences were all different sequences with varying length. (**C**) 4-5 lengths with a separation of four nucleotides were pooled together. (**D**) Denaturing PAGE analysis of different stages of the assay, shown with pool 7. Lanes from left: ss linear strands; circularized ss template; nicked ds circle separates into ss circle and ss linear strand in denaturing conditions; ligated ds circles, marker. Splints run out of the gel. (**E**) Denaturing PAGE of ligated ds circles. Green circles: Two topoisomers have different linking numbers (Lk). (**F**) Denaturing PAGE of the pools after BAL-31 nuclease digest. Only ds circles without defects resist nuclease digest. (**E/F**). (**G/H**) The helical repeat (h = N_bp_/Lk) is plotted for each observed gel band with N_bp_ after ligation (G) and nuclease digest (H). Additional gel data: **Figure S9, Figure S11**. Vertical green lines mark N_bp_ where topoisomer splitting is observed. Secondary y axis; average twist angle ϕ (ϕ = 360°/h) between adjacent base pairs (°); secondary x-axis: radius of a circle with a circumference of N_bp_ ⋅ 0.34 nm/bp.

To be able to analyze tens of lengths in one gel, we pooled 4-5 template stands with a length difference of 4 nt into one reaction (Figure 2 C; E). Figure 2D shows the different stages of the reaction with pool 7 (5 strands). Single-stranded circles formed with near-quantitative yields, as most of the complex is single-stranded and therefore very flexible (Figure 1B), entropically favoring circularization over dimerization. Although they have almost identical molecular masses, circular strands have a lower electrophoretic mobility than linear templates. The addition of counterstrands forms nicked circles but strands are separated in the denaturing PAGE gel conditions. The counterstrands (blue) are added in a 2-fold molar excess over circular templates. Ligation of nicked circles forms ligated ds circles in near-quantitative yields with a reduced electrophoretic mobility.

Figure 2E shows a gel that was run longer to achieve better separation of the ds circles and fast-migrating bands mostly ran out. Each circle with *N*_*bp*_ forms one sharp band, except for circles with *N*_*bp*_ *=*73, 83, 92-93 or 103 bp (circled in green), split into two bands. Split bands indicate the formation of two topoisomers with the same *N*_*bp*_, sequence and molecular mass, but different linking numbers *Lk*, which is the number of times one strand wraps around the other in ligated a double-stranded circle. For instance, the upper strand of *N*_*bp*_ = 83 has *Lk* = 7 and the faster migrating lower band *Lk* = 8. While circles can be clearly recognized in atomic force (AFM) images, it is difficult to identify defects (Figure S7).

The ligated circles were therefore treated with BAL-31 (Figure 2F) or S1 nuclease (Figure S8), which have both exonuclease activity and some endonuclease activity for single-stranded regions. Both nucleases therefore digest all ssDNA, dsDNA with exposed ends, and cleave ds circles at defects followed by complete digest from exposed ends. Therefore, only ds circles without internal defects survive this digest. Uncropped gel images and repeats at reduced ligase concentrations are shown in Figure S9; an experiment with an extended range from *N*_*bp*_ = 46-103 bp in Figure S11.

### Mapping Elastic Limits in *h*-Plot

Next, we mapped the gel data for all *N*_*bp*_ on an “*h* plot” (Figure 2H-I). The linking number *Lk* of any ds circle can only have integer values and cannot be changed after ligation without cutting one of the strands. Therefore, the average helical repeat *h* (base pairs per turn; averaged over the entire circle) is also fixed at *h* = *N*_*bp*_/*Lk*. For any *Lk, h* increases linearly with *N*_*bp*_ with a slope of 1/*Lk*. Only a subset of combinations of *N*_*bp*_ and *Lk* is observed in the gels (black dots). Note that most circles form only one topoisomer. For instance, ligation of nicked *N*_*bp*_ = 80 *bp* circles, only yields the *Lk* = 7 isomer (*h* = 11.4 *bp*/*turn*) but not *Lk* = 8 (*h* = 10 *bp*/*turn*) or *Lk* = 6 (*h* = 13.3 *bp*/*turn*) isomers. A higher value of *h* means that more bp are needed to complete a full helical turn and therefore DNA is underwound. While *Lk* cannot directly be determined from the gels, assigning other *Lk* values would shift all datapoints up or down to the next *Lk* which would lead to implausibly high or low average values of *h*.

The smallest circles had in general lower ligation yields, were kinking, and were digested more readily than larger rings due to higher bending stress. Moreover, circles, irrespective of their size, were digested when *h* > ~10.9. The smallest nuclease-stable circle was only 54 bp (*h* = 10.8 bp/turn) and had a radius of curvature of only 2.9 nm and is to our knowledge the smallest stable ds circle made so far.

This data suggests a simple rule to predict the stability of DNA with a mixed sequence: DNA is stable if the local radius of curvature *r* > ≈ 3 *nm* and 10.1 *bp*/*turn* < *h* < 10.9 *bp*/*turn* (see Figure 6 for overwound circles). This rule is consistent with findings from other minicircle assays and longer DNA plasmids ^28^, where the local curvature was however not controlled, and OxDNA simulations (Figure S10) showed the qualitatively the same trend.

### Determining *h*_0_

The second goal of this work was to determine if DNA unwinds due to twist-bend coupling or if *h*_0_ is constant. Unwinding can be determined from the gel data in two ways: either from the location of ligation yield maxima, or the location of topoisomer splitting (see below). Two ends can only be ligated in a stacked conformation when they are close enough for covalent bond formation. The ends in a classical looping experiment (Figure S1) or in our nicked circles (Figure 1H) have the highest stacking and ligation probability when they are torsionally relaxed (see Figure 3 A-B) at *N*_*bp*_ = *nh*_0_ and have minimal ligation probability at 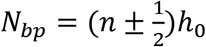 where the most torsional energy has to be overcome to bring ends together. Therefore, we originally expected to observe band intensities with a periodic trend as in Figure 1B. However, ligation was very efficient at all *N*_*bp*_ as the ends cannot dissociate and the absolute probability for ligatable configurations is orders of magnitudes higher than in looping experiments.

**Figure 3.**
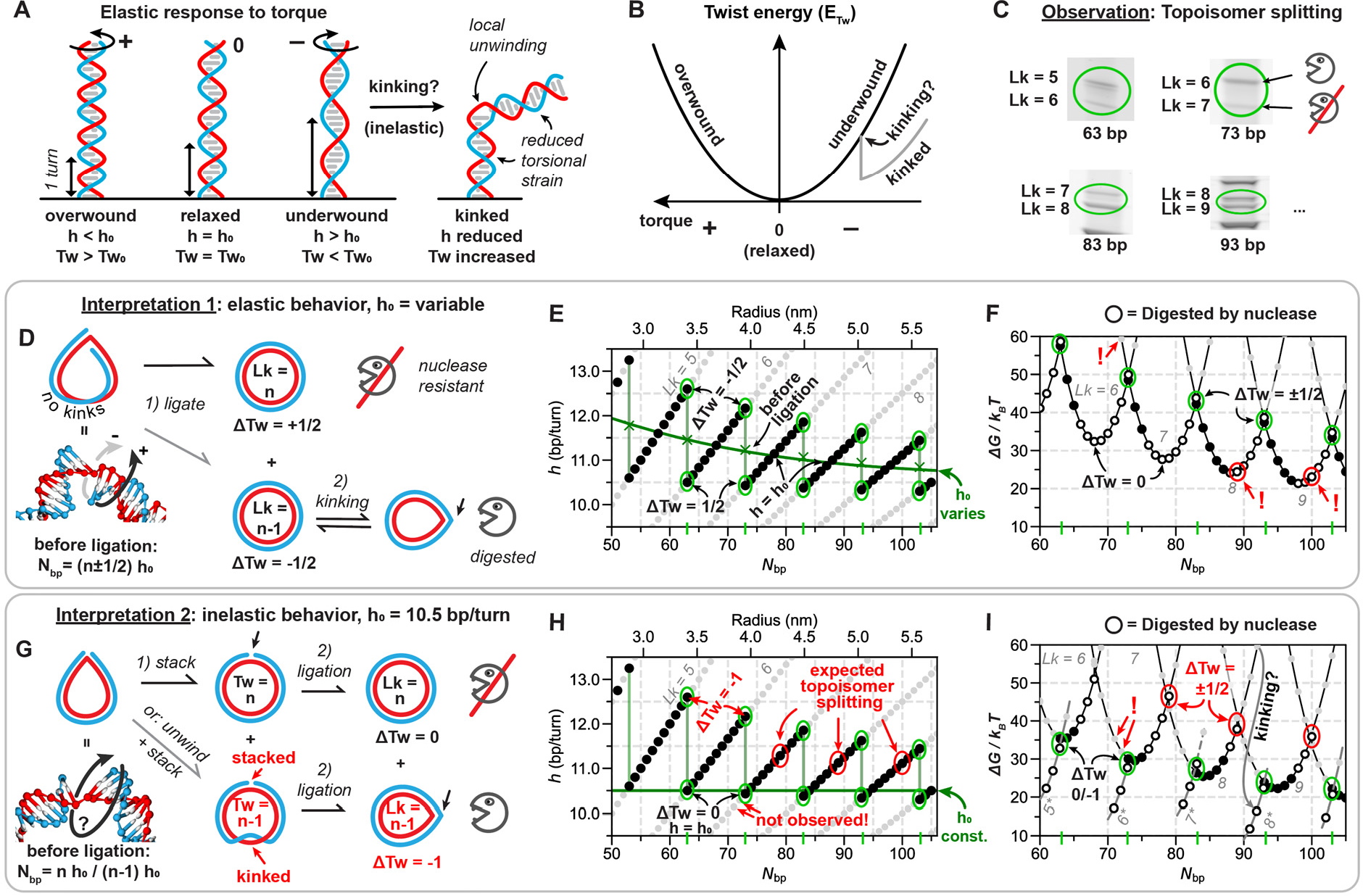
Is h_0_ constant? (**A**) Elastic and inelastic response of DNA to torsional strain, shown for linear DNA. (**B**) Torsional energy (=Twist energy E_Tw_) in a DNA molecule. (**C**)After ligation, two topoisomers appear at N_bp_ = 63, 73, 83, 93 and 103 bp and are used to find h_0_ in interpretation 1. (D, G) Schematic drawing of two interpretations for topoisomer splitting. (D-F) Interpretation 1: DNA remains elastic without kinking and h_0_ is variable. In this assumption, nicked circles are kinked and twisted before ligation (D, left). In Interpretation 2 (**G-I**) proposed by Du et al. ^37^, circles are torsionally relaxed at N_bp_ = nh_0_ before ligation. Kinking, substantial unwinding and stacking of the nick must occur before ligation for the ΔTw = −1 isomer. (**E/H**) h-plot of observed circles before nuclease digest. Note, that h-plots are identical for both interpretations, but the torsional state is interpreted differently. Green fit/line: h_0_. (**F/I**) Plot of the Gibbs free energy (ΔG = E_Tw_ E_bend_ − TΔS; see Supplementary Note 3). Assuming that h_0_ is variable (**F**) or fixed at 10.5 bp/turn (**I**), local E_Tw_ minima for the different topoisomers are at different N_bp_. In interpretation 2, kinking (*) must reduce the free energy by tens of k_B_T. Highlighted red: Implausible datapoints and states that speak against interpretation 2: Red circles at N_bp_ = 79, 89, 100: Topoisomer splitting should have been observed if h_0_ was 10.5 bp/turn without kinking. Green circles: actual topoisomer splitting (same for both interpretations).

For a constant *h*_0_ = 10.5 *bp*/*turn*, ligation yield maxima should have been observed around 73.5 bp (*Lk* = 7), 84 bp (*Lk* = 8) and so on. However, maxima shifted to longer lengths (~78-81 and ~87-90), which is best visible in experiments with a reduced ligase concentration (Figure S9). While this already indicates a significant unwinding, *h*_0_ could not be accurately determined from gel band intensities. Alternatively, *h*_0_ can be more accurately determined from those *N*_*bp*_ where topoisomer splitting is observed (Figure 3C). The band splitting can, however, be interpreted in two different ways: 1) *h*_0_ changes with curvature (Figure 3D-F) or 2) *h*_0_ is constant (Figure 3G-I).

### Interpretation 1: *h*_0_ Changes

Nicked minicircles or DNA in a looping assay can contain torsional strain in addition to bending strain. For interpretation 1, we assume DNA to be a harmonically elastic torsional spring, behaving like a twistable rubber tube as in the TWLC model (Figure 3A-B, additional discussion in Supplementary Note 3). Without torque, the DNA is torsionally relaxed, *h* = *h*_0_, and *Tw* = *Tw*_0_. Applying positive torque overwinds DNA so that *h* < *h*_0_ and *Tw* > *Tw*_0_, whereas negative torque unwinds DNA so that *h* > *h*_0_ and *Tw* < *Tw*_0_. If a kink, in which hydrogen bonds remain intact, occurs in double-stranded DNA with a fixed twist, the DNA will locally unwind at the kink due to the helical nature and asymmetry of grooves ^3,37^. As a result, the torsional stress in the remaining double-stranded region is reduced (Figure 3A-B).

If a nicked circle is in a teardrop-shaped kinked + twisted conformation without torque (Figure 3D), the two ends cannot be stacked and ligated through bending alone but must also be over- or underwound to reach integer multiples (*n*) of *Tw*. Thermal fluctuations cause rapid fluctuations of *Tw* around *Tw*_0_ and eventually, a ligatable, stacked configuration can form. If a circle is ligated, any torsional stress *E*_*Tw*_ is locked in and can only be released by cutting one strand or through inelastic kinking or melting of base pairs.

Ligation of nicked circles at most *N*_*bp*_ form only one topoisomer with a single *Lk* (Figure 3E-F), as any other topoisomer would have a higher torsional stress and would therefore have a lower probability to form in thermal equilibrium. If, before ligation, 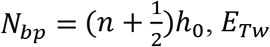 for over- or underwinding is the same and two topoisomers are formed with roughly equal probabilities, and therefore *h*_0_ = *N*_*bp*_*/Tw*. For example, the nicked circle at *N*_*bp*_ = 83 *bp* forms both *Lk* = 7 and *Lk* = 8 topoisomers. Before ligation, *Tw* was therefore 7.5 turns and *h*_0_ was 83 bp / 7.5 turns = 11.07 bp/turn, which is significantly underwound compared to consensus value of ~10.5 bp/turn. *h*_0_ was calculated the same way for all other split bands (Figure 3E green crosses), showing and increasing unwinding of DNA as circles get smaller (green fit).

In Figure 3F (and I), we plot the Gibbs free energy *ΔG* (see Supplementary Note 3) with different *N*_*bp*_, which lie on parabolas representing the different linking numbers. At the local minima, ligated circles have only bending strain, but no torsional stain as *ΔTw* = *Tw* – *Tw*_0_ = 0 and *h* = *h*_0_. The asymmetric response to nuclease digest can be explained by the local unwinding at kinks ^3,37^(Figure 3A-B): Kinking in overwound DNA is energetically costly but energetically favorable in underwound DNA. A similar behavior is known in supercoiled DNA plasmids ^28^. Torsionally relaxed circles around 68 and 78 bp contain so much bending stress that they kink in the absence of stabilizing positive torque.

Increasing *N*_*bp*_ from a local minimum at a given *Lk* increases *E*_*Tw*_ to a point where circles are so underwound that it is energetically equally or more favorable to form a circle with *Lk* + 1, which is overwound. In thermal equilibrium, the probability *P* of observing a state with energy *E* is

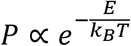

Two topoisomers are formed when their free energies are similar, or when 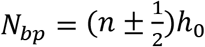 by unwinding or overwinding and where the energies of two topoisomers are roughly the same and where two parabolas intersect (*N*_*bp*_ *=*63, 73, …). The relative intensities of two bands depend on whichever *N*_*bp*_ is closer to (*n* ± 1/2)*h*_0_, but accounting for this would only insignificantly shift parabolas sideways.

While interpretation 1 is compatible with the classic harmonically elastic behavior for all lengths before ligation, it contradicts the common assumption that *h*_0_ ≈ 10.5 bp/turn and independent of local curvature.

### Alternative Interpretation 2: *h*_0_ is Constant

In a variation of the classical looping experiments, Du *et al*. devised a cyclic ligation assay with long sticky ends ^37^ to overcome low cyclization yields and also produced a small subset of small circles down to 63 bp. After ligation of fragments, they observed topoisomer splitting and nuclease sensitivity consistent with the results from our nicked minicircle assay (Figure 3C). They, however interpreted the data applying the established assumption that *h*_0_ was constant at ~10.5 bp/turn and independent of curvature. In the following paragraphs we argue that this interpretation is not plausible.

The gel data (Figure 3 C) and the *h*-plots (Figure 3 E/H) are identical for both interpretations, but the torsional stress (*ΔTw*) in the observed circles would be different and as a result, some of the data points become difficult to explain with a constant *h*_0_ (Figure 3 F, H-I: highlights in red). For instance, the digested upper band of a topoisomer splitting would be considered highly unwound to *ΔTw*~ − 1 (instead of −0.5 in our interpretation) and the nuclease-stable isomer to be relaxed at *ΔTw*~0 (instead of +0.5). Therefore, most of the observed smaller circles would be underwound up to *ΔTw*~ − 1 while overwound (*ΔTw* > 0) circles are not formed. For example, the 73 bp circle would be considered relaxed at *Lk* = 7, but the 72 and 71 bp circles do not form *Lk* = 7 topoisomers although the additional torsional and bending stress should be minimal (highlighted in Figure 3I). By contrast, in interpretation 1, the *Lk* = 7 isomers of *N*_*bp*_ < 73 do not form as their free energy is much higher than the *Lk* = 6 isomers (highlighted in Figure 3H-I).

Interpretation 2 therefore requires DNA to be extremely hard to overwind but extremely soft to unwind due to kinking, but no such asymmetry is observed in circularization assays of larger circles. While Du ^37^ acknowledges that for long circles with two isomers (205 bp), the conventional interpretation of *ΔTw* = ±0.5 must be applied and that longer circles do not kink, it is unclear why overwound topoisomers start to disappear at shorter lengths and instead highly underwound circles up to *ΔTw* = −1 start to form. If, on the other hand, DNA remained a harmonic torsional spring and *h*_0_=10.5 bp/turn was constant, topoisomer splitting should have occurred around ~69, 79, 89 and 100 bp (Figure 3H-I, red circles).

To form the unwound topoisomer (*ΔTw*~ − 1) by ligation, the nicked circle would require a stacked nick as well as a long, melted region or multiple internal kinks, as one kink can only accommodate up to *Tw*~ − 0.25 ^3^. Moreover, as the two bands form in the same experiment, the two states would have to have the same energy. Therefore, kinking would have to provide a large amount of unwinding and/or high energetic stabilization compared to an elastic behavior (tens of *k*_*B*_*T*, see arrows to kinked states in Figure 3 I, and Figure S2) exceeding the total bending energy.

In summary, while interpretation 2 keeps *h*_0_ at the established textbook value of ~10.5 bp/turn, many consequences of this assumption seem implausible, but can all be resolved by interpretation 1.

A central mechanistic difference between the two interpretations is the timepoint at which kinking occurs: In interpretation 1 nicked circles before ligation only kink at the nick; kinks form after ligation. In interpretation 2, a highly underwound nicked circle with *ΔTw*~ − 1 with one large defect or multiple small defects must be formed before ligation of the nick, but the nicked must still be stacked for ligation to occur. To test when kinking occurs and which interpretation is correct, we performed all-atom MD simulations as well as additional experiments.

### Excess Twist in MD Simulations

We set up two atomistic simulations of nicked circles with 84 bp with the OL21 force field, in which *h*_0_ ≈ 10.5 *bp*/*turn* for long, mixed sequences in a stacked starting configuration. One nicked circle was underwound with *Tw* = 7 (*ΔTw*~ − 1; Figure 4A-E), which is the configuration that needs to form in thermal equilibrium for interpretation 2 (Figure 3G bottom) before ligation. For comparison, we simulated another nicked circle under positive strain with *Tw*~9 (*ΔTw*~1, Figure 4F). During a 50 ns simulation, we tracked *h* (Figure 4B) as well as broken hydrogen bonds and stacking between neighboring base pairs in the DNA (Figure 4D, see methods for details). The plots in Figure 4E show the total number of broken interactions during the simulation.

**Figure 4.**
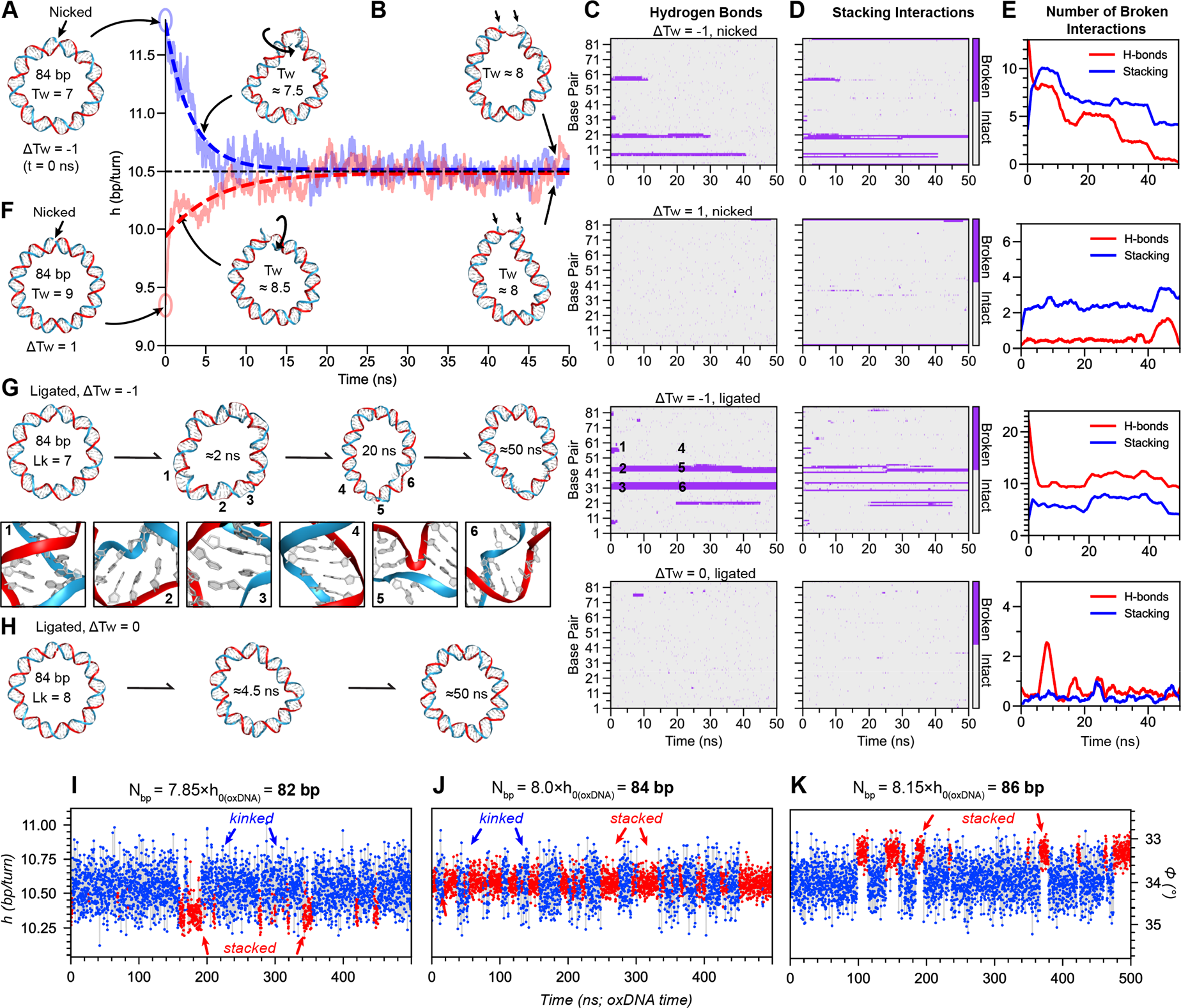
**MD simulations of different circles with** N_bp_ = 84. (A-H) atomistic and I-K oxDNA simulations. (**A-E**) nicked, Tw = 7 (ΔTw~ − 1); (**F**) nicked, Tw = 9 (ΔTw~ 1); (G) ligated ΔTw = −1; (H) ligated ΔTw = 0. (**A, F, G, H**) Starting configurations at t = 0 ns; other snapshots at indicated times. (**1-6**) Enlarged segments at 2 and 20 ns. Full simulations are shown in Supplementary movies 4-8. (**B**) Analysis of h averaged out over the nicked circles from 0-50 ns. Analysis of intact or broken (purple) hydrogen bonds (**C)** or stacking interactions (**D**). The base pairs numbers are counted anticlockwise from the nick/top. (**E**): Total number of broken H-bonds or stacking interactions of the different systems during the simulation. (**I-K**) Plots of oxDNA simulations showing the dynamic transitions between stacked states (red points) and kinked states (blue) from 500 ns simulations of different circles with 82 (**I**), 84 (**J**) and 86 bp (**K**). h (scale on left) was calculated from the average twist angle ϕ between adjacent base pairs (scale on right; h = 360°⁄ϕ). Note that kinked conformations fluctuate around the equilibrium h_0_ of oxDNA (10.55 bp/turn), while stacked conformations can be overwound (**I**) or underwound (**K**).

In both simulations, the nicks kink within the first nanosecond of the simulation and the torsional stress in both simulations is completely resolved by rotation around the nick within only ~20 ns, after which *h* fluctuates around *h*_0_ = ~10.5. The unwound circle developed >15 internal defects during equilibration, but the overwound circle did not (Figure 4E). This result is consistent with the experimental observation from nuclease digests that underwound DNA kinks more readily than overwound and the local unwinding at kinks (Figure 3A). As the torsional stress is released, all broken hydrogen bonds and most of the broken stacking interactions of the underwound circle were restored while no new long-lived defects form in either nicked circle (Figure 4C-E).

Next, we simulated two ligated circles with a *ΔTw* ≈ −1 and *ΔTw* ≈ 0 (Figure 4G-H) that are postulated to form after ligation of nicked circles with *ΔTw* ≈ 0 in interpretation 2 (Figure 3G) with about the same probability / free energy. The underwound circle contains an average of 10.9 broken H-bonds and 6.3 broken stacking interactions. As breaking the hydrogen bonds of one base pair costs on average ~1 k_B_T of energy and breaking the stacking even ~1-4 k_B_T ^38^, these defects carry a high energy cost. Moreover, the defects do not release the entire torsional stress and the intact segments containing ~70 bp are still underwound to *h* ≈ 10.72 bp/turn (*ΔTw* ≈ 0.17 turns) resulting in a residual twist energy of *E*_*Tw*_ ≈ 2.4 k_B_T.

In the relaxed, ligated circle with *ΔTw* ≈ 0 on the other hand, only very brief disruptions of H-bonds and stacking interactions were observed. The energetical difference between the two circles is tens of k_B_T and forming a ligatable stacked transition state with *ΔTw* = −1 as required by interpretation 2 (Figure 3G-I) is therefore exceedingly unlikely, supporting interpretation 1 (Figure 3D-F) and a changing *h*_0_.

As transitions from kinked to a stacked configuration were rare in the time scales of atomistic simulations, we performed additional oxDNA simulations on a slightly overwound (*N*_*bp*_ = 82, Figure 4 I), relaxed (*N*_*bp*_ = 84, Figure 4 J) and slightly unwound circle (*N*_*bp*_ = 86, Figure 4 K). Transitions between stacked and kinked conformations are most frequently observed in the relaxed circle J and *h* of the stacked states differ (red datapoints in Figure 1I-K; Figure S5, Figure S6 Supplementary Movies S1-4). Note that diffusion is accelerated in oxDNA and therefore processes in an experiment or all-atom MD simulations would likely take ~1-2 orders of magnitude longer ^39,40^.

### Synthesis Errors in Oligonucleotides?

The simulations above have defect-free sequences, but experimentally, internal kinks could also form at sequence errors such as base substitutions, insertions or deletions that might be introduced during chemical oligonucleotide synthesis ^15,16^. However, we purchased oligonucleotides with the highest commercial purity grade with a per base error rate of < 0.1 %. Moreover, due to the capping step after each synthesis cycle, most remaining errors are 5’-truncations, which cannot be ligated due to the specificity of ligases around the ligation site. Most double-stranded circles should therefore be error-free. In control experiments, circles containing designed mismatches or insertions/deletions were fully digested by nucleases (Figure S12), whereas the band intensities of circles without defects remains the same after nuclease treatment (Figure S8), which proves that they are free of defects.

Alternatively, we enzymatically extended the splint with T4 DNA polymerase ^41^ to produce circles with an even lower error rate (Figure S13). Ligation of the resulting nicked circles produced topoisomer splitting at the same *N*_bp_ as in experiments with fully synthetic strands. We conclude that synthesis errors cannot explain our results and that mechanical stress must be responsible for kinking in unwound DNA.

### Kinking Before or After Ligation?

While both oxDNA and aa MD simulations suggest that nicks are the weakest link and that kinking away from the nicks are rare and only short-lived events, ligation experiments take orders of magnitudes longer than the timescales accessible by simulations. To experimentally test if kinking of underwound circles occurs before or after ligation, we created nicked circles with a total of 11 phosphorothioate (PS) bonds between nucleotides around the nick that nucleases cannot efficiently cleave. A PS-protected nicked circle with *N*_bp_ = 63 (Figure 5A1) is digested much slower than a PS-protected nicked minicircle with a mismatch (Figure 5A2) or a minicircle with regular phosphate linkages (Figure 5A2).

**Figure 5.**
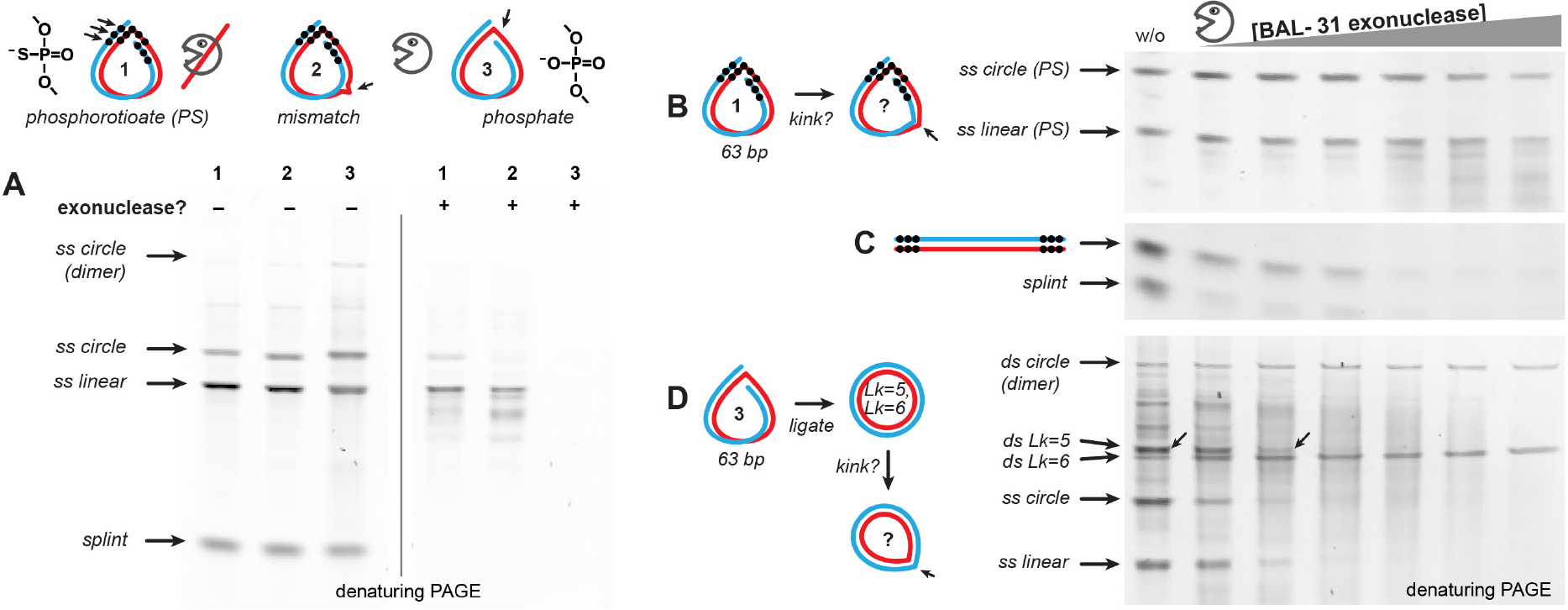
Digest assay with protected nicked minicircles. Denaturing PAGE gels of: (**A**) nicked circles (N_bp_ = 63; with a total of 11 PS bonds around the nick (**1**); PS protection and an additional mismatch (**2**), and the control nicked circles with regular phosphate bonds (**3**)) before and after digest with BAL-31 nuclease. (**B**) PS protected nicked minicircle at increasing BAL-31 concentrations. (**C**) Linear PS protected duples. (**D**) Crude ligation reaction of the nicked minicircle with regular phosphate. (**B-D**) Amounts of nuclease per reaction: 0.023; 0.032; 0.047; 0.068; 0.098 and 0.16 U.

We next performed a series of digests with increasing concentrations of BAL-31 nuclease with a nicked, PS protected circle (Figure 5B) and a linear dimer with PS protections at the ends (Figure 5C), and the crude ligation product of the nicked 63 bp circle with standard phosphates (Figure 5C). The nicked starts to show some degradation at higher nuclease concentrations, but the linear PS dimer is digested as a much faster rate (Figure 5B-C). This indicates that the BAL-31 nuclease has a residual activity despite PS-protections. The ligation of the nicked 63 bp circle forms two double-stranded topoisomers with *Lk* = 5 and 6 as seen before (Figure S11 and ref. ^37^). Before digest, the band of the *Lk* = 5 isomer is more intense but disappears even at low nuclease concentrations while the overwound *Lk* = 6 isomer and the circular dimers persist digest.

These results also demonstrate that small, nicked circles do not significantly kink before ligation, or at least much less than the ligated *ΔTw* = −1/2 isomer, supporting our interpretation 1 (Figure 3). We suggest the following mechanism for topoisomer formation and kinking: *h* fluctuates quickly in nicked circles (ps to ns timescales, see Figure 4B, I-K), but kinked and twisted minicircles will only rarely reach a ligatable *ΔTw* = ±1/2 configuration (see Figure 1H, minima). Internal kinking is rare and very short-lived in nicked circles (Figure 4B). Once nicked ends stack, the bending stress is slightly increased, and any torsional stress is locked in by ligation. After ligation, *ΔTw* = −1/2 circles develop kinks that are stable enough to be recognized and digested by nucleases while the overwound *ΔTw* = 1/2 isomers are mechanically stabilized. We therefore conclude that the observed increase of *h*_0_ in tightly bent DNA is not an effect caused by internal kinks before ligation.

### Alternative Explanations for Unwinding

Physical models often treat DNA as an elastic rod or beam where three fundamental modes of deformation exist: stretching, bending, and twisting. Twist can be influenced by either stretching or bending. Although stretching at forces below 1 pN can straighten entropically coiled DNA and allow one to determine the bending persistence length ^42,43^, twist-stretch coupling is only observed at higher forces ^44,45^ where DNA counterintuitively first slightly overwinds at small forces (< 20-30 pN) and then unwinds at higher forces (> 30 pN). The tension in minicircles of our study is, however, only ~3-5 pN (see Supplementary Figure S10 and Supplementary Note 4), and the influence of twist-stretch coupling on the measured helical repeat is therefore negligible.

Theoretically, the T4 DNA ligase might also influence the assay results. While *h*_0_ of DNA in ligase complexes is not significantly changed ^46,47^, a stabilization of internal defects by ligase was reported ^48^. On the other hand, an increased helical repeat consistent with TBC and our data also appears in ligation-free circularization experiments where *j*-factor maxima ^14^ and the uncircularization kinetics ^21^ also suggest *h*_0_ ≈ 11-11.3 bp/turn for small circles. Different *h*_0_ found in ligation-based circularization experiments ^5,8^ can be explained by sticky end geometries and kinks at nicks (ref. ^21^, Supplementary Note 1). Thus, the observed increase of *h*_0_ does not seem to be linked to ligases.

### Intercalators, Histones, and Temperature

The helical repeat of DNA can be influenced by many factors including temperature, salts, and intercalators such as ethidium bromide and DNA-binding proteins. We next determined this shift in tightly bent DNA.

For this we added Ethidium bromide (EtBr), a common intercalating dye that unwinds DNA ^30^, to ligation and digest experiments (Figure 6A, gels in Figure S14). As expected, topoisomer splitting shift right to longer *N*_*bp*_ and some circles appear at a lower *Lk* than without additives and in the h-plot data points shift up (blue arrows). In addition, EtBr stabilizes underwound circles during nuclease digest up to *h* ~12 bp/turn.

**Figure 6.**
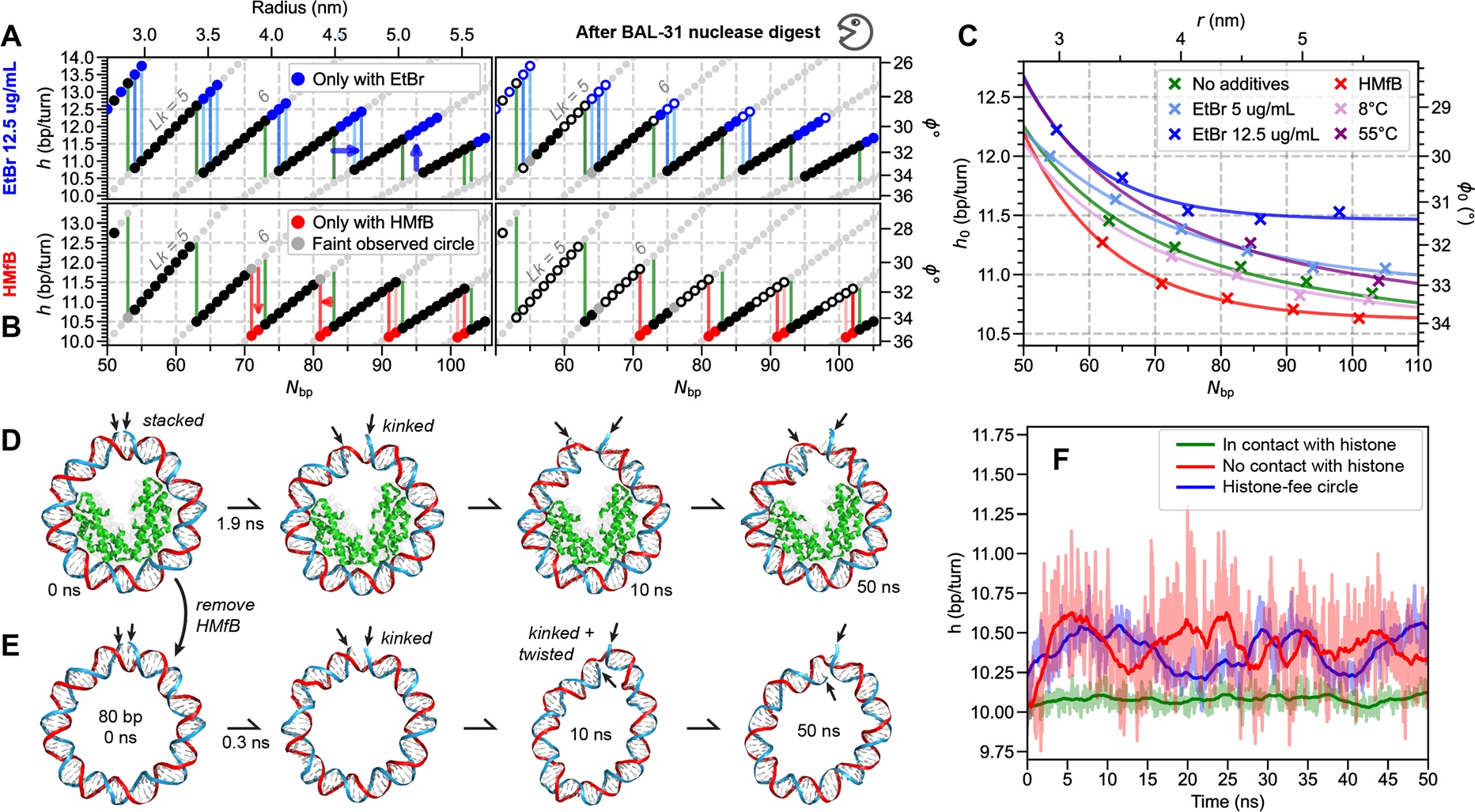
Effect of EtBr, histones and temperature. **A)** h-plot of results from ligation results of nicked circles in the presence of EtBr (gels, see Figure S14) before and after BAL-31 nuclease digest. Blue datapoints: topoisomers that are only formed with EtBr; blue vertical lines: topoisomer splitting; green vertical lines: topoisomer splitting without additives for reference (compare Figure 2). **B**) ligation in the presence of HMfB histone-like proteins (gels, see Figure S15). Red: Topoisomers that only form in presence of HMfB. Before nuclease digest (right), HMfB proteins were removed. (C) Plot of curvature-dependent h_0_ at different EtBr concentrations, HMfB and temperatures. **D**) Atomistic MD simulation snapshots of a nicked 80 bp circle containing two histone-like HMfB dimers. (**E**) MD simulation snapshot of the nicked circle as in (**D**), but without HMfB. (**F**) Analysis of h of the DNA segments that are in contact with HMfB (green); the remaining DNA of (**D**) that is not in contact with HMfB (red); and the histone-free circle (blue).

Next, we added HMfB proteins at a 6-fold molar excess to nicked minicircles and ligated them. HMfB is a small, positively charged histone-like protein that is highly conserved in archaea and has a structure that is almost identical to eukaryotic histone proteins ^49^ and positively supercoils DNA ^28^. The overwinding is also observed in our assay and topoisomer splitting occurred at circles that were 1-2 bp smaller than in the control (Figure S15). This shifts datapoints in the h-plot down, and topoisomer splitting left (Figure 6B, red).

After ligation, proteins were removed by protease, and the remaining dsDNA circles were treated with nuclease. The newly formed overwound topoisomers had the lowest *h* that we produced in this study and were nuclease stable (Figure 6A-B, red datapoints) while those circles with *h* > ~10.9 were still digested, consistent with a mechanical stabilization of overwound DNA against kinking. No new circles were observed with *Lk* = 6, presumably due to insufficient space for HMfB dimers in the center of the circles or a reduced binding affinity at overly tight curvatures.

Adding a high concentration of potassium salt that is present in the nucleus only slightly increases *h*_0_ (Figure S17) while lowering the ligase concentration 10- or 100-fold did not change topoisomer splitting (Figure S9). Increasing the temperature unwinds linear DNA ^50^ and is also seen in oxDNA simulations and ligation experiments of nicked minicircles (Figure S16). Figure 6C summarizes the curvature-dependent *h*_0_ for the different conditions.

The overwinding due to HMfB could be reproduced in an atomistic MD simulation of a nicked circle containing two HMfB dimers, which we generated by modifying the crystal structure of an HMfB-DNA complex (PDB:5T5K, ref. ^49^, see methods). An initially stacked circle kinked quickly, but the HMfB proteins remained stably bound to the DNA (Figure 6D). The identical circle without proteins were relaxed quickly into a kinked and twisted configuration (Figure 6E) indicating that the HMfB proteins overwound the DNA. This is also reflected in the graph in Figure 6E, where the segment of DNA that is in contact with Histone proteins is overwound to ~10.1 bp/turn (green), whereas the protein-free segments (red) and protein-free circle (blue) fluctuate around *h*_0_ ≈ 10.45 *bp*/*turn*.

### Conclusions and Biological Implications

The genetic information of organisms is best protected in dsDNA while kinking increases the probability for physically, chemically or biologically damaging DNA. After decades of research, the elastic limits of DNA are still hotly debated ^3–22^. Most studies focused either on bending or torsional stress. Due to the inability of established assays to simultaneously control *r* and *h*, it has not been possible to map which combinations of bending and torsional stress kink DNA. The nicked minicircle assay, that we describe here, allows us to systematically test the elastic limits in the bend/twist parameter space. By changing circle circumferences in 1-bp steps, we could control the average radius of curvature *r* in 0.05 nm increments down to *r* ≈ 2.7 *nm* and the twist from *h* ≈ 10 – 13 bp/turn. We found that DNA is mechanically stable if *r* ≥ ~3 *nm* and *h* ≤ ~10.9 bp/turn even in the absence of stabilizing proteins. Underwound DNA with *h* > ~10.9 bp/turn kinks even at lower radii of curvature. Qualitatively, the nuclease sensitivity of unwound DNA was also observed in plasmid DNA ^28^, but due to writhing and plectoneme formation, the local curvature and twist vary in such large templates while circles *N*_*bp*_ < 174 do not writhe ^51^.

A second important finding was that the natural helical repeat *h*_0_ of DNA is not constant at ~10.5 bp/turn but increases to over 11 bp/turn at biologically relevant curvatures. Unwinding of tightly bent DNA as a result of twist-bend coupling was theoretically predicted three decades ago by Marko and Siggia ^36^ but had not been experimentally quantified, yet.

The length-dependent, or rather curvature-dependent change of *h*_0_ and the magnitude of unwinding (~10 %), is in almost perfect agreement with the early predictions ^36^, data from classical circularization experiments (black datapoints from ref. ^30–32^ and asymptotically converges to the consensus *h*_0_ ≈ 10.45 (±0.05) *bp*/*turn* for mechanically relaxed DNA (additional data with long circles, see Figure S18).

However, according to more recent estimates for the magnitude of TBC ^52–55^, *h*_0_ should only widen by ~ 0.2 % at a comparable curvature of nucleosomal DNA, which would have been practically indetectable in our assay (Figure 7A, red curve; see Supplementary Note 2). While the original MS equation (Supplementary Note 2, eq. 19) fits excellently to our experimental data in Figure 3B, the high value of the fitted TBC parameters *G*/*C* = 2.7 may indicate that additional physics is at play. General stability requirements of the MS model are difficult to satisfy for a linear model when *G*⁄*C* > 1, and non-linearities might have to be introduced (Supplementary Note 2). In contrast, the TBC parameter estimates from ref. ^52–54^ satisfy *G*/*C* < 1 naturally, but the effect on the helical repeat is irreconcilably weaker than our experimental findings.

**Figure 7.**
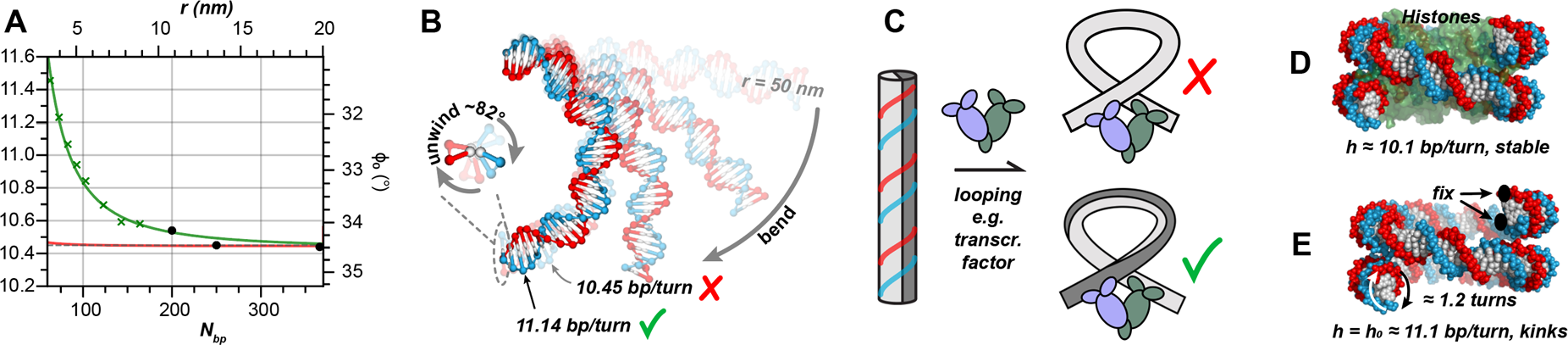
Conclusion and biological implications. **A**) Relaxed helical repeat (h = h_0_) of a duplex bent with a constant radius of curvature r (top axis) matching that of a minicircle of N_bp_ base pairs (bottom axis). Green crosses: h_0_ determined from topoisomer splitting at N_bp_ = (n 1/2)h_0_. **Black dots**: h_0_ data points from classical circularization experiments (5–7). **Red:** Prediction of the change of h_0_ due to TBC with parameters from ^52,53^. (**B**) Illustration of a 39 bp DNA fragment with decreasing radii of curvature (r = 50, 12, 6.4 or 4.2). Transparent overlay: h_0_ = 10.45 bp/turn; the unwinding for a semicircle due to TBC is about 82°. (**C**) Looping of DNA by transcription factors or DNA-binding proteins. Considering unwinding, the relative orientation of required binding sites to avoid torsional stress changes. (**D**) DNA in a nucleosome (PDBID: 3LZ0) has an average helical repeat of ~10.1 bp/turn ^35^. (**E**) If one could remove the histone proteins while maintaining the overall curvature of the nucleosomal DNA (r~4.5 nm), it would be torsionally relaxed at ~11.1 bp/turn but would be more susceptive to kinking; it follows that ΔTw 1.2 turns.

The unwinding due to twist-bend coupling plays a role wherever DNA is tightly bent or twisted in biology, and the associated mechanical stress influences the energetics of all molecular processes involving DNA. For example, transcription factors can loop DNA into small radii of curvature and DNA should unwind it. Consistent with our findings, an increased helical repeat was reported for various transcription factors including the *lac* repressor ^56^, araCBAD ^57^ or in HU protein induced loops ^58^. The unwinding will significantly change the relative geometry of binding sites for DNA-binding proteins including transcription factors or DNA-modifying enzymes (Figure 7C). Considering unwinding, binding sites that would be on the same side of straight DNA could face one another (dark half cylinders touch). Similarly, this changing orientation could either expose or conceal binding sites for other proteins depending on the local configuration of the DNA and molecular complexes.

In eukaryotes, ~72% of the total DNA in the nucleus is nucleosomal DNA with a radius of curvature of ~4.5 nm (Figure 7D). Our data suggests that protein-free DNA with this curvature would be torsionally relaxed at *h*_0_ ≈ 11.1 bp/turn (Figure 7E), but instead, nucleosomal DNA is overwound to *h* ≈ 10.1 bp/turn ^35^. Over the length of 147 bp, this results in a *ΔTw* ≈ 1.2 and *E*_*Tw*_ ≈ 60 *k*_*B*_*T* ≈ 35.4 *kcal*/*mol* (Supplementary Note 3, eq. 28) in addition to a bending stress *E*_*bend*_ ≈ 60 *k*_*B*_*T* (eq. 26). The total mechanical stress of the DNA in a nucleosome would therefore be *E*_*mech*_= *E*_*Tw*_ + *E*_*bend*_ ≈ 120 *k*_*B*_*T*, which must be compensated through interactions between the DNA and histone proteins, and must affect wrapping and unwrapping and therefore gene regulation and expression ^59^. Our findings of the torque-dependent elastic limits also suggest that histones have evolved to overwind DNA to oppose the intrinsic unwinding due to twist-bend coupling, providing mechanical protection against kinking.

Tight bends are also observed in virus capsids or bacterial genomes ^60^, which would also change the relative orientation of protein binding sites or sequence motifs. Both curvature-dependence of the helical repeat and torque-dependent elastic limits will also have to be considered in the design of DNA nanostructures that contain small minicircles such as DNA catenanes ^61^, rotaxanes ^62^, DNA-encircled lipid nanodiscs ^63^, tightly bent DNA origami structures ^64,65^ or in the template length-dependent amplification bias during rolling circle amplification ^66^. Finally, a consequence of TBC is that torque from helicases, polymerases or DNA packing motors will induce bending. We conclude that torque-dependent elastic limits and the increased helical repeat caused by TBC must be considered in all processes where DNA is tightly bent or twisted.

## Data and materials availability

Raw experimental or simulation data as well as code is available upon request.

## Supplementary Materials

Supplementary Text

Figs. S1 to S18

Tables S1-S2

Movies S1 to S14

Supplementary Data are available at NAR online.

## Acknowledgments

We thank Dr. Ned Seeman, Dr. Michael Matthies, Dr. Petr Sulc, Dr. Christopher Maffeo, Dr. Aleksei Aksimentiev, Dr. Hamza Balci and Dr. Alexander Vologodskii for helpful discussions.

## Author contributions

T.L.S. and J.P. conceived the study; S.C., R.B, M.M, M.S. and D.H. performed experiments; D.R.H., T.S. and R.B. performed oxDNA simulations and analysis; F.F. established the all atom MD simulation protocols; E.M., R.B. and F.F. performed and analyzed all-atom MD simulations; all authors analyzed and visualized the data; T.L.S. acquired funding; T.L.S. and J.P. supervised the study; T.L.S. wrote the initial draft; all authors revised and expanded the manuscript.

## Materials and Methods

### Materials

All long (45-105 nt) DNA oligonucleotides were purchased from Eurofins Genomics (EXTREmer purity grade) suspended in TE buffer (10 mM TRIS, 0.1 mM EDTA, pH 8.0) to 10 µM. The splint oligonucleotide (26 bases) was purchased from Integrated DNA Technologies (IDT) in lyophilized form and resuspended in TE buffer to 100 µM. GeneRuler Ultra Low Range DNA ladder was purchased from Thermo Fisher Scientific (cat. no. SM1211). All enzymes [T4 Polynucleotide Kinase (cat. no. M0201L), T4 DNA ligase (cat. no. M0202L), Hi-T4 DNA ligase (cat. no. M2622S) and Salt-T4 DNA ligase (cat. no. M0467S)] were purchased from New England Biolabs (NEB). BAL-31 nuclease (cat. no. 2510A) was purchased from Takara Bio and S1 nuclease (cat. no. EN0321) was purchased from Thermo Scientific. All steps (phosphorylation, annealing, ligation) to form the ligated double-stranded minicircles were performed in ligase buffer (50 mM TRIS-HCl, 10 mM MgCl_2_, 1 mM ATP, 10 mM DTT, pH 7.5) purchased from New England Biolabs (cat. no. B0202S). Tris(hydroxymethyl)aminomethane was purchased from Sigma Aldrich (cat. no. 252859), Boric acid (cat. no. BDH9222) and disodium salt of EDTA were purchased from VWR (cat. no. BDH4616). EtBr (cat. no. 15585011) was purchased from Thermo Scientific. HMfB (cat. no. 10948P) was purchased from Beta Lifescience. 15% TBE-Urea gels were purchased from Thermo Fisher Scientific (cat. no. EC68855BOX). All gels were run in 1X TBE (100 mM TRIS, 100 mM Boric acid, 2 mM EDTA, pH 8.3). SYBR safe DNA gel stain was purchased from Thermo Fisher Scientific (cat. no. S33102).

### Experimental Methods

Oligonucleotides of 67-105 bases (total of 39 different oligonucleotides) were pooled together in 8 different pools such that each pool contained 4-5 oligonucleotides with *ΔN*_*bp*_ = 4 *nt* as indicated in Fig. 2G.

#### Phosphorylation of Template Strands

Each pool containing a total amount of 500 pmol of oligonucleotides (100 pmol of each of the 5 template oligonucleotides of 10 µM starting concentration) was phosphorylated in separate PCR tubes. For this, the oligonucleotide stocks were mixed with 50 U (5 µL of a 10 units/µL stock; 10 units of kinase per 100 pmol of template oligo) of T4 Polynucleotide Kinase and 10X ligase buffer to a final concentration of 1.66 µM per oligonucleotide to a final volume of 60 µL in 1X ligase buffer. The reaction was incubated at 37 °C for 30 min followed by heat inactivation of the enzyme at 65 °C for 20 min and kept at room temperature until further processing.

#### Splint Annealing

The phosphorylated oligonucleotide pools (60 µL) were mixed with 25 µL of a 5-fold molar excess of the splint oligonucleotide (100 µM stock concentration; 500 pmol of splint for every 100 pmol of phosphorylated oligo) in 1X ligase buffer to obtain a final concentration of 1.14 µM of each phosphorylated oligonucleotide and a total reaction volume of 89 µL. The splint was annealed by heating to 80 °C for 5 min followed by cooling to 50 °C at a rate of −1 °C per minute and from 50 °C to 25 °C at a rate of −5 °C per minute. For minicircles shorter than ~70 bp, different lengths of splints were tested (data not shown). Our 26 nt standard splint showed a better circularization efficiency than shorter or asymmetric splints. However, the annealing was modified such that the heated samples were quickly cooled in an ice bath or with fast cooling to 4 °C by the PCR cycler to favor monomeric circles over dimerization.

#### Ligation

2000 U of T4 DNA ligase (5 µL of a 400 units/µL stock; 400 units per 100 pmol of template oligo) were added to the annealed oligonucleotides (89 µL) such that the final concentration of each oligonucleotide was now 1.07 µM and total volume was 94 µL. The reaction was incubated at 25 °C for 60 min and at 30 °C for 60 min. The ligase was heat inactivated at 65 °C for 10 min. After this, circularized single-stranded template oligonucleotides were aliquoted and frozen for subsequent nicked and ligated double-stranded minicircle syntheses.

#### Phosphorylation of Complementary Strands

Each pool containing a total of 1 nmol of complementary oligonucleotides (200 pmol of each of the 5 complementary oligonucleotides of 10 µM starting concentration; 2-fold molar excess of complementary strands to circular template strands) was phosphorylated using 100 U (10 µL of a 10 units/µL stock) of T4 polynucleotide kinase in 1X ligase buffer in a total volume of 250 µL such that the final concentration of each oligonucleotide was 0.8 µM. 10 U of kinase were used per 100 pmol of each oligonucleotide at 37 °C for 30 min followed by heat inactivation of the enzyme at 65 °C for 20 min.

#### Formation of Nicked Minicircles

Phosphorylated, linear complementary strands (250 µL) were added at a 2-fold molar excess to ligated single-stranded minicircles (94 µL) and 344 µL of this mixture was annealed by heating to 80 °C for 5 min followed by cooling to 50 °C at a rate of −1 °C per minute and from 50 °C to 25 °C at a rate of −5 °C per minute. The final concentration of the circular template oligonucleotides was 0.30 µM (each).

#### Ligation of the Double-Stranded Minicircle

167 µL of nicked double-stranded minicircles (consisting of 50 pmol of each of the 5 template strands, 0.30 µM) was incubated in a thermocycler at the desired ligation temperature for about 15 minutes. Then, 1000 U (2.5 µL of a 400 U/µL stock) of T4 DNA ligase were added to each oligonucleotide pool (total volume = 169.5 µL) and incubated for 30 min. 400 units of ligase were used per 100 pmol of the oligonucleotide. The enzyme was not heat inactivated to prevent changes in the helical repeat by heating and heat-induced enhancement of ligase activity. Hence, the enzyme was chemically inactivated by adding 19 µL of a 300 mM EDTA stock to a final concentration of 30 mM in all pools.

#### Variation of [Ligase]

All steps up to formation of nicked minicircles were performed as described above. Each pool of the nicked double-stranded minicircles (167 µL) was incubated at 37 °C. Meanwhile, T4 DNA ligase was diluted 100-fold in 1X ligase buffer to make a 0.01X stock of the enzyme. After about 15 minutes, 10 U (2.5 µL of a 4U/µL stock) of diluted T4 DNA ligase were added to all pools and incubated for 30 min. 4 units of ligase (0.01X) were used to ligate 100 pmol of each oligonucleotide. After 30 min, 19 µL of a 300 mM stock of EDTA was added to deactivate the enzyme and the reactions were analyzed by 15% TBE-urea gels.

#### Variation of Temperature

All steps up to the formation of nicked circles were performed as described above. Four sets of nicked double-stranded minicircles (each set containing 167 µL per pool) were incubated at 8 °C, 25 °C, 37 °C or 55 °C for 15 minutes before 1000 U (2.5 µL of a 400 U/µL stock) of T4 DNA ligase were added to all 8 oligonucleotide pools simultaneously in a total reaction volume of 169.5 µL. For samples at 55 °C, an engineered heat-resistant version of the wild-type T4 ligase called the Hi-T4 ligase was used because the wild-type is rapidly heat inactivated by high temperatures. For the other temperatures, 400 U of regular T4/Hi-T4 ligase (1X) were used to ligate the pools containing 100 pmol of each oligonucleotide. Samples at 8 °C were incubated for 12 h whilst samples at 25 °C, 37 °C and 55 °C were incubated for 30 min, 30 min and 20 min respectively. Enzymes were inactivated with 19 µL of a 300 mM stock of EDTA to obtain a final concentration of 30 mM and pools were analyzed by 15% TBE-Urea gels.

#### Variation of [Salt]

To nicked, double-stranded minicircles (167 µL per pool), 21 µL each of 2.66 M KCl and 100 mM NaCl were added in addition to the Mg^2+^ present from the ligase buffer (necessary for ligase activity) to obtain a final volume of 210 µL and a final concentration of 266 mM KCl and 10 mM NaCl in solution. Samples were incubated at 37 °C for 30 min. For this experiment, an engineered salt-tolerant version of the wild type T4 DNA ligase called Salt-T4 DNA ligase was used because regular T4 ligase is less active at high monovalent ion concentrations. 1000 U (2.5 µL of 400 U/µL stock) of salt-T4 DNA ligase were added per pool. In other words, 400 U of the salt-T4 DNA ligase (1X) were used per 100 pmol (total). 30 min after enzyme addition, 23.5 µL of a 300 mM stock of EDTA were added to a final EDTA concentration of 30 mM for enzyme inactivation. Pools were analyzed by 15% TBE-Urea gels.

#### Variation of [EtBr]

To nicked, double-stranded minicircles (20 µL per pool, 10 pmol), 2.4 µL of a 50 µg/mL stock of EtBr (in water) were added to a final concentration of 5 µg/mL. Samples were incubated at 25 °C for 60 min. 400 U (1 µL of 400 U/µL stock) of T4 DNA ligase were added per pool. Samples were incubated at 37 °C for 30 min. 30 min after enzyme addition, 2.5 µL of 300 mM EDTA were added to a final EDTA concentration of 30 mM for enzyme inactivation. Pools were analyzed by 15% TBE-Urea gels. Similarly, to 20 µL per pool (10 pmol each), 3 µL of a 100 µg/mL stock of EtBr (in water) were added to a final concentration of 12.5 µg/mL.

#### Addition of HMfB and Ligation

To nicked, double-stranded minicircles (20 µL per pool, 10 pmol each, 50 pmol total), a 6-fold molar excess (300 pmol) of HMfB histone protein was added. For this, 3.5 µL of 85 µM HMfB solution (1 mg/mL in water) were added and incubated for 90 min at 37 °C. Then, 400 U (1 µL of 400 U/µL stock) of T4 DNA ligase were added per pool and incubated at 37 °C for 30 min. HMfB and ligase were then digested using Proteinase K using a protocol adapted from ref. ^28^. For this, 0.5 µL of 20 mg/mL stock of Proteinase K were added and incubated at 45 °C for 60 min in the presence of 1 % SDS. The denatured proteins were purified/ buffer-exchanged by affinity column purification (DNA clean, Zymo research). Caution needs to be exercised as affinity columns tolerate only up to 0.1 % SDS. Hence, the sample must be sufficiently diluted before loading onto the column as SDS in the sample will form a precipitate with the guanidinium in the DNA binding buffer. The precipitate, however, is removed by the aqueous ethanol solution in the washing step. The sample is retrieved in water (20 µL) and pools were analyzed by 15% TBE-Urea gels.

#### BAL-31 Nuclease Digest

To ligated, double-stranded minicircles (~25 µL per pool, 10 pmol each, 50 pmol total) in water or 1X ligase buffer, equal volumes of 2X BAL-31 buffer [40 mM Tris-HCl (pH 8.0), 1,200 mM NaCl, 24 mM MgCl_2_, 24 mM CaCl_2,_ and 2 mM EDTA] and 1 µL of BAL-31 nuclease (2U/µL) enzyme were added. EDTA deactivation after ligation must be omitted as divalent cations are cofactors for the BAL digest. Samples were incubated at 30 °C for 10 min. The BAL-31 enzyme was deactivated with 5.6 µL of 300 mM EGTA (final concentration of 30 mM) and pools were analyzed by 15% TBE-Urea gels.

#### S1 Nuclease Digest

To ligated, double-stranded minicircles (~25 µL per pool, 10 pmol each, 50 pmol total) in water or 1X ligase buffer (EDTA deactivation should not be done), add 6 µL of 5X reaction buffer (200 mM sodium acetate (pH 4.5 at 25 °C), 1.5 M NaCl and 10 mM) and 0.5 µL of S1 nuclease (100U/µL) enzyme. Samples were incubated at 25 °C for 30 min. The enzyme was deactivated with 3.3 µL of 300 mM EDTA (final concentration of 30 mM) and pools were analyzed by 15% TBE-Urea gels.

#### Denaturing Gel Electrophoresis

15% Novex(TM) TBE-Urea gels containing 7 M urea were purchased from Thermo Fisher Scientific and run in a chamber filled with 1X TBE buffer that was pre-heated to ~55 °C at 200 V for around 2.5 h. To maintain temperature, the electrophoresis chamber was placed in an isolating Styrofoam box that was filled with ~1 L of hot water (~55 °C). Loading volumes were calculated such that each band contained 1.25 pmol per oligonucleotide. For the typical 5 oligonucleotides in a pool, a total of 6.25 pmol of template oligonucleotides were loaded per lane, mixed with 2X denaturing loading buffer (50% formamide, 10 mM NaOH, 0.01% bromophenol blue, 0.02% xylene cyanol) in equal volumes and loaded in the sample wells. 2 µL of 0.05 µg/µL stock of the GeneRuler ultra-low range DNA ladder (Thermo Fisher Scientific) was added in denaturing loading buffer as a reference. When electrophoresis was complete, gels were post-stained in 30 mL of a solution of 1X TBE (final) and 1X SYBR Safe DNA gel stain. To this, an additional 5-10% ethanol (v/v) were added to prevent the stain from adhering to the plastic dish during staining. The gel was shaken in the staining solution on a rocking plate for 15 min and finally imaged on a GE Typhoon FLE 9500 gel scanner using a 473 nm excitation laser and a 510 nm long pass emission filter. The photo-multiplier tube (PMT) gain was set to 500 and the pixel size was 25 µm. Images were analyzed by Fiji imageJ ^67^.

### Computational Methods (1): oxDNA Simulations

#### Preparing Initial Configurations

Initially, we used oxView to create our minicircles by creating polygonal shapes out of straight duplexes and ligating the corners to form a topologically closed catenane. OxView is a browser-based application (URL: oxview.org) that allows one to create initial configurations, connect to a server for relaxing initial configurations, view simulation trajectories, create movies of trajectories, and other interactive tools ^68,69^.

More recently, we used a legacy oxDNA analysis tools utility program called *generate*.*py*, which can generate perfect, non-eccentric minicircle configurations. In both cases, fully ligated rings were then relaxed using default initial CPU relax (default oxView settings: 1e5 steps MC at 30 °C) and a longer final GPU relax (default oxView settings: 1e6 steps MD at 20 °C). We generally added the nick to the minicircles after the relaxation step, resulting in starting production runs in a stacked conformation. Although this is a slightly biased initial configuration, it is unlikely that starting with a stacked configuration significantly biases the data, because transitions between kinked and stacked configurations are frequently observed (see Figure 1C-E and lifespan analysis in Supplementary Figure S5). Sequences of minicircles were all chosen randomly.

#### OxDNA MD Parameters

Simulations were run using the oxDNA2 model, which incorporates minor and major grooves, with the following parameters unless otherwise specified. The typical length of a production run was 500 million MD time steps with a print interval between snapshots of 10,000 time steps (totaling 50,000 printed simulation snapshots). The time scale between simulation steps, dt, was 0.005 time simulation units (~0.015 ps oxDNA time). The total time duration of our simulations is ~7.5 μs with a print interval of ~0.15 ns. Simulations were run at *T* = 20 °C = 293 K and [NaCl] = 0.5 M. The so-called “john” thermostat was used with defaults of 103 Newtonian steps and a diffusion coefficient of 2.5. The average-base model was used, in which the stacking and base-pairing interaction energies are identical regardless of sequence. The random number generator was uniquely seeded for all production and relaxation runs. All other parameters were kept at the default, as specified in the example MD input files provided in the oxDNA package. All simulations and analyses were carried out at the Ohio Supercomputer Center (http://osc.edu/ark:/19495/f5s1ph73).

##### Structural Kink Detection

The structural kink detector is based on the method described by Harrison *et al*. ^20^. The angles *θ*_*i*_ are determined between two base-normal vectors, 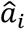 and 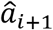, of two consecutive bases on the intact strand, which is shown in Supplementary Figure S2A. When two bases are stacked in the oxDNA model, typically 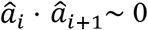, or, equivalently, *θ*_*i*_ is small. Five total angles 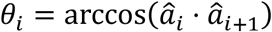 are determined between each consecutive pair of base-normal vectors in this region (Supplementary Figure S2B). We also define the max angle *θ*_*max*_ = max(*θ*_1_, *θ*_2_,…, *θ*_5_). Harrison *et al*. originally defined a configuration as stacked when *θ*_*max*_ < 90°, but we chose *θ*_*max*_ < 40° for a configuration to be considered stacked after studying probability distributions of *θ*_*i*_ in nicked minicircles.

Moreover, we added one additional condition. There exists one edge case that results in a false positive stacked readout in Harrison’s method ^20^. It is possible that the normal of both ends of the helix about the nick site are approximately parallel to each other, yet the ends are disjointed, as in Supplementary Figure S2C. The effect on the results is typically small, usually affecting only about 1% of the readouts in our typical minicircle simulation runs and is most noticeable in minicircles with a small, stacked fraction. The false positive stacked configurations can be recognized by an additional distance check. A configuration is considered stacked by the structural kink detector if 1) *θ*_*max*_ < 40° and 2) the backbone sites of the two nucleotides nearest to the nick site on the nicked strand are separated by a distance *R* ≤ 2 simulation units of distance (~1.7 nm).

#### Error Analysis

For Figure 1B, each simulation was split into four equally long quarters. The statistics in each quarter of the simulation were determined independently. The propagated error was taken to be the standard deviation of the independent fractions in each quarter. This is essentially equivalent to analyzing four independent simulations consisting of 125 million steps, as the typical duration of kinking events for the minicircles with a single nick is short (Supplementary Figure S5), making the results independent of the initial configuration.

#### Kymograph Generation

(Figure 1, Supplementary Figure S4). For each configuration, the conformation of the ring is analyzed. To determine the conformation of the ring, the stacking interactions and hydrogen-bonding potentials of the duplex were used. Blue data points represent kinked configurations and red data points represent stacked configurations. To be precise, for a configuration to be colored blue, the duplex must be kinked at the nick site. If the duplex is not kinked at the nick site, the configuration is considered stacked and colored red. The average twist angle, *ϕ*, of the dinucleotide step is computed for each configuration, where the ending two base pairs of both terminals of the nick site were excluded from the analysis as these are prone to fraying. The twist angle between adjacent base pairs is computed using a scheme described in the supplement of ref. ^52^.

#### Tension Analysis

One stationary harmonic trap was placed for each nucleotide at both ends of the duplex such that the coordinates of the trap, 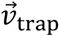, was chosen to be the COM (center of mass) coordinates of the nucleotide in the starting configuration with (high) spring constant *k* = 1000 model units of spring constant ~ 57000 pN/nm. This totals to four harmonic traps, with two for each complementary base pair at each end of the duplex. The force of tension on each nucleotide of the base pair was then determined independently for each snapshot of the trajectory. First, the instantaneous displacement 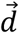 of the COM of a nucleotide from the harmonic trap’s location is

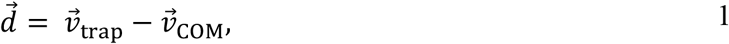

where 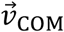 is the COM coordinates of the nucleotide in the current configuration.

The harmonic trap exerts a force

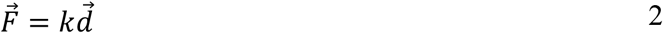

on the nucleotide. The instantaneous local tangent vector of the helix was defined as

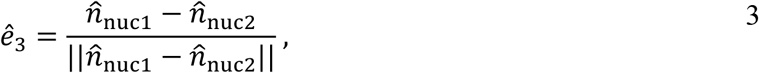

where 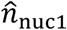 and 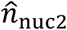 are the base-normal vectors of the first and second nucleotide of the base pair, respectively. The component of the force along the local tangent of the helix then is

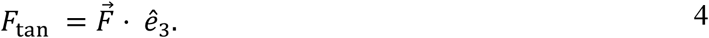

Once this has been computed for both nucleotides, the average force along the local tangent of the helix felt by each nucleotide is reported as

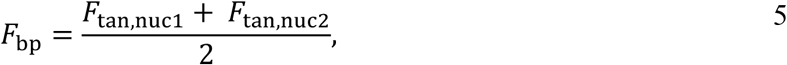

where *F*_tan,nuc1_ and *F*_tan,nuc2_ are the components of the force along the local tangent for the first and second nucleotides in the base pair, respectively.

In addition, any configuration containing any melted complementary base pairs was excluded from the analysis. A base pair is considered melted if the model hydrogen-bonding interaction energy |*V*_HB_| < .1 (max |*V*_HB_|). This resulted in the exclusion of approximately 15% of snapshots from each run of the circularly bent 81-mer results shown in Supplementary Figure S10.

### Computational Methods (2): All-atom MD Simulations

Nicked DNA circles of variable length (same sequences as experimental, see Supplementary Table S1) were constructed with the MC DNA web server ^70^ and modified with BIOVIA Discovery Studio ^71^ to ligate one of the two strands that were initially unconnected. Each system was solvated in a cubic box of explicit TIP3P water and neutralized with sodium ions, followed by adding 0.15 M additional Na and Cl ions. The nicked circles were relaxed through the steepest descent minimization approach (until the maximum force, F_max_, is no greater than 1000 kJ mol^-1^ nm^-1^) to ensure that the system has no steric clashes or inappropriate geometry. After minimization, all systems were equilibrated at constant N, V, and a constant temperature of 300 K for 100 ps, followed by a 100 ps equilibration in the NPT ensemble at 1 bar before MD runs. Bonds involving hydrogens were constrained by the LINCS algorithm ^72^, and short range non-bonded interactions were cut off at 0.9 nm. Long-range electrostatic interactions were treated by the particle mesh Ewald method (PME) ^73^. The cubic simulation box had dimensions ranging from approximately 13 to 14.5 nm with a periodic boundary condition.

Production runs were performed at a constant pressure of 1 bar and a temperature of 300 K using a V-rescale ^74^ thermostat and a C-rescale barostat. The MD time step was 2 fs, and the duration of the run was 50 ns.

### Preparation of Nicked Circles Containing HMfB Proteins

To create a model of a nicked minicircle with two HMfB dimers, the structure for the archaeal histone-DNA complex (Protein Data Bank PDB ID: 5T5K) was modified using PyMOL and BIOVIA Discovery Studio 2021. We removed the two nucleotide sticky ends, 8 bp from one end of the DNA duplex and two histone proteins. Two phosphodiester bonds were manually constructed between the O3’ and P on the free ends of the duplex to form a continuous, ds DNA circle. The manually created phosphate bonds were overstretched; hence, this was corrected by a 150 ns simulation. The last frame of the simulation was extracted and used as the initial structure for the production run. The extracted structure was placed in a cubic box with dimensions approximately (12.7 nm)^3^, ensuring a minimum distance of 1.0 nm between the DNA and box boundaries. The protein-DNA system was solvated in explicit TIP3P water and neutralized with sodium ions. All subsequent simulation parameters (energy minimization, equilibration protocol, thermostat, barostat, constraints, and electrostatics treatment) were identical to those described above. All molecular dynamics simulations were performed using GROMACS 2024.4 ^75^, employing the AMBER14 force field for the protein and the AMBER OL21 force field for DNA^76,77^. In addition, the CUFIX corrections were applied to correct the overestimated ion-phosphate interactions ^78–80^. All simulations were run on GPUs at the Ohio Supercomputer Center ^81^. PyMOL ^82^ (Schrödinger, LLC) was used for visualizations and preparation of snapshots.

### Analysis of MD Trajectories

The twist at each base pair step was calculated using do_x3dna ^83^, which employs the 3DNA package ^84^. The twist averaged over the base pairs was calculated using a Python script written in-house. The helical repeat was calculated from these averages over the span of the simulations, plotted using an in-house Python script.

The geometrical criterion defined in the GROMACS manual is used to determine the existence of a hydrogen bond. The donor-acceptor distance must be less than or equal to 0.35 nm and the hydrogen-donor-acceptor angle must be less than or equal to 30°. Only one hydrogen bond per Watson-Crick base pair is analyzed to determine if the bond is intact or not. The donor-acceptor distance and the hydrogen-donor-acceptor angle are measured using the ‘distance’ and ‘angle’ GROMACS utilities.

Base stacking is characterized geometrically. The stacking definition proposed by the Florian group ^85^ is used. The distance between the centers of mass of two nucleobases and the angle between their planes are combined to produce a single reaction coordinate, χ. Adjacent nucleobases are considered stacked if χ is less than or equal to 0.6 nm. The plane of a nucleotide was determined using the definition presented by the Turner group ^86^. The plane of a nucleotide is defined by two vectors, *a* and *b*, whose cross products define each base’s normal vectors. For purines, *a* is defined from the center of mass (COM) to the C8 atom and *b* is defined from the COM to the nitrogen (resp. oxygen) attached to the C6 atom in adenine (resp. guanine). For pyrimidines, *a* is defined from the center of mass to the oxygen group attached to the C2 atom and *b* is defined from the center of mass to the oxygen (resp. nitrogen) attached to the C4 atom in thymine (resp. cytosine). The vector and COM coordinates were measured using the GROMACS ‘traj’ utility. Stacking is considered broken for a base pair when either nucleotide stacking interaction is not intact. Python scripts were written in-house to plot analysis data.

## Supplementary Materials For

### Supplementary Text

#### Supplementary Note 1: Classical looping experiments are ill-suited to determine h_0_ of tightly bent DNA

Circularization or looping experiments with DNA fragments of different lengths are historically the most important experiments to extract the equilibrium helical repeat as well as bending and torsional stiffness constants ^29–33^. For this, the *j*-factor ^23,87^ was introduced, which is defined as

**Figure S1.**
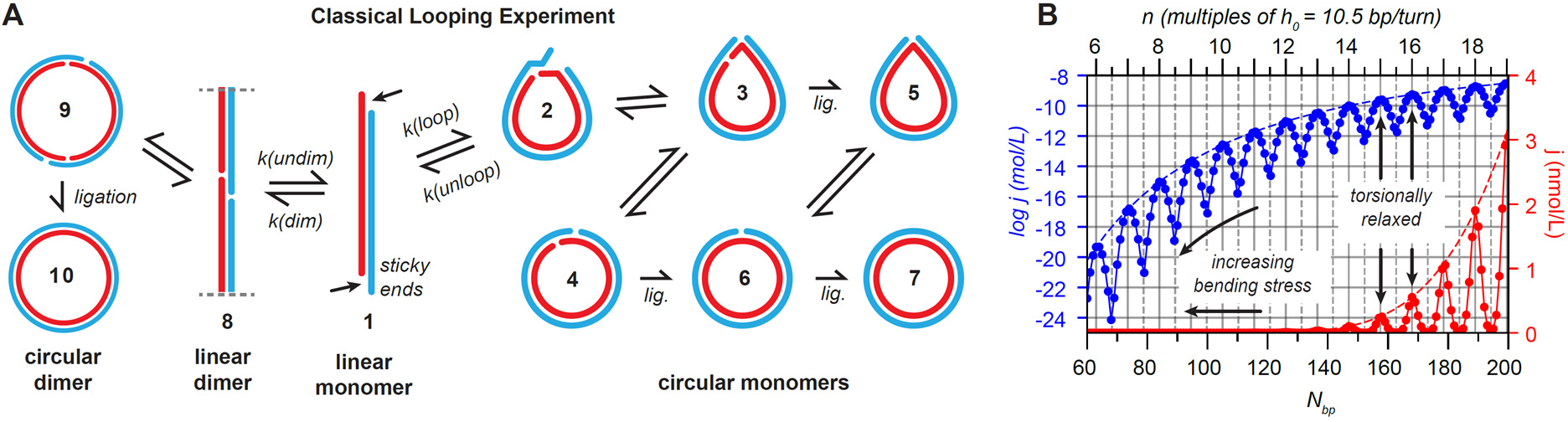
Classical circularization experiments. (**A**) In a classical looping or circularization experiment ^29–33^, a linear monomer (**1**) with self-complementary sticky ends (arrows) can circularize and be ligated into a double-stranded circle (**7**). However, this transition does not occur directly. Instead, (1) is in equilibrium with looped monomers (**2-7**) and linear and circular dimers (**8-10**). The final ligated conformation (**7**) is formed through various kinked transition states (**2, 3, 5**) with reduced bending and torsional stress ^15,19–21^. (**B**) Plot displaying j-factors, or theoretical circularization probabilities for DNA fragment lengths of N_bp_ = 60 − 200 bp calculated from the Shimada-Yamakawa model (Ref. ^23^, Supplementary Note 5). **Blue**: logarithmic scale; **Red**: linear scale.

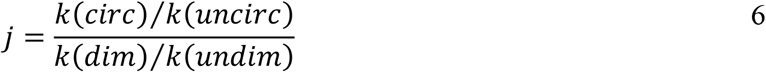

and can be interpreted as the effective molar concentration of the two ends of the linear fragments, where a small *j*-factor indicates a low looping probability. Figure S1B demonstrates that circularization becomes highly improbable for contour lengths around and below the persistence length (~150 bp or 50 nm) due to the increasing bending stress. For fragment lengths with a non-integer number of helical turns, a twist energy must be overcome to align the fragments and enable circularization. Consequently, the *j*-factor has an oscillating subpattern with a period corresponding to the equilibrium helical repeat *h*_0_ (Figure S1B).

The classical TWLC model and *j*-factor graphs assume *h*_0_ is constant and independent of curvature. In fact, circularization results for DNA fragments that are longer than the persistence length are in excellent agreement with the TWLC model, the canonical persistence length *l*_*p*_ ≈ 150 *bp* ≈ 50 *nm* and an equilibrium helical repeat of *h*_0_ = 10.45 ± 0.05 *bp*/*turn* ^29,30,32^.

However, some experiments with DNA fragments around or below 100 bp showed much higher circularization efficiencies than expected, which seem to contradict the TWLC model, and extreme bendability was suggested as an explanation ^5,14^. This interpretation was, however, criticized due to the chosen melting temperatures of sticky ends and other experimental conditions that heavily change kinetic constants ^16,15,20^. Specifically, the melting temperature of the sticky ends must be low, and the ligase must be very dilute to ensure that the equilibria between the looped and linear states are not influenced. This condition was not given in those experiments.

Furthermore, the single-stranded sticky ends are very flexible, and therefore looping is likely to be initiated in a distorted teardrop-like shape and fully hybridized looped fragments can release mechanical stress through sharp bending or “kinking” at either of the two nicks, and teardrop shapes exist in equilibrium with circles (^15,19–21^, Figure S1). Such kinks provide a mechanism to release torsional strain and can explain why torsional constraints for the helical repeat did not influence looping of short fragments as much as expected. Finally, the exact orientation of the sticky ends in respect to the center-helix axis strongly influences the looping kinetics ^21^ which also strongly influences the rate-limiting transition from **1**→**2**. Eventual torsional stress can be relaxed at a kink in state **3** and FRET experiments cannot distinguish the states **2-4**. For these reasons, Kim argued that the *j*-factor is not an experimentally accessible quantity for fragments around or below 100 bp anymore ^21^. Their analysis of the decyclization kinetics (**2**→**1**) shows an increased helical repeat consistent with our data and interpretation ^21^.

In contrast, our nicked minicircle assay (Figure 1) can only fluctuate between a nicked circular that is identical to intermediary step (state **5**) and a stacked circular state **6**. The probability for these states depends mostly on torsional stress.

##### Production of j-factor graphs

The Shimada-Yamakawa (SY) theory for the twisted worm-like chain (TWLC) details an analytic solution for the *j*-factor applicable to small length *L* helical polymers. The result is reproduced here for convenience without major modifications from ref. ^23^. The *j*-factor (in units of mol/L) can be expressed as

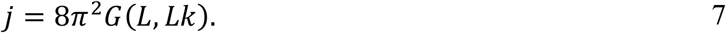

The SY expression for *G*(*L, Lk*) is

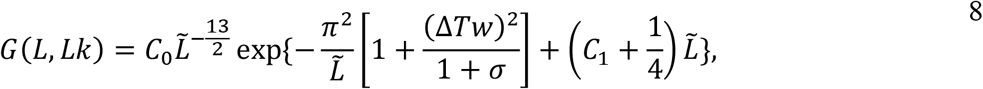

where 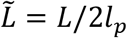, and *σ* is the Poisson’s ratio defined by

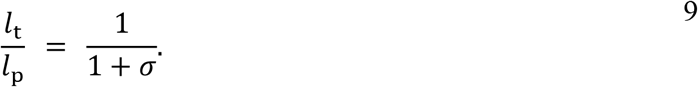

The parameters *C*_0_ and *C*_1_ are given by

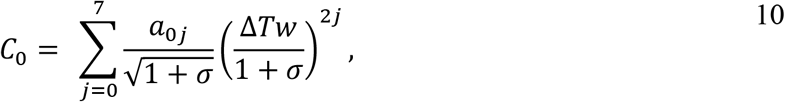

and

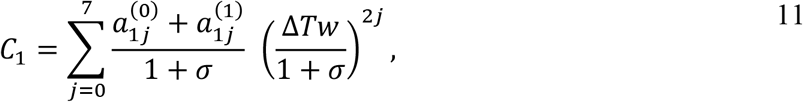

with parameters in the sums given by

| $j$ | $a_{0j}$ | $a_{1j}^{(0)}$ | $a_{1j}^{(1)}$ |
| --- | --- | --- | --- |
| 0 | 2.784 | 0.2639 | -0.0383 |
| 1 | 2.113 | 0.1399 | -0.0827 |
| 2 | 0.6558 | -0.1131 | 0.0125 |
| 3 | 1.719 | 0.6500 | -0.2170 |
| 4 | -2.478 | -1.1223 | 0.3961 |
| 5 | 2.588 | 1.0320 | -0.3991 |
| 6 | -1.210 | -0.4601 | 0.1899 |
| 7 | 0.2437 | 0.0829 | -0.0367 |

The TWLC *j*-factor is a sum of contributions of *j*_*ΔTw*_ over all Δ*Tw* such that the linking number *Lk* = *Tw*_0_ + Δ*Tw* is an integer (eq. 9),

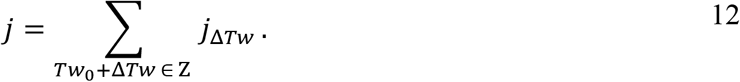

All values of Δ*Tw* that meet this condition can be generated via

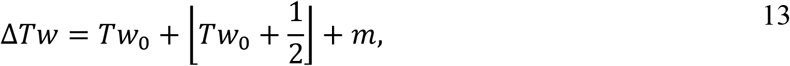

where *m* is an integer, written so that increasing |*m*| generally increases |Δ*Tw*|.

If the condition

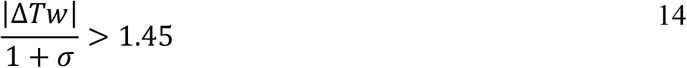

is met, the minimum configuration has significant writhe, so it is omitted from the sum. So, the final *j*-factor as plotted in Supplementary Figure S1B (with *h*_0_ = 10.5 bp/turn, *l*_p_= 150 bp, and *l*_t_ = 220 bp) is effectively

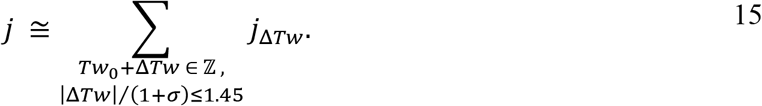

For these parameters, this sum includes at most three contributions each with |Δ*Tw*| ≲ 1. To convert from molecules per volume to molar concentrations, one must multiply by scale factor

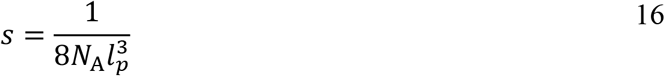

where *N*_*A*_ is Avogadro’s number and 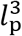 is written in units of liters (L). Finally, the maximum envelope (dotted line Fig. S1B) is

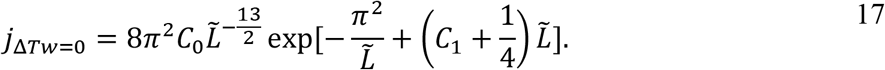

#### Supplementary Note 2: The Magnitude of Experimentally Observed Twist-Bend Coupling

Marko and Siggia (MS) introduced a model that extends the TWLC and includes twist-bend coupling due to the asymmetry of bending about the major and minor grooves ^36^. MS calculated the excess twist density perturbatively for small κ/ω_0_, where κ is the curvature and ω_0_ is the intrinsic relaxed twist density expressed in rad/nm. For a circularly bent duplex with constant curvature κ, twist-bend coupling leads to an unwinding of DNA with twist of up to second order in κ/ω_0_

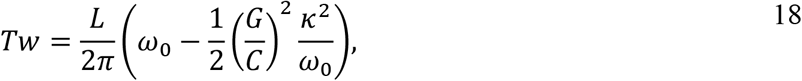

where the length of DNA, *L* = *N*_bp_*l*_bp_, is expressed in terms of the number of base-pairs *N*_*bp*_ with relaxed helical rise *l*_bp_ = 0.34 nm, *C* is the twist stiffness, and *G* is twist-bend coupling parameter (in notation of ref. ^53^). For a circularly bent duplex with κ = 1/*r* = (2*π*/*L*), eq. 18 can be written in terms of the helical repeat as

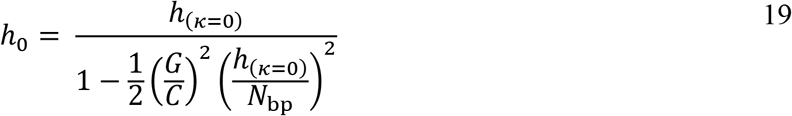

where *h*_(K=0)_ = 2*π*/ω_0_*l*_*bp*_ denotes the relaxed helical repeat when unbent. Fig. 4B shows the relaxed helical repeat inferred from the split bands in our ligation experiments of nicked minicircles, and the relaxed helical repeat for longer minicircles determined in classical circularization experiments reported decades ago ^29–33^. Eq. 19 provides an excellent fit to both sets of data with fitting parameters *G*/*C* = 2.7 and *h*_(K=0)_ = 10.42 *bp*/*turn*. Indeed, these experiments were originally highlighted by MS who concluded that *G*/*C* 2.4 could account for the change in helical repeat. The change in the relaxed helical repeat of the smaller minicircles (order 10%) is much larger than the previous reported measurements of larger circles (order 1%).

The close agreement of the experimental data with functional form given by eq. 19 (fit in Figure 7) suggests that the widening of the helical repeat is a consequence of twist-bend coupling. Still, a value of *G*/*C* > 1 is puzzling. Stability of the MS model requires *A*_2_*C* > *G*^2^ and (*A*_1_ + *A*_2_)*C*/2 > *G*^2^ where *A*_1_ and *A*_2_ the two bending coefficients. These relations are also required for the renormalized bend and twist coefficients derived for the MS model recently ^88^. These conditions are not easily satisfied when *G*/*C* > 1. Estimates based on oxDNA2 model ^52^ and fits to magnetic tweezer experiments ^88^ suggest values of *G*/*C* = 0.3, about an order of magnitude smaller than suggested by the measured helical repeat in our data. For *G*/*C* = 0.3, the change in helical repeat due to TBC is not significant for minicircles of this size range, as shown in Figure 7A.

#### Supplementary Note 3: Conditions for Forming Different Topoisomers Without Kinks (Interpretation 1)

In the following, we are discussing topoisomer splitting without internal kinks in more detail (Interpretation 1; see Figure 3D-F).

Supercoiling is often observed in DNA plasmids, where the linking number, *Lk*, is distributed between twist, *Tw* (in helical turns), and writhe, *Wr*, such that

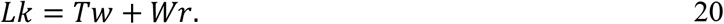

The linking number is an integer denoting the number of helical repeats in a ligated circle. The helical pitch (in number of base pairs per helical turn) of a duplex with *N*_*bp*_ base pairs is

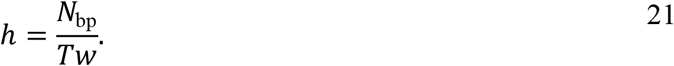

Since writhe is experimentally not observed for circles shorter than 174 bp ^51^, *Lk* ≈ *Tw* for the smaller, ligated circles in our experiments. For ligated circles, *Tw* takes on approximately integer values, but for nicked circles, *Tw* can adopt non-integer values via kinking and twisting (see Figure 1G)

We define the relaxed helical repeat *h*_0_ as the value of *h* where there is no torsional strain. Any other *h* will be either underwound (*h* > *h*_0_) or overwound (*h* < *h*_0_).

Alternatively, the twist (or linking number *Lk* in ligated circles) can be written as

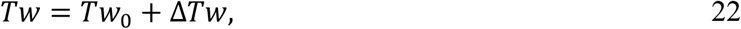

where

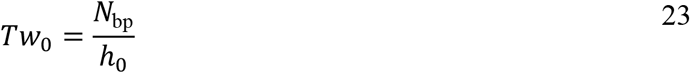

is the number of turns a linear or nicked circular duplex makes without torsional strain, and Δ*T*w is the number of turns relative to the torsionally relaxed conformation needed to form a minicircle with linking number *Lk*. Here, Δ*T*w > 0 and Δ*T*w < 0 correspond to overwound and underwound duplexes, respectively.

To understand why band splitting only occurs near 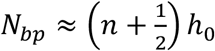 for integer *n* according to our interpretation 1 (Figure 3D-F), we consider the probability of forming a minicircle of radius *r* for a DNA segment with length *L* and linking number *Lk*. Within the TWLC model ^23^, the energy to form a perfect circle of radius *r* is given by

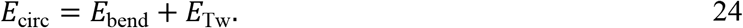

where *E*_bend_ is the bending energy necessary to form a circular conformation, and *E*_Tw_ is the torsional energy necessary to rotate the ends to be in register (i.e., so that *Tw* is an integer). The bending energy can be expressed as

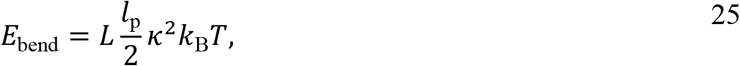

where κ is the curvature, *l*_p_ is the bending persistence length, and *k*_B_*T* is the thermal energy scale, and *L* is the contour length of the DNA. For a circular conformation of radius *r, L* = 2*πR*, and the curvature is κ = 1/*r* = 2*π*/*L*, so that the bending energy can be expressed as

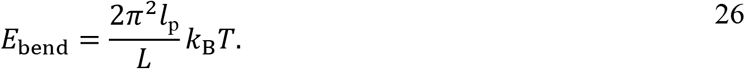

The twist energy can be written as

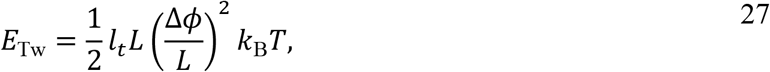

where *l*_*t*_ is the torsional persistence length, and Δ*ϕ* is the rotation needed to put the ends in-register. Substituting Δ*ϕ* = 2*π*Δ*T*w, the twist energy becomes

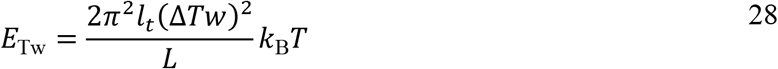

For each linking number, the local minima of *E*_circ_ corresponds to DNA lengths with zero torsional strain; that is, *N*_*bp*_ ≈ *nh*_0_ which has a relaxed twist *Tw*_0_ equal to the linking number. Assuming the ligation yield in the experiment is proportional to the probability of forming a stacked conformation, i.e.

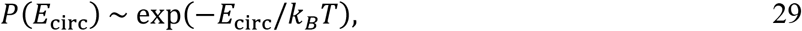

the yield is maximal for torsionally relaxed minicircles. For other DNA lengths, the dominant topoisomer has a linking number corresponding to the lowest energy configuration. Consequently, for most DNA lengths, only ligated circles with one *Lk* is observed because topoisomers with higher torsional energy are unlikely (Figure 3I). When *N*_*bp*_ ≈ (*n* + 1/2)*h*_0_, the energies of two adjacent linking numbers are comparable, so that a topoisomer with linking number *Lk* + 1 and Δ*Tw* = 1/2 and another with linking number *Lk* and Δ*Tw* = −1/2 form with roughly equal probabilities (Figure 3D). Adding or subtracting an entire helical turn (*ΔTw* = 1) as an alternative explanation of band splitting in previous experiments with circles of similar size ^37^ would incur a high twist energy and/or energetic cost for kink formation and is therefore physically implausible.

This argument focuses on the deformation energy of the minimum energy conformation of the TWLC for simplicity. The free energy for minicircles of a TWLC has been solved exactly ^33^, as well as by approximation by Shimada and Yamakawa (SY) that includes the perturbative harmonic fluctuations about the minimum energy ^23^. It is convenient to consider this SY approximation which has been shown to be accurate for DNA length shorter than 2.5 *l*_*p*_. The free energy for a minicircle of length *L* and linking number *Lk* is Δ*G*(*L, Lk*) = −*k*_B_*T* ln *G* (*L, Lk*) where *G*(*L, Lk*) is the probability of forming the minicircle. Within the SY approximation, the free energy can be written as

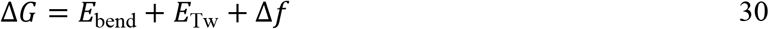

with *E*_bend_ and *E*_Tw_ given by Eqs. 26 and 28, and

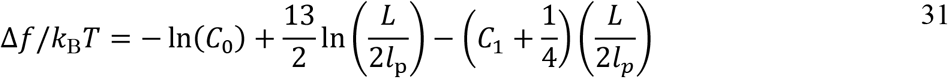

is the free energy due to harmonic fluctuations about a uniform minicircle of length *L* and linking number *Lk*. The parameters *C*_0_ and *C*_1_ are specified in the next section.

The free energy is about 10 *k*_B_T lower than the mechanical deformation energy due to uniform (non-fluctuating) circular conformation.

#### Supplementary Note 4: Tension in Minicircles

From experiments with molecular torque tweezers on long supercoiled DNA it is known that a low tension up to few pN increases the torsional stiffness *C* and stretches the DNA ^88,89^. Increasing tension further up to 20-30 pN induces a slight overwinding of DNA whereas even higher forces (tens of pN) unwind DNA ^45^. Likewise, application of torque influences the length ofthe DNA fragment. We initially hypothesized that this twist-stretch coupling might have to be considered in tightly bent nicked circles. For *N*_*bp*_ = 89, *L* = *N*_*bp*_⋅0.34 *nm*⁄*bp* = 30 *nm* and *l*_*p*_ = 50 *nm, E*_bend_ ≈ 33 *k*_*B*_*T* (eq. 25).

The force *F* to keep ends of a linear strand in a looped conformation is the derivative of the looping energy with respect to the circumference *L*:

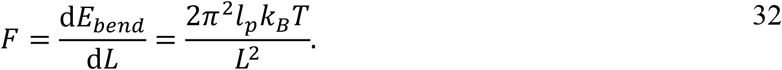

Next, the thermal energy *k*_B_*T* ≈ 4.1 *pN* ⋅ *nm* at room temperature. For an *N*_*bp*_ = 89 circle, a tension of *F* ≈ 5 *pN* is expected anywhere in the circle. This calculated force is ~27 % higher than the force to keep ends in a looped conformation in oxDNA (see Supplementary Figure S10 and Methods), which we attribute to teardrop deformations in the simulation. In any case, tension is so small that twist-stretch coupling is negligible ^45^, but large enough to increase the torsional persistence length to about ~100 nm in small circles ^89,90^.

#### Supplementary Note 5: Model for Topoisomer Splitting due to Internal Kinking

The model for topoisomer splitting of circular DNA involving disturbances and kinked minicircles (Figure 3G, interpretation 2) was proposed by Du and Vologodskii ^37^. According to this model, roughly equal topoisomer populations are formed with *ΔTw* ≈ 0 and highly unwound kinked minicircles with *ΔTw* ≈ −1. In the following we discuss the energetics required for the model to describe the observed topoisomer splitting within the harmonic energetic plots like Figure 3F.

Within the harmonic elastic limit, the free energy of a minicircle without kinks of length *L*, the torsional persistence length *l*_*t*_ and excess twist *ΔTw* can be written as the sum of the twist energy and bending energy

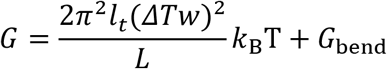

A kinked minicircle on the other hand has disturbances which relieve some bending stress of the minicircle and provide a local unwinding δ*Tw*. The local unwinding is not constrained by the harmonic twist constraints of the intact DNA, so that twist energy depends on *Δ*Tw – δTw. So, the free energy of a kinked minicircle can be written as

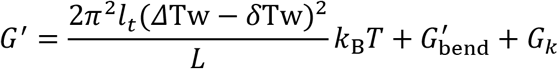

where 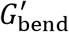 is the bending energy of the kinked minicircle and *G*_*k*_ is the free energy associated with the local disturbance due to broken base pair interactions for example. Since the kinks reduce the bending stress, we can write the bending energy of the kinked minicircle as

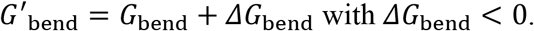

Figure S2a shows a plot of the free energy as a function the number of base pairs for two intact minicircles and one kinked minicircle. Relative to undisturbed circle (Lk = 6), the free energy of the kinked minicircle is shifted towards larger circles by an amount determined by free twist of the kink, *h*_0_δ*Tw*, and stabilized by δ*G*_*k*_ = *ΔG*_bend_ + *G*_*k*_. The crossing of free energy curves of adjacent linking numbers for the kinked circle (6*) and unkinked circle (7) also shifts to larger circles compared to the crossing of unkinked minicircles.

**Figure S2.**
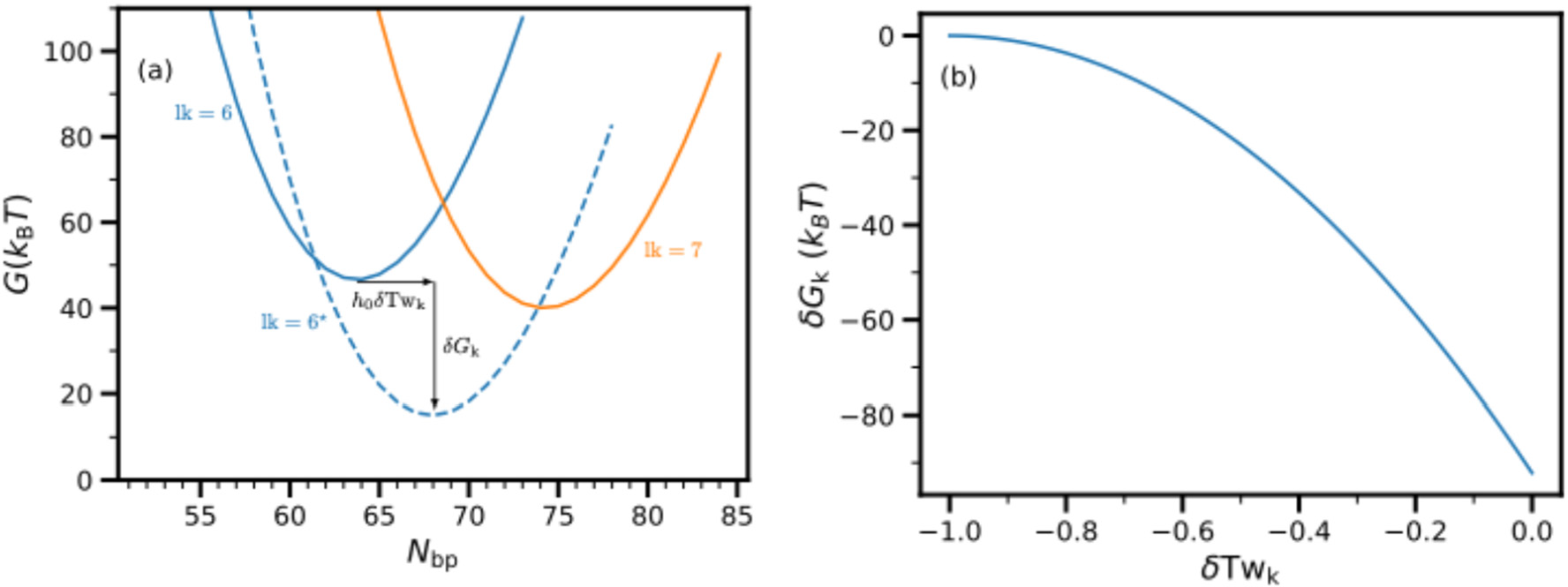
Energy stabilization from kinking. Free energy vs number of base pairs for intact minicircle (solid lines) for linking number Lk = 6 (blue) and Lk = 7 (orange). Free energy for kinked minicircle (dashed) for linking number Lk = 6 with ΔTw = −0.75. The bending free energy is taken to be the bending energy given in eq. 26. (b) Stabilization free energy vs the amount of free twist required for topoisomer splitting to occur at ΔTw = 0 and ΔTw = −1. Parameters for these plots are l_p_ = 51nm, h_0_ = 0.55bp/turn, and l_t_ = 100nm.

Interpretation 2 postulates topoisomers transition to occur where one circle is relaxed with *Δ*Tw ≈ 0 and the other is underwound and kinked at *Δ*Tw ≈ −1. This gives the following relation between δTw and δ*G*_*k*_:

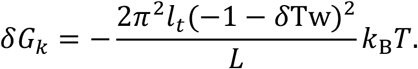

This stabilizing energy comes from the relaxation of the bending energy and the local free energy cost of the kinks, is shown in Fig S2 (b). Multiple kinks (for example two kinks along a diameter) allow for greater reduction in bending free energy than a single kink. Still, |δ*G*_k_| can not be greater than the bending energy of the intact minicircle suggesting that the free twist δTw is required for this mechanism to be substantial (δTw ⪆ 0.4 for linking number 6). Another way to say this is that when the free twist is relatively small (say δTw ⪅ 0.2) the stabilizing energy required to explain the topisomer splitting is unreasonably large.

## Supplementary Figures

**Figure S3:**
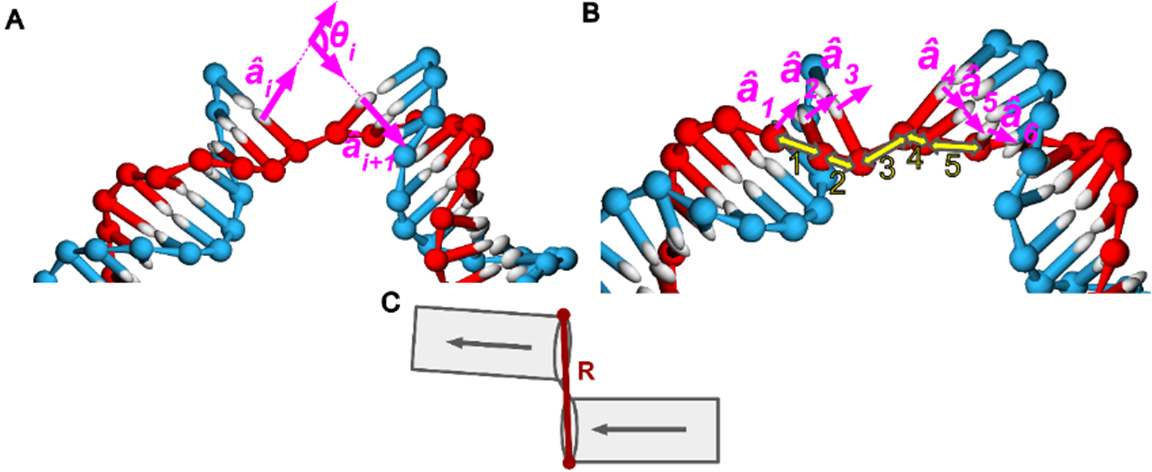
Geometrically detecting kinks. (**A**) Annotated oxView image of the angle θ_i_ between base-normal vectors of two consecutive bases on the intact strand. (**B**) Five total angles {θ_i_} are determined (between pairs indicated by yellow arrows) in a six-base region centred by the nick site. (**C**) Finally, a distance R is defined between the sites adjacent to the nick. A configuration is considered kinked if 1) max{θ_i_} > 40° or 2) R ≳ 1.7 nm.

**Figure S4.**
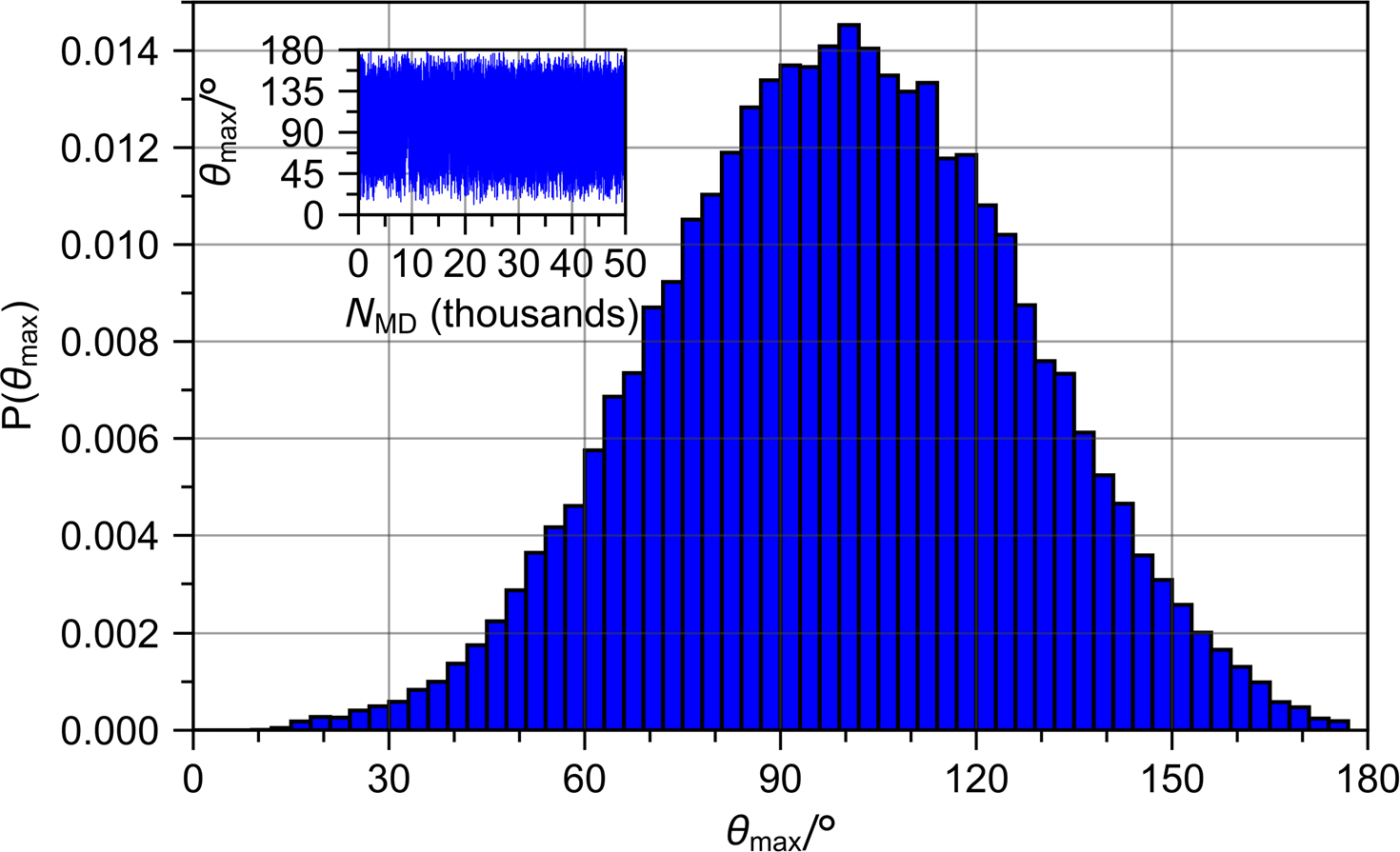
Kink angle analysis. Probability density of the maximum angle between base-normal vectors θ_max_, determined at the base pairs near the nick site for a nicked 89-mer minicircle in oxDNA (see Methods). Because the nicked 89-mer has a 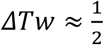 and is therefore nearly always kinked, this distribution is representative of the approximate bending angle for small, kinked minicircles. The mean of the distribution is approximately 100°. The inset plots the time-evolution of this bending angle as a function of simulation snapshot (50,000 total snapshots) and demonstrates the extremely fast fluctuations. No additional patterns or metastable states were observed.

**Figure S5:**
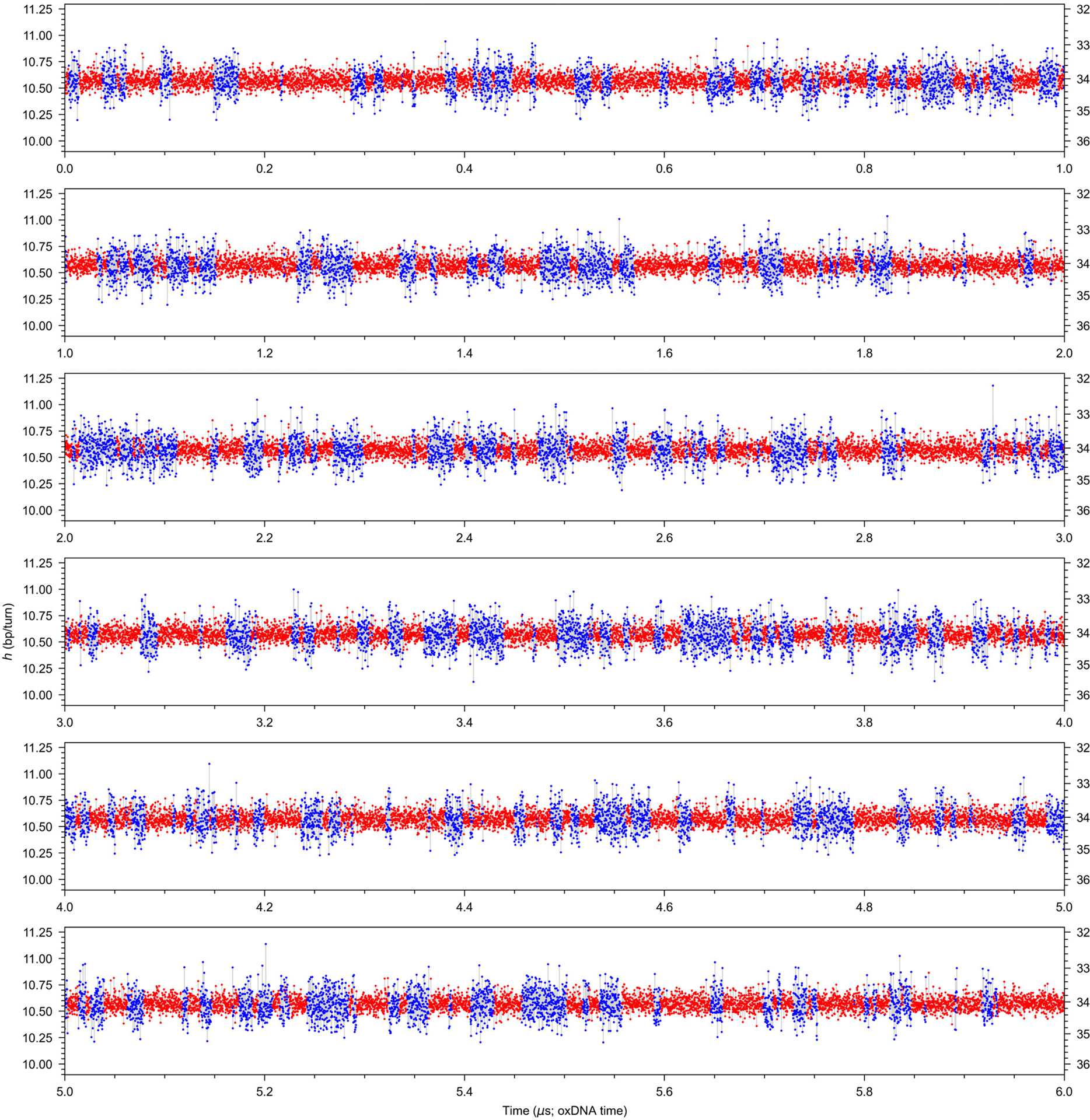
Kinking and stacking. Analysis of a 6 µs oxDNA simulation of a nicked 84 bp circle (ΔTw ≈ 0). The helical repeat (left scale) was calculated from the average angle between two consecutive base pairs (right scale) and plotted for each snapshot. Red: stacked configuration, blue: kinked configuration. Lifespan analysis in Supplementary Figure S5. Note that diffusion is accelerated in oxDNA and therefore processes in an experiment or all-atom MD simulations would likely be ~1-2 orders of magnitude slower ^39,40^.

**Figure S6:**
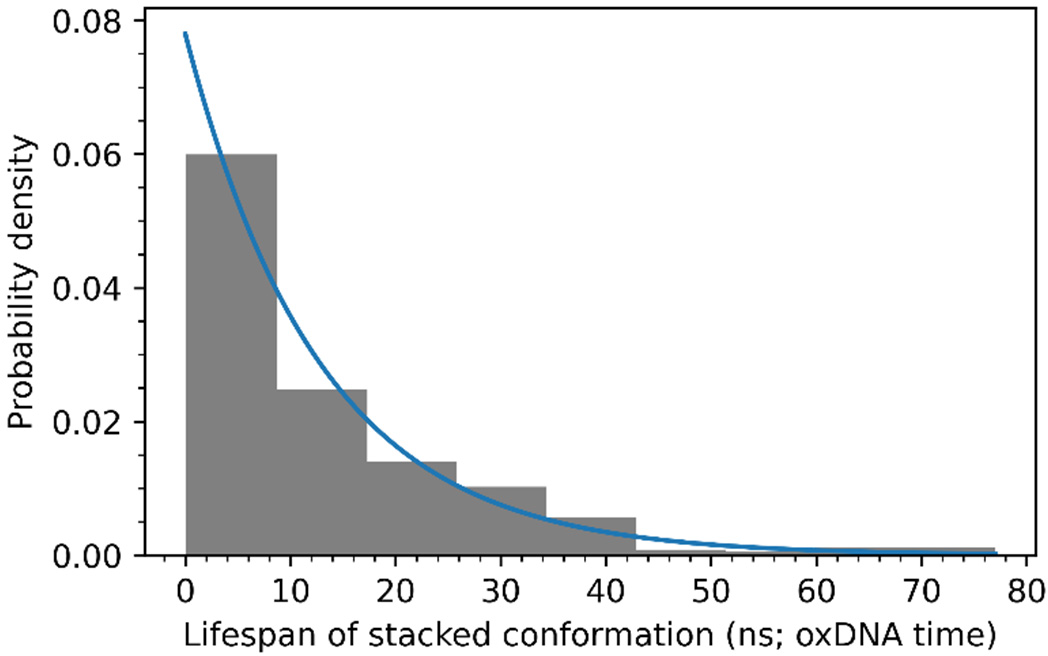
Lifespan of stacked conformations. The lifespan of stacked configurations of the oxDNA simulation of a nicked 84 bp (ΔTw ≈ 0) circle was analyzed. An exponential decay curve was fitted to the data. The number of uninterrupted sequences of stacked configurations consisting of at minimum two configurations observed was 337. The mean lifespan is ~13 ns with a corresponding standard deviation of 13 ns. Note that diffusion is accelerated in oxDNA and therefore lifetimes in an experiment or all-atom MD simulations would likely be ~1-2 orders of magnitude longer ^39^. Still, lifespans are many orders of magnitudes faster than the second or minute lifetimes in looping experiments ^14,21^.

**Figure S7.**
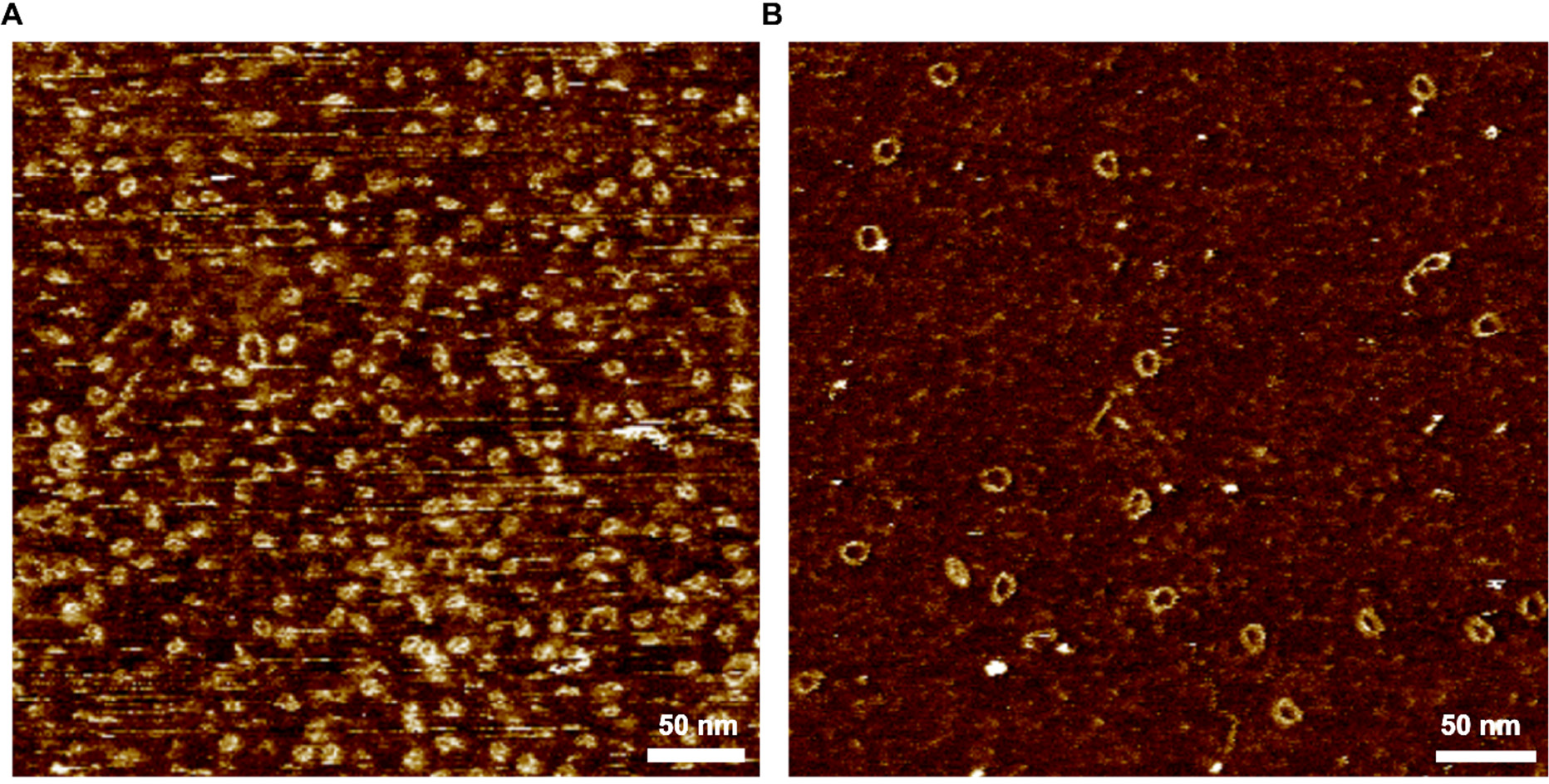
AFM images of ligated 72 bp (left) and nicked 103 bp circles (right). Even with a sharp tip that distinguishes the major and minor grooves in DNA, it was not possible to clearly determine if the minicircles were kinked or not, especially for smallest circles.

**Figure S8.**
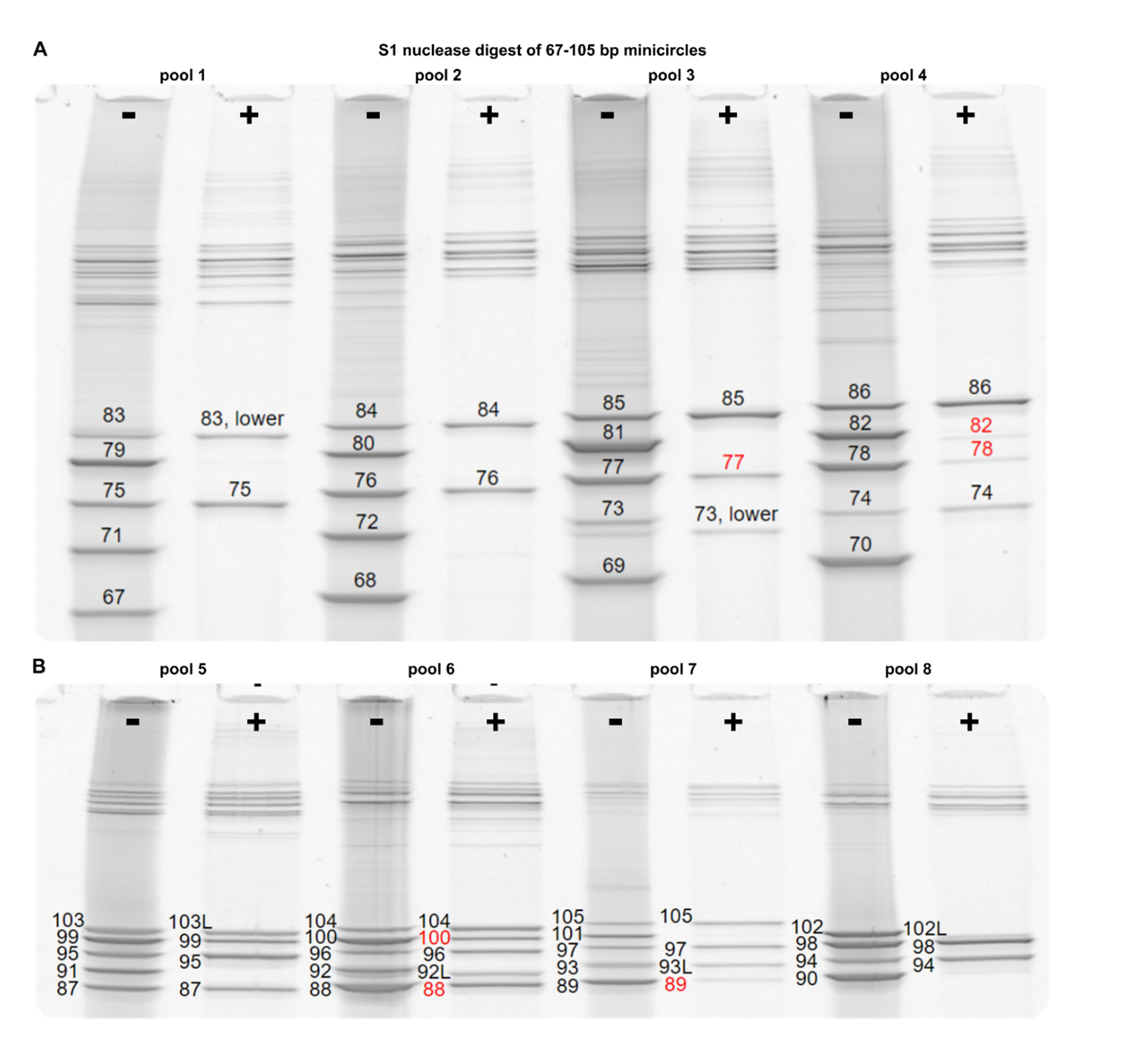
Digest experiments with S1 as an alternative to BAL 31 nuclease. Results show that S1 does not digest unwound as aggressively as BAL31 and some circles highlighted in red resist S1 nuclease treatment, but not BAL-31. Note that all circles with low helical repeat resist either nuclease.

**Figure S9.**
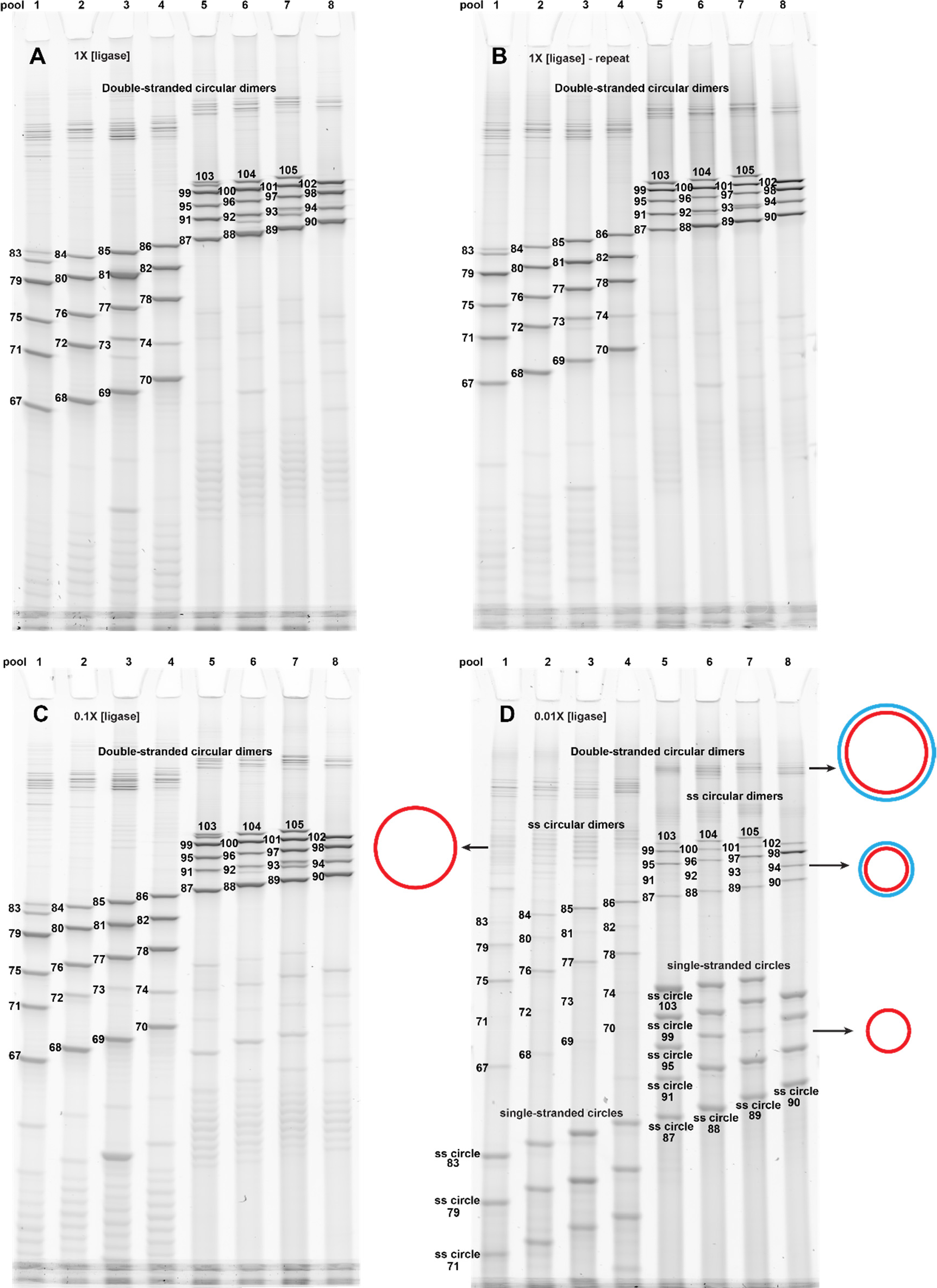
Uncropped gels of ligation experiments at different ligase concentrations. Lifetimes of both looped and linear configurations in classical circularization experiments can be on the order of seconds to minutes (ref. ^14,19^, Supplementary Figure S1, Supplementary Note 1) and ligations in these experiments must be performed at very low ligase concentrations to avoid shifting the equilibria ^8^. As ligation experiments take minutes to complete while stacking/unstacking in our assay has nanosecond kinetics (Supplementary Figure S5), we hypothesized that the concentration of the ligase will not influence equilibria in our assay. The starting concentration of oligonucleotides in all experiments was the same, but different ligase concentrations were used in the final ligation step of nicked circles. (**A**) Standard 1X ligase concentration (400,000 U/ml). This is the same gel as in Figure 2, but uncropped. (**B**) Independent repetition of experiments with 1X ligase. (**C**) Ligation with 0.1X diluted ligase concentration (40000 U/mL). (**D**) Ligation with 0.01X diluted ligase concentration (4000 U/ml). Minima in (**D**) are at the same lengths where split bands in (**A**) are found (73, 83, 93, 103), and weak split bands can also be seen in (**D**) at 83 and 93 bp. It is difficult to exactly localize intensity maxima, but they are clearly found between the split bands/minima at ~67, 75-78, 85-90, ~98 bp. This demonstrates that high ligase concentrations do not influence the observed helical repeat. Note that linear and circular dimers are side products from the circularization of the linear template. This dimeric side product could in principle be further reduced by diluting the ligation reactions, but then concentration would become unpractically low for gel analysis. In (**D**), a much higher remaining concentration of ss circles remain due to the lower ligase concentration in the final ligation step. This demonstrates that the formation of ds rings occurs with near-quantitative yields at high ligase concentrations. In summary, we found that decreasing the ligase concentration by 1-2 orders of magnitude does not significantly change the measured helical repeat in our assay.

**Figure S10.**
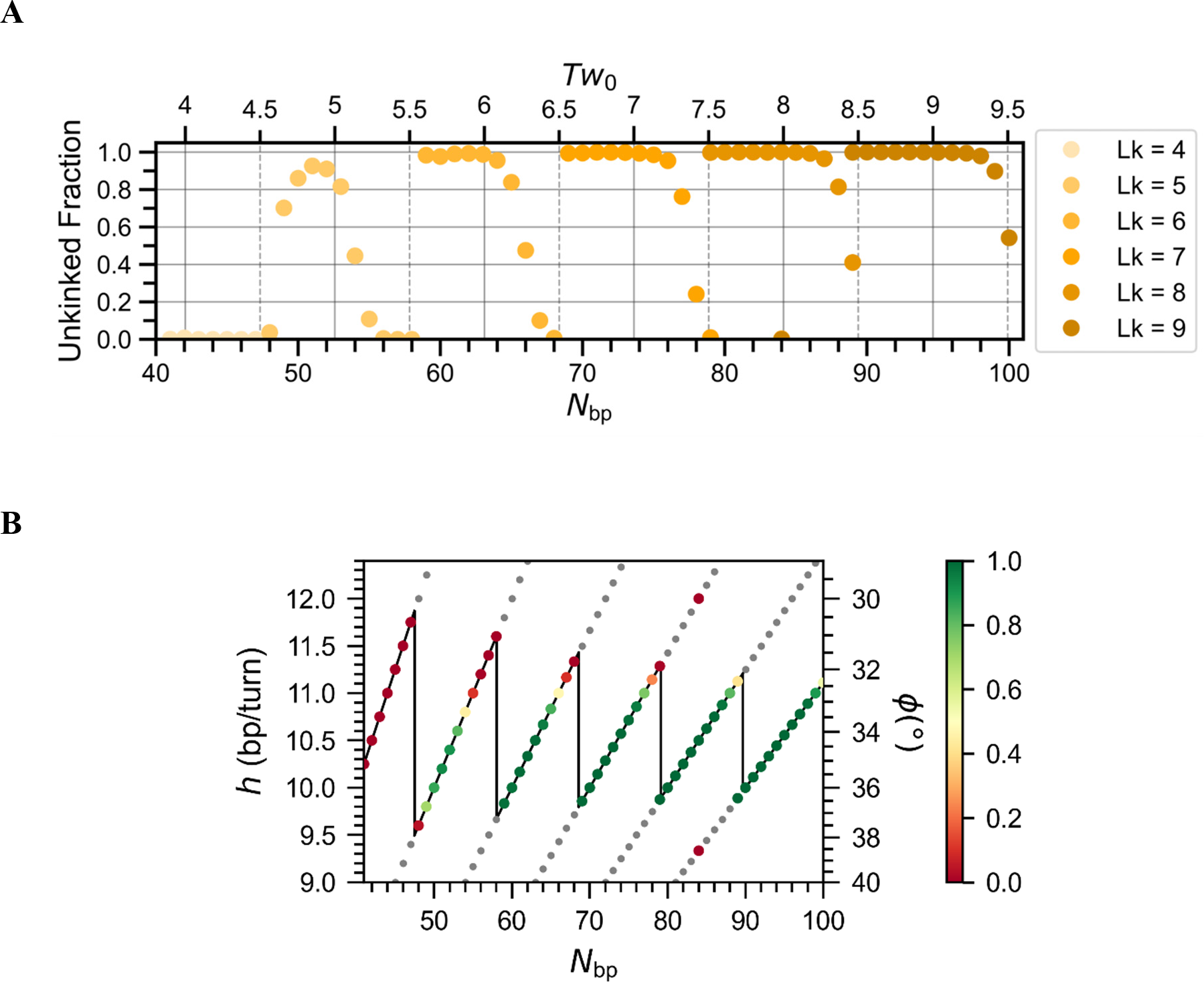
Stability of ligated double-stranded circles in OxDNA simulations. **A**) The fraction of unkinked circles from simulations of circles with N_bp_ = 40 − 100. **B**) The same data, mapped in an h-plot. The color code represents the fraction of intact circles without kinks. Note the similarity with the h-plots in Figure 2 G-H and that overwound circles with a given Lk are stabilized better then the unwound circles. As OxDNA does not show the unwinding due to twist-bend coupling, the N_bp_ in simulations and experiments are not the same.

**Figure S11.**
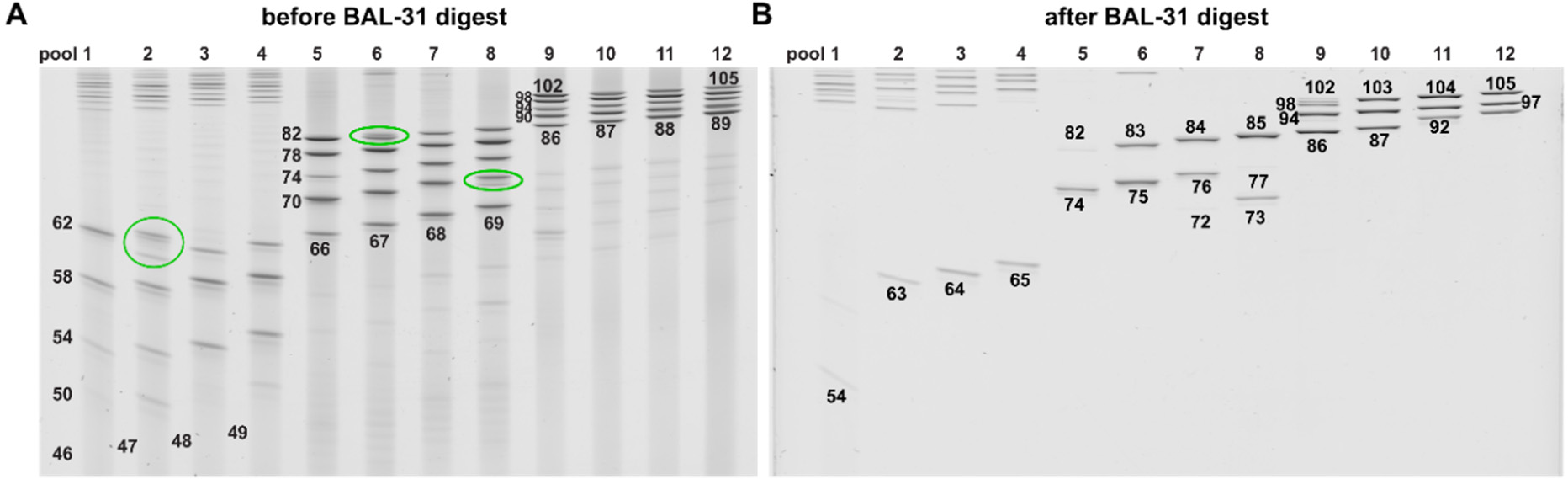
A repeat of an extended nicked circle ligation assay. In these experiments, shorter nicked circles were added to cover the range from N_bp_ = 46 − 105 bp. Right, aliquots of the same ligation reactions were treated with BAL-31 nuclease.

**Figure S12.**
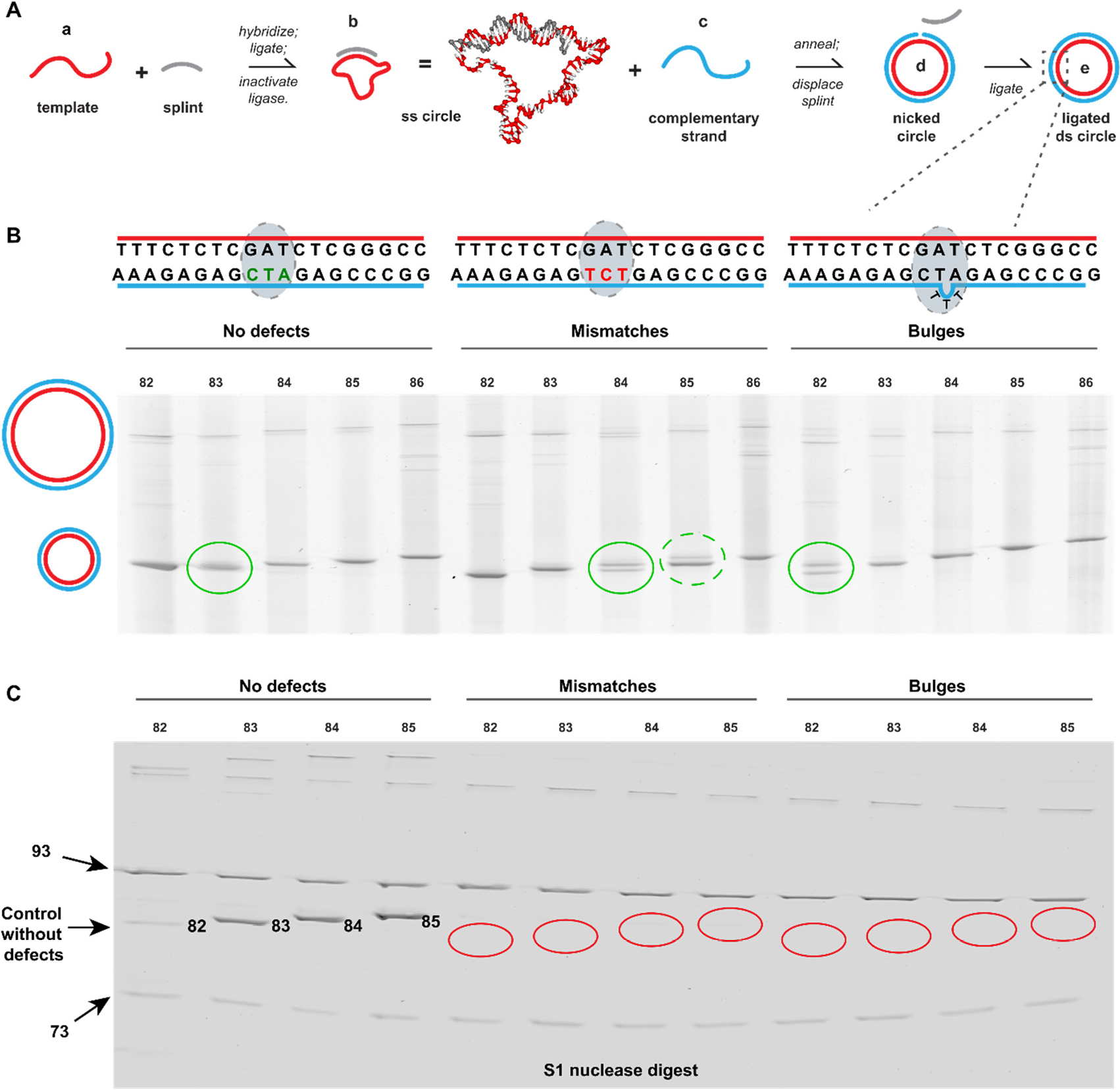
Introduction of errors into the sequences. A) Different versions of circles are generated without defects, with three bp mismatches or a bulge. Mismatches shift topoisomer splitting to larger N_bp_ circles, while bulges shift topoisomer spitting to shorter N_bp_. C) Aliquots of the reactions above were treated with S1 nuclease and all circles with defects are completely digested. As internal controls, nuclease-stable 73 and 93 bp circles that contain no defects were added. Note that the intensity of the control bands is not changed by the nuclease digest indicating error-free sequences.

**Figure S13.**
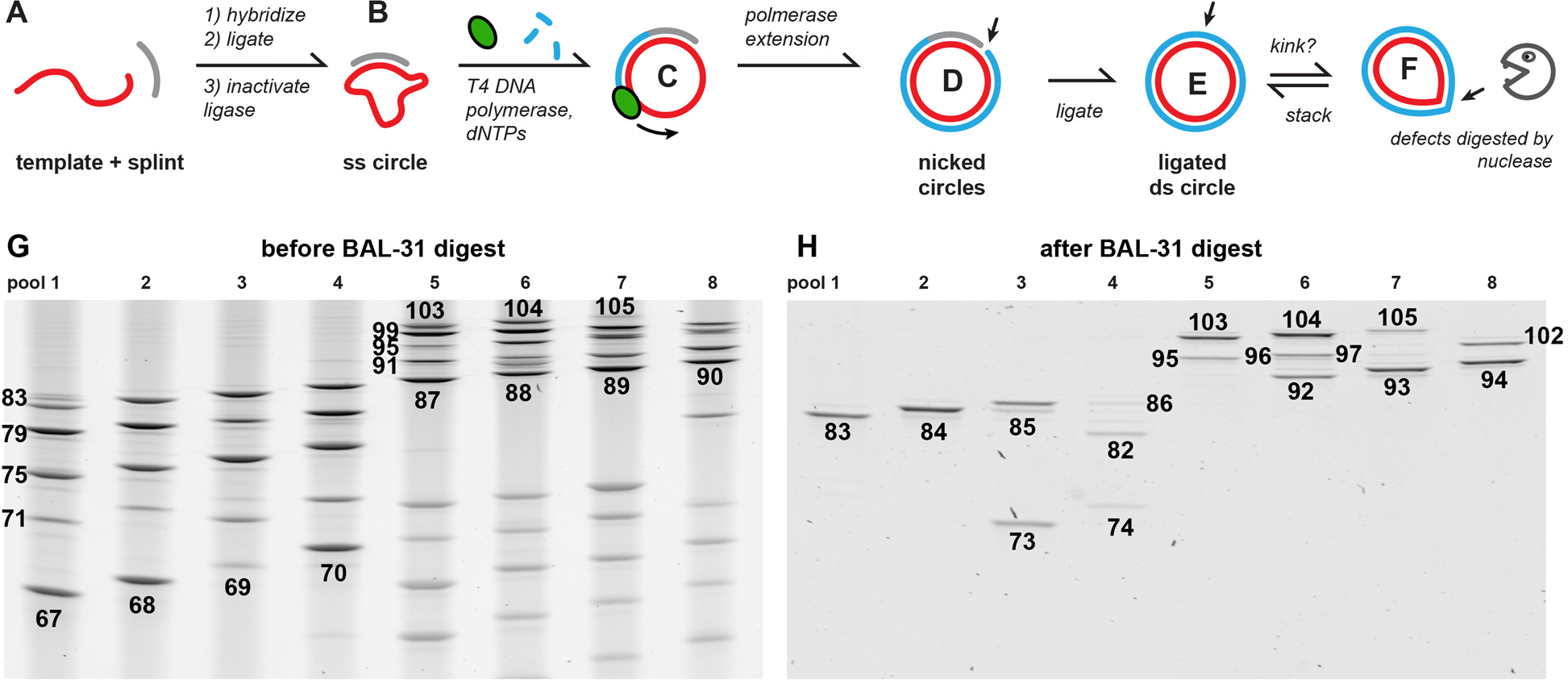
Polymerase extension of splint and ligation. (**A-B**) splint ligation of ss circles is performed as usual. **C**) when adding T4 DNA polymerase and dNTPs, the splint acts as a primer for primer extension. T4 DNA polymerase does not have a strand displacement capability and stops when reaching the other end of the primer, leading to nicked ds circles (**D**). Molecules with eventual errors in the template sequences coming from chemical oligonucleotide synthesis at any position (deletion, insertion, base substitution) would receive a perfectly complementary stand (blue). As a result, all nicked ds circles would be free of defects. These nicked circles can be ligated (**E**) and treated with nuclease as in the standard protocol. Denaturing PAGE gel of the pools after ligation and before (**G**) and after treatment with BAL-31 nuclease (**H**). The yield of ds circles was generally lower than with fully synthetic strands and many single-stranded rings are seen in (**G**) below the ds circles in pools 5-8 indicating an incomplete polymerase extension. However, the location of topoisomer splitting does not change significantly.

**Figure S14.**
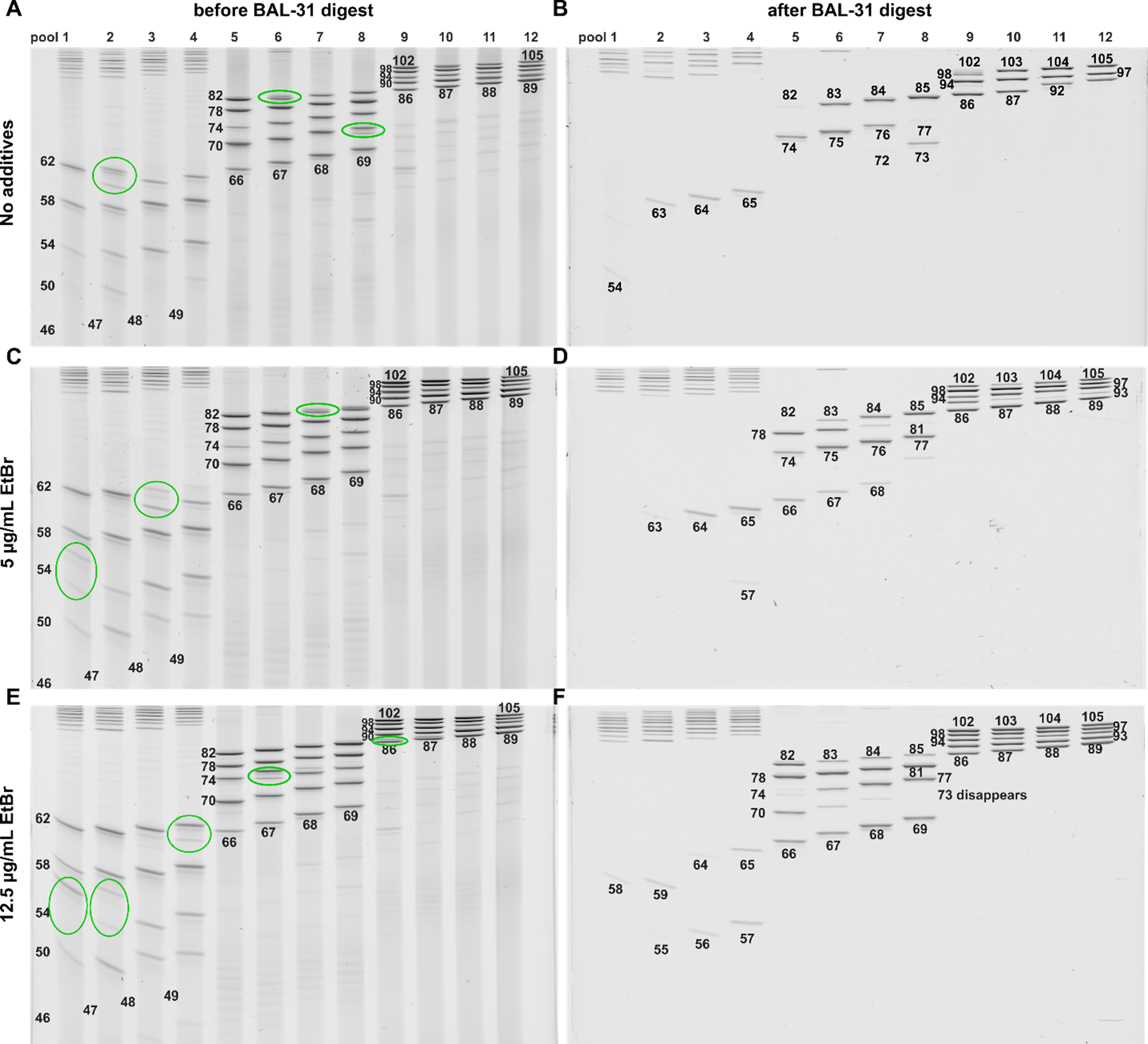
Addition of ethidium bromide. Varying concentrations of ethidium bromide (EtBr) were added to nicked circles before ligation (left gels). EtBr was also present during nuclease treatment (right gels). Note that the topoisomer splitting shifts to longer N_bp_ indicating un unwinding of nicked circles and that EtBr helps stabilizing underwound circles against nuclease digest and that only highly underwound circles are digested. Only the h-plot of the reaction with the highest EtBr concentration is shown in **Figure 6**. Note that the topoisomer splitting is difficult to identify for circles with N_bp_ > 97 bp at high EtBr concentrations as at least one topoisomer of all circles resist nuclease digest.

**Figure S15.**
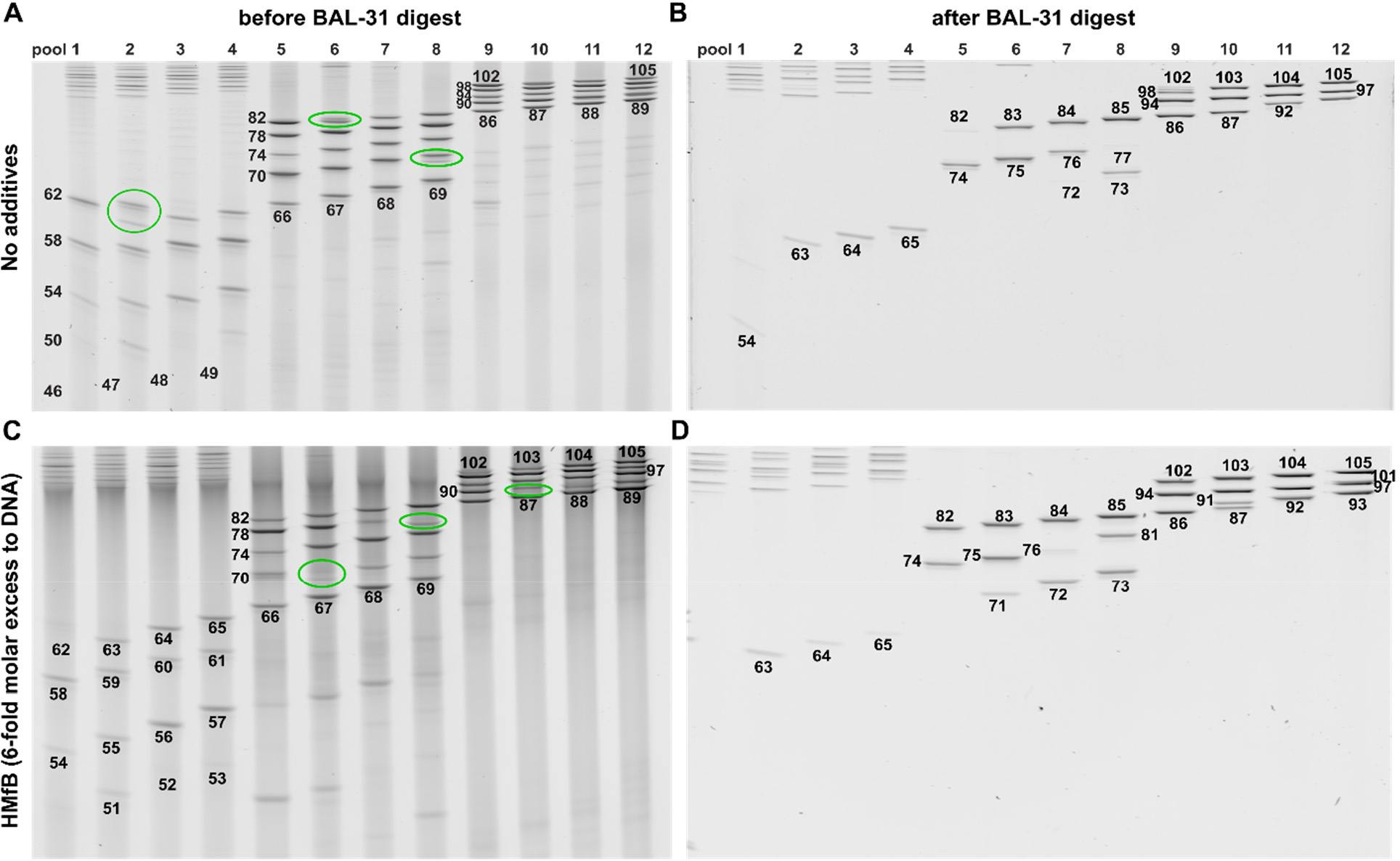
Addition of HMfB. Denaturing PAGE gels of ligation of nicked minicircles in presence of HMfB histone protein. (A-B) without additives; (C-D) after adding the archaeal histone protein HMfB at a 6-fold molar excess over nicked circles. (A/C) Before or (**B/D**) after BAL-31 nuclease treatment. Note that the topoisomer splitting indicated by green circles occurs at a lower N_bp_ indicating an overwinding of the nicked circles by histone proteins. The newly formed bands (lower band = high Lk isomer of 71, 81 bp, 91 bp) are overwound and nuclease stable.

**Figure S16.**
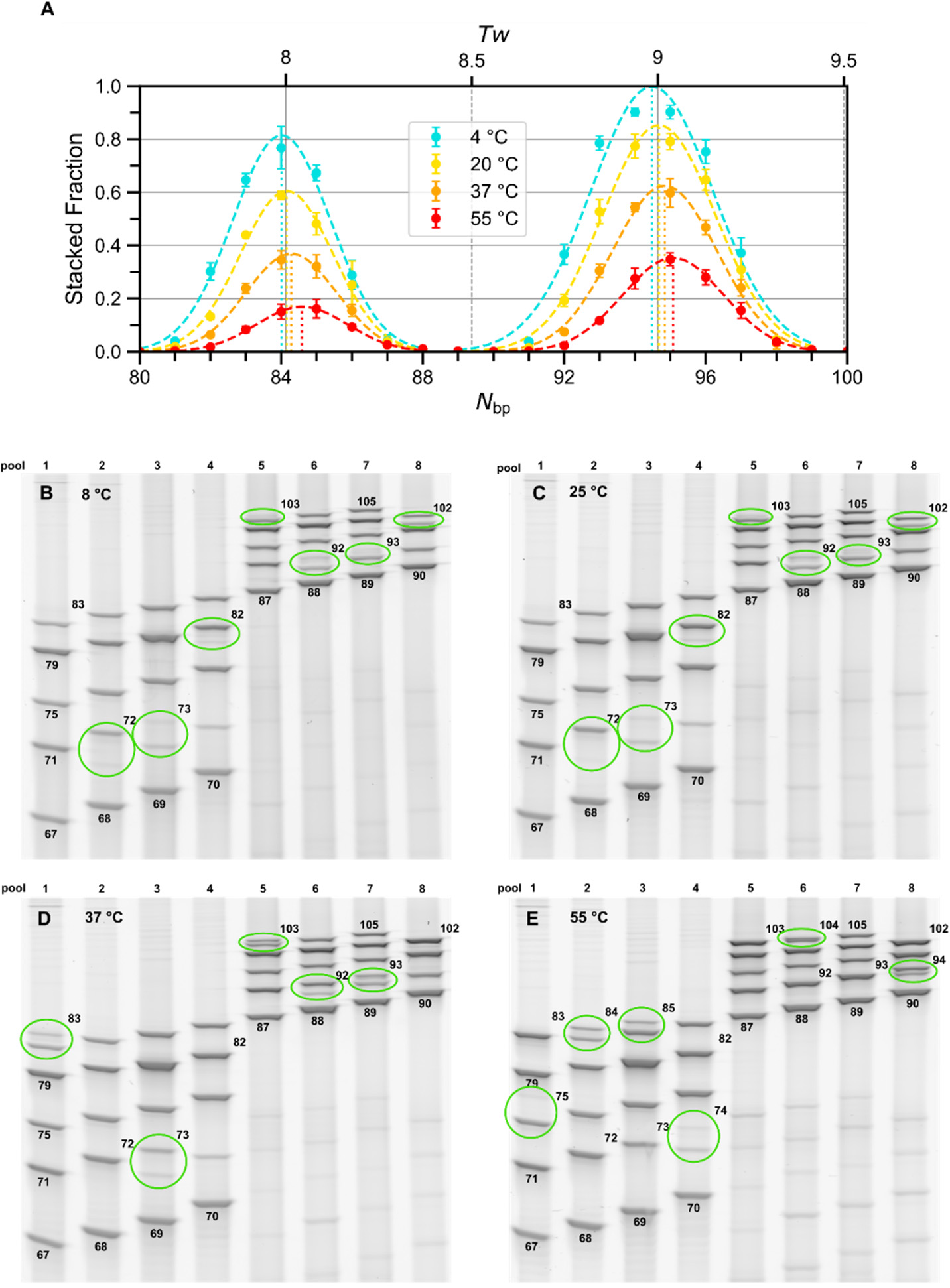
Influence of temperature on the helical repeat in simulated and experimental small circles. (**A**) oxDNA simulations. The fraction of stacked configurations is plotted for 4 sets of simulations, each run at the indicated temperature. Each set contained 21 simulations where sizes were varied from N_bp_ = 80 − 100 bp. The size where stacking is maximal is determined from the maxima of the fitted Gaussians. Increasing the temperature leads to a widening of the helical repeat or unwinding, while lowering the overall probability for stacking. (**B-E**) Denaturing PAGE gels of four sets of experiments in which the final ligation of the nicked ds circle is performed at the indicated temperatures. The ligation time was increased for the reaction at 8 °C due to a lower enzyme activity (8 °C: 12 h; 25 °C: 30 min; 37 °C: 30 min; 55 °C: 20 min). Increasing the temperature shifts topoisomer splitting to longer N_bp_, or changes the relative intensities of split bands, where the upper is the lower Lk isomer.

**Figure S17.**
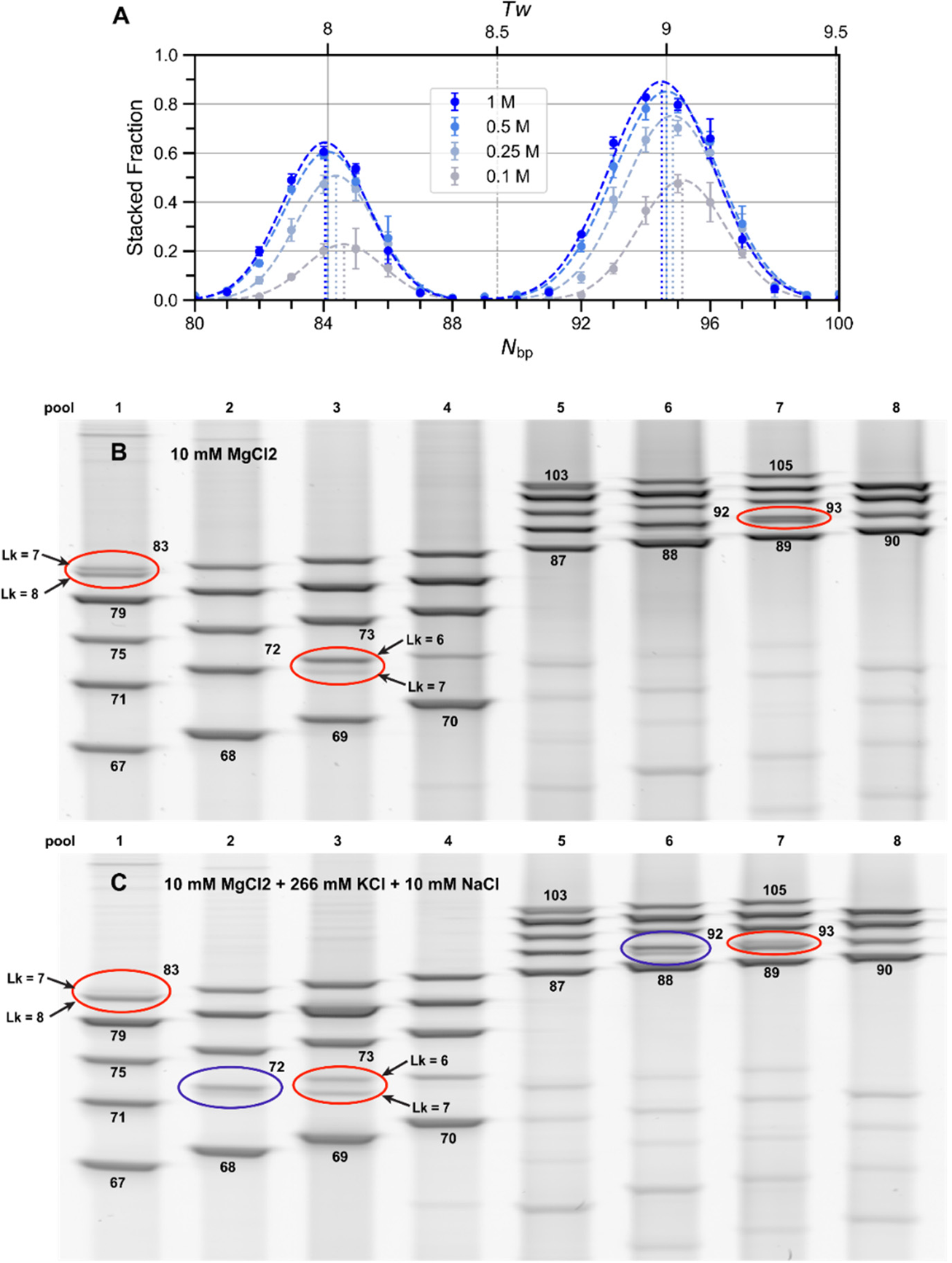
Influence of salt concentration on helical repeat. (**A**) oxDNA simulations and (**B-C**) experiments. **A**) The fraction of stacked configurations in the entire simulation is plotted for 4 different monovalent cation concentrations and for 21 different minicircle lengths ranging from 80-100 bp. Maxima of fitted Gaussians indicate maximum stacking probability for a given salt concentration. As evident from the graph, increasing salinity increases stacking probabilities while shifting the peak to the left indicating overwinding or a decrease of the helical repeat. (**B-C**) Denaturing PAGE gels of ligation experiments at different salt concentrations: (**B**) 10 mM MgCl_2_ with standard T4 DNA ligase and (**C**) 10 mM MgCl_2_ plus an additional 266 mM KCl + 10 mM NaCl with high-salt-T4 DNA ligase to mimic the higher salinity in the nucleus of eukaryotes. Increasing salinity decrease h_0_ or overwinds of DNA as seen by an increase in the proportion of topoisomers with higher linking numbers (lower band of the split band). Red ellipses in (**B**) and (**C**) compare changes in band intensities in split bands with 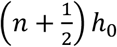. Violet ellipses in (**B**) and (**C**) also indicate the overwinding of DNA in these split bands upon addition of monovalent cations as an increase of the band intensity of the isomer with the higher Lk. These changes are, however, much smaller than the change coming from TBC.

**Figure S18.**
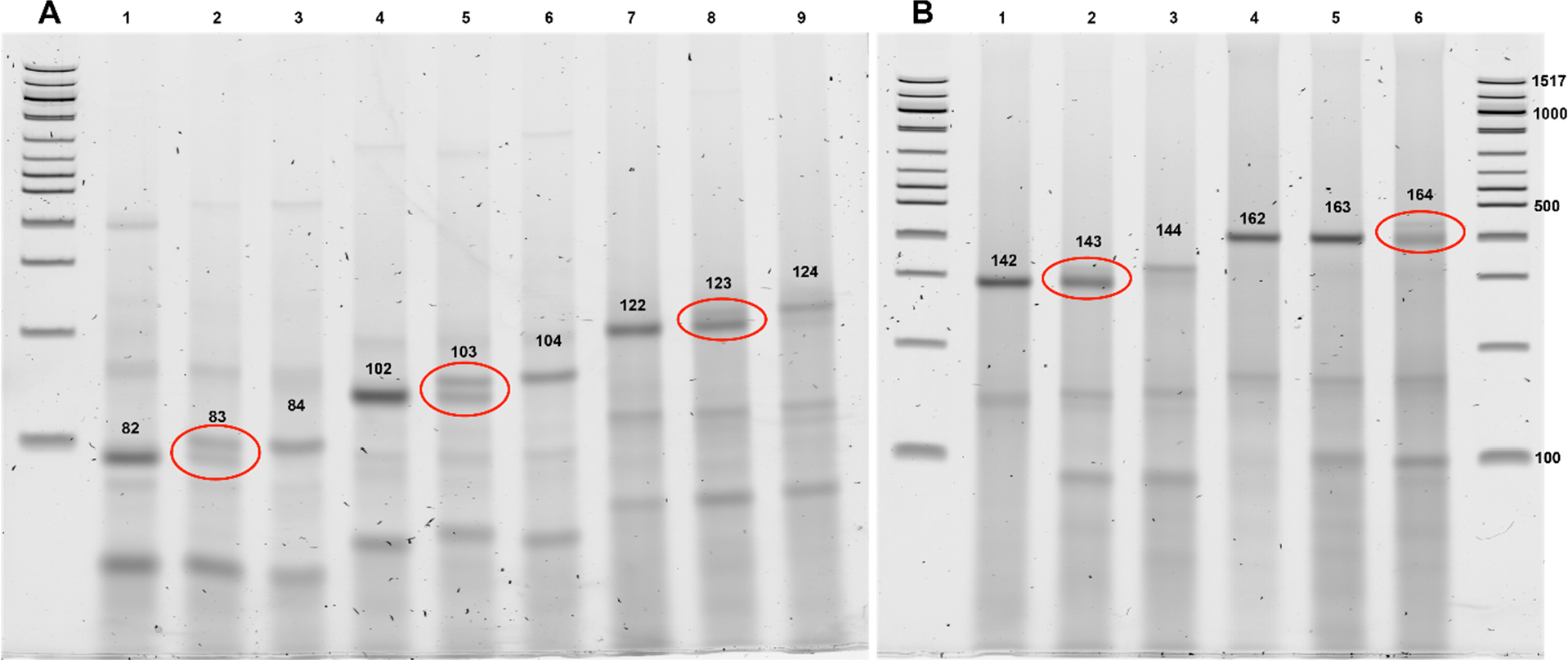
Topoisomer splitting in longer circles. 82-84 and 102-104 bp share the same splint site as the circles shown in Figure 2, but the remainder are other random sequences. Topoisomer splitting is also observed at 123 bp, 143 bp and 164 bp indicating that before ligation these circles had a Tw of 12½, 14½ and 16½ helical turns.

**Figure S19.**
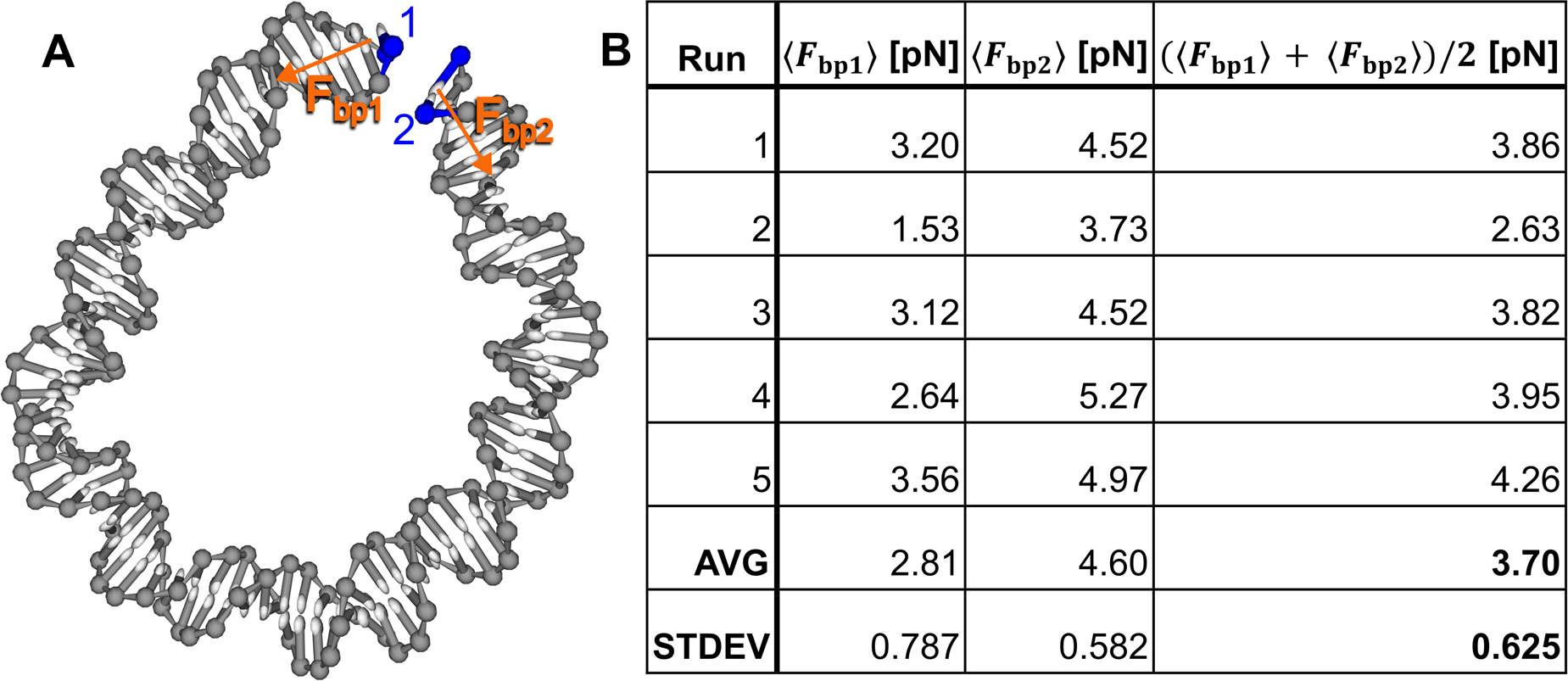
Tension in small minicircles. (**A**) Representative snapshot from an oxDNA simulation that was used to determine the force needed to keep ends of a linear strand double strand in a looped conformation. The initial configuration was created from an 84-mer (h = 10.5 bp/turn with L_k_ = 8), from which we removed three consecutive base pairs to prevent potential stacking. Harmonic traps were placed on both ends (blue) of the duplex to hold them in place. The force of tension on the traps was then measured as described in the Methods section. (**B**) Table of the average forces determined from 5 runs of 10^9^ simulation steps (100,000 snapshots). Note that the tension fluctuations are very high throughout the simulation due to Brownian motion. Forces are about 21% lower in these simulations compared to theoretical calculations, because the traps allow for a relaxation or the circle into a teardrop-shape, which reduces the tension compared to the circular conformation in the theoretical calculation. The distortion into a teardrop shape in the simulation is also the reason why the vectors in (**A**) do not point in exactly opposite directions. Importantly, the force of tension is too low for a significant influence of twist-strech coupling to h_0_.

**Table S1.**
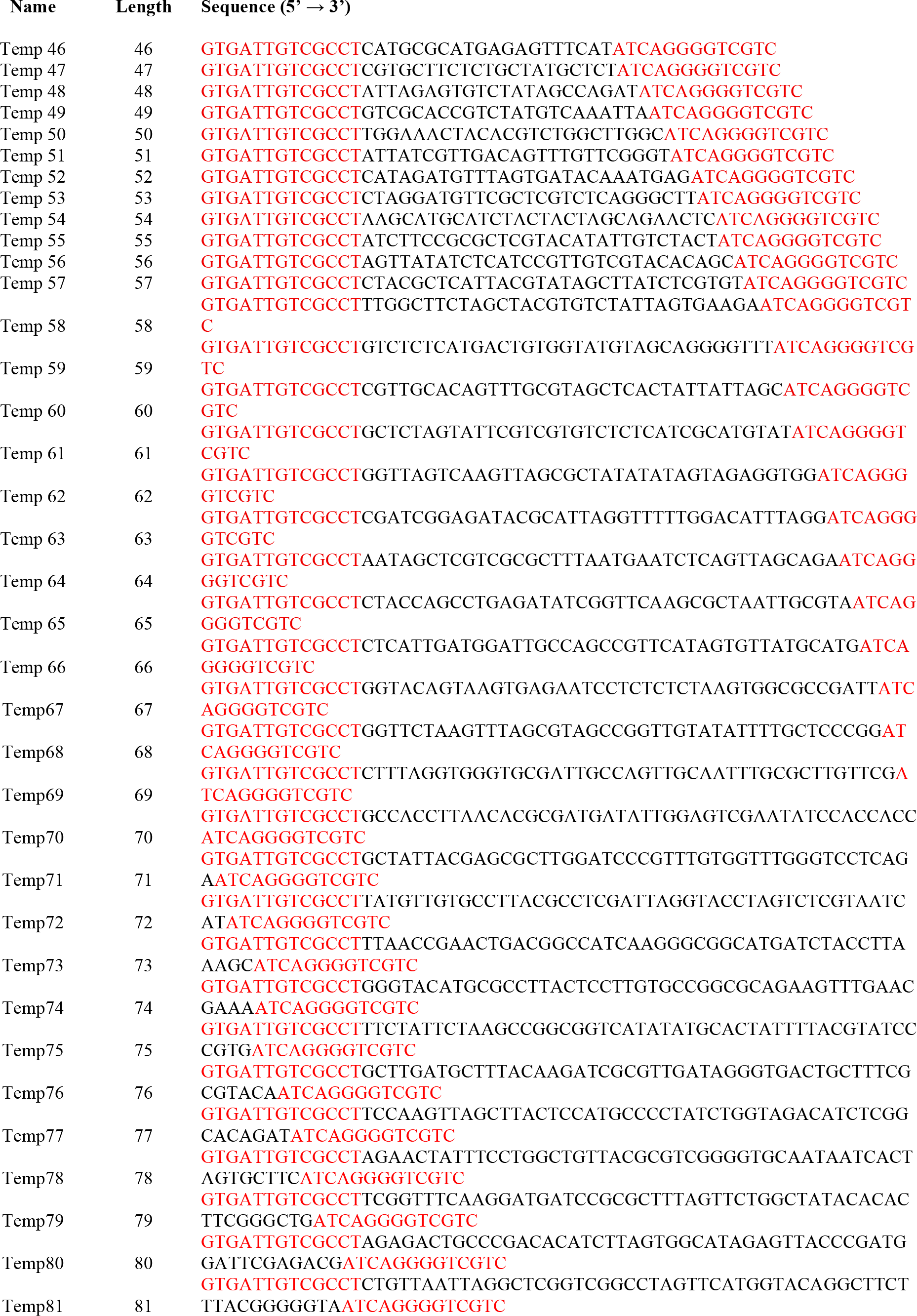
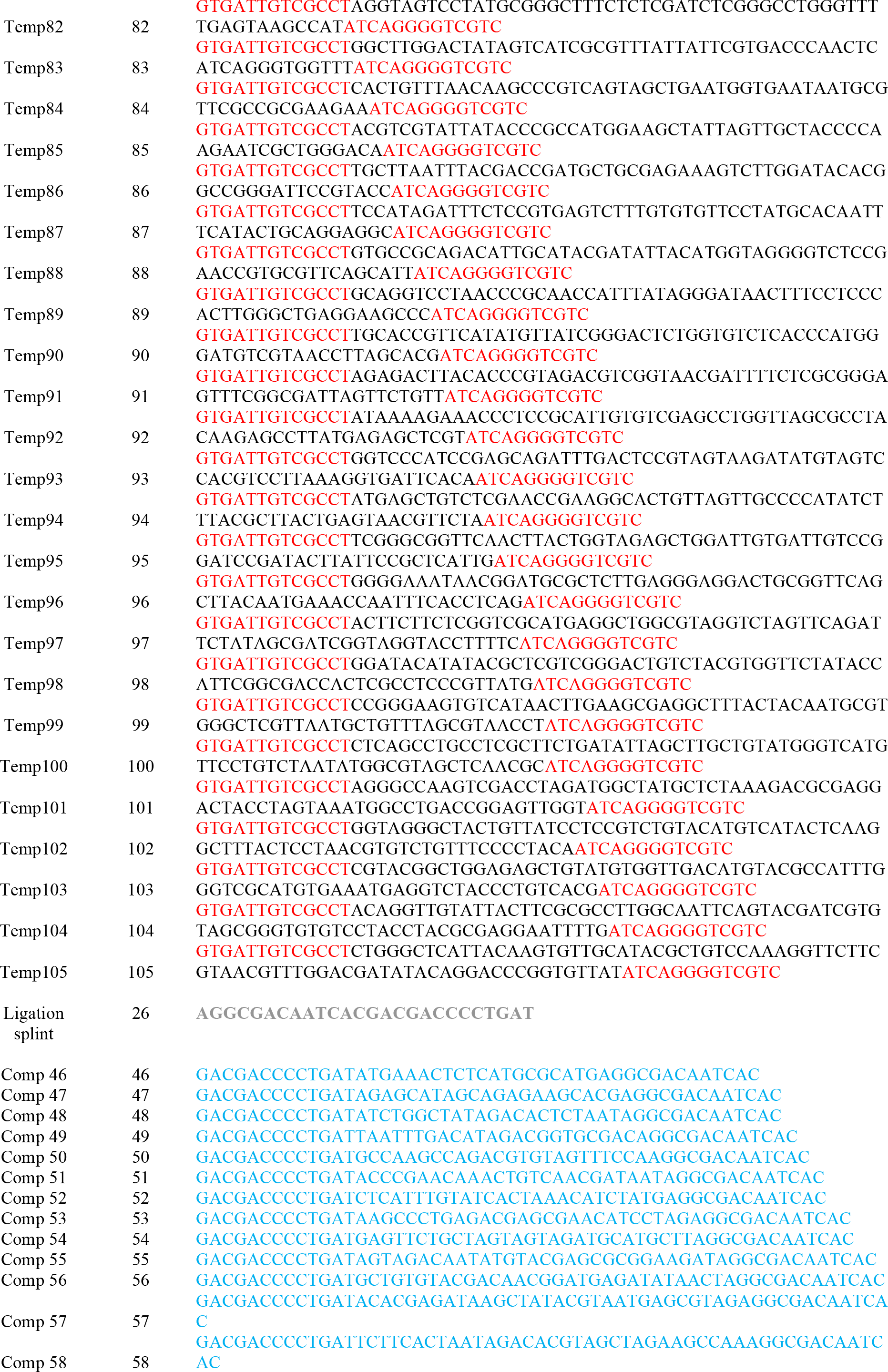
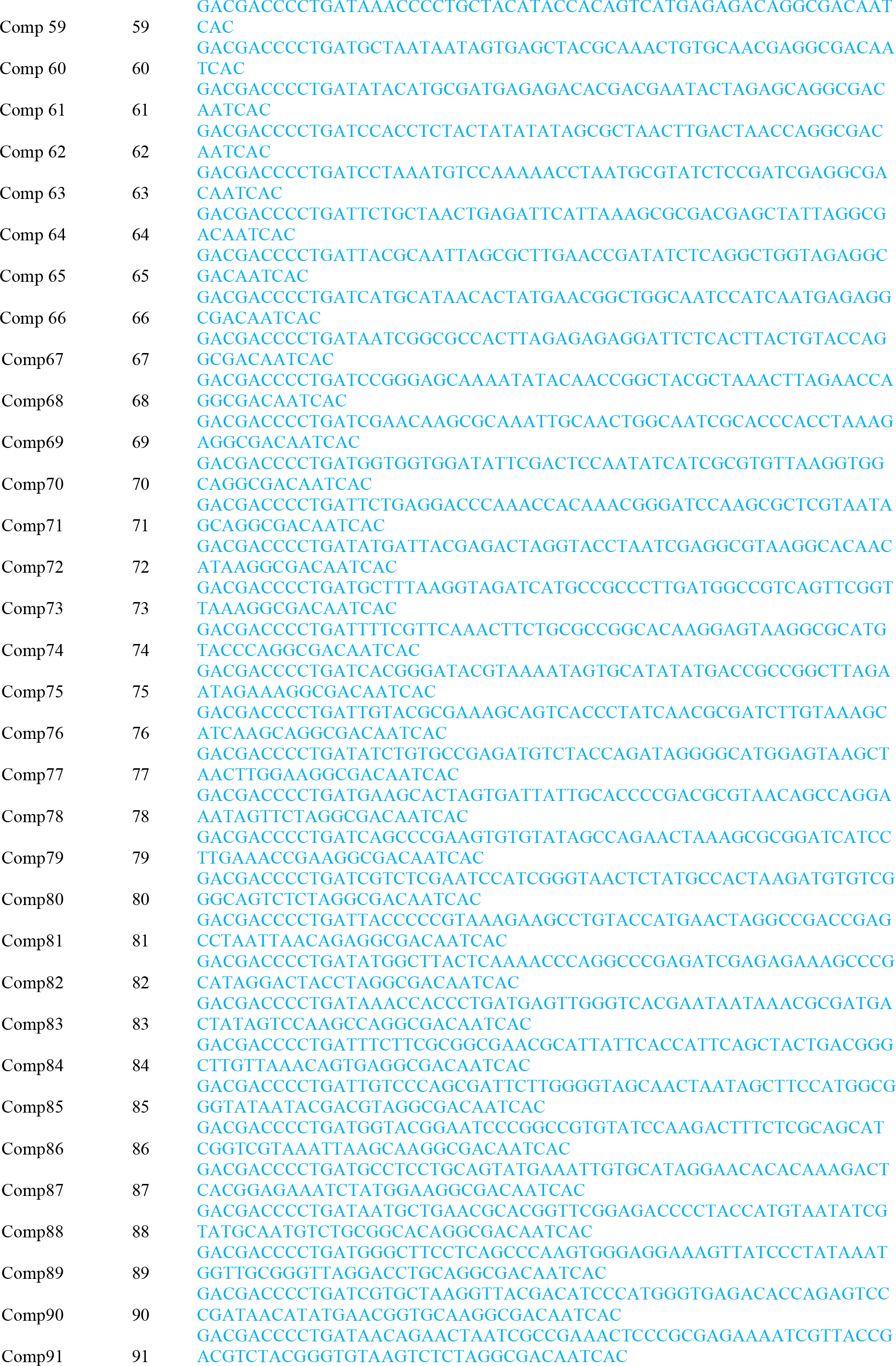
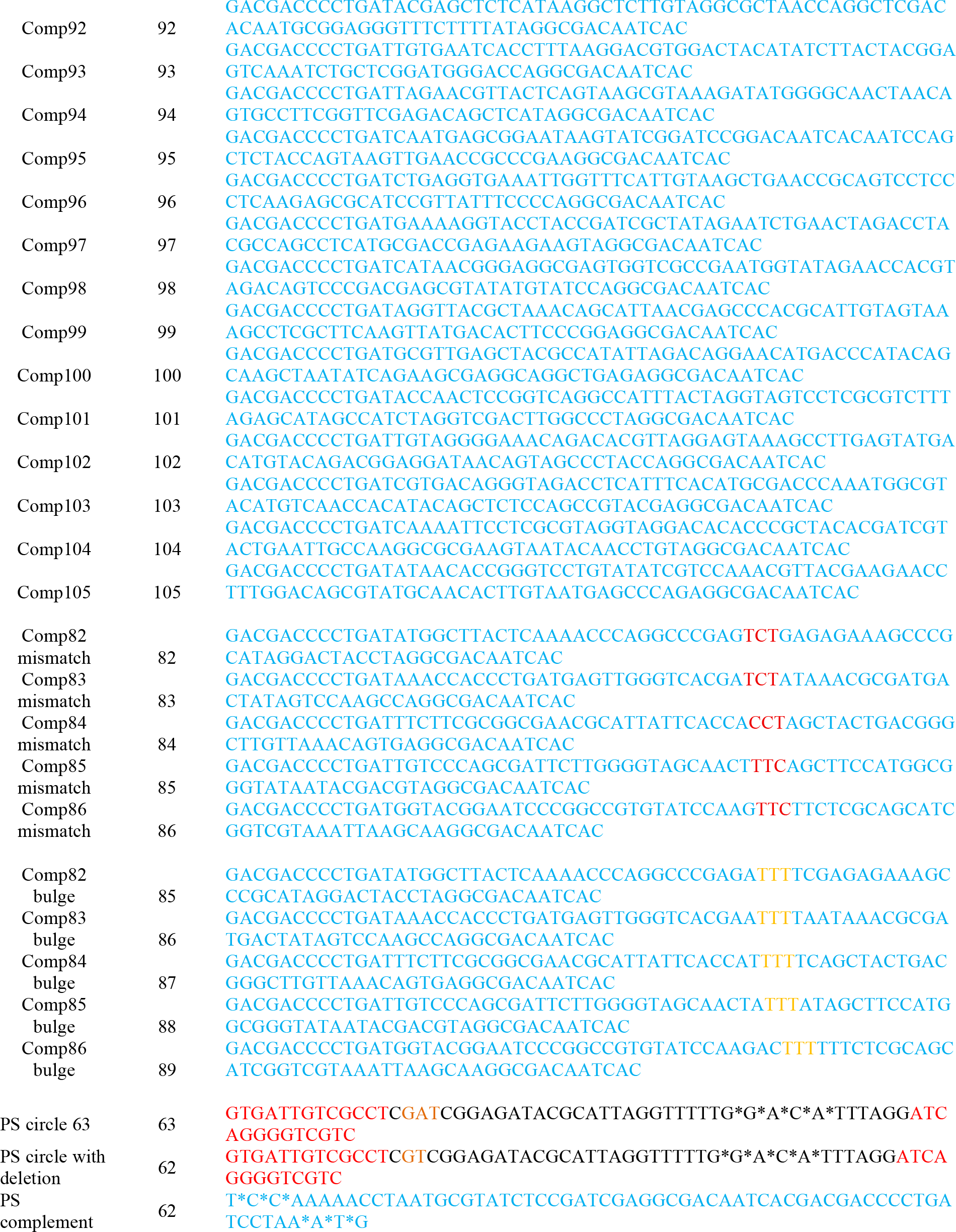
The sequences of all oligonucleotides used in this study. Highlighted in red: common, complementary sequences for ligation splint.

## Supplementary movies

| Movie # | Model | $N_{bp}$ | $Tw$ (start) | $\Delta Tw$ (start) | Nicked? | Sim. Time | Comments |
| --- | --- | --- | --- | --- | --- | --- | --- |
| S01 | oxDNA | 84 | 8 | 0 | Yes | 55 ns | Coarse time steps. Multiple stacking and kinking events, OxDNA time |
| S02 | oxDNA | 84 | 8 | 0 | Yes | 30 ns | Slow-motion of unstacking and stacking events |
| S03 | oxDNA | 89 | 8.5 | 0 | Yes | 55 ns | Coarse time steps, no stacking observed |
| S04 | oxDNA | 89 | 8 | 0.5 | Yes | 5 ns | Slow-motion of unstacking of an initially stacked and torsionally twisted circle |
| S05 | OL21 | 84 | 9 | -1 | Yes | 50 ns | Stack at the start of simulation |
| S06 | OL21 | 84 | 7 | 1 | Yes | 50 ns | Stack at the start of simulation |
| S07 | OL21 | 84 | 7 | -1 | Ligated | 50 ns | Highly underwound and unstable DNA structure |
| S08 | OL21 | 84 | 8 | 0 | Ligated | 50 ns | Torsionally relaxed, stable |
| S09 | OL21 | 80 | 8 | +0.4 | Yes | 50 ns | With HMfB proteins |
| S10 | OL21 | 80 | 8 | +0.4 | Yes | 50 ns | Same as above, but HMfB removed before start of simulation. |

## References

1. Basu, A., Bobrovnikov, D. G. & Ha, T. DNA mechanics and its biological impact. Journal of Molecular Biology 433, 166861 (2021).

2. Garcia, H. G., Grayson, P., Han, L., Inamdar, M., Kondev, J., Nelson, P. C., Phillips, R., Widom, J. & Wiggins, P. A. Biological consequences of tightly bent DNA: The other life of a macromolecular celebrity. Biopolymers 85, 115–130 (2007).

3. Crick, F. H. C. & Klug, A. Kinky Helix. Nature 255, 530–533 (1975).

4. Kahn, J. D., Yun, E. & Crothers, D. M. Detection of localized DNA flexibility. Nature 368, 163–166 (1994).

5. Cloutier, T. E. & Widom, J. Spontaneous Sharp Bending of Double-Stranded DNA. Molecular Cell 14, 355–362 (2004).

6. Cloutier, T. E. & Widom, J. DNA twisting flexibility and the formation of sharply looped protein–DNA complexes. Proceedings of the National Academy of Sciences 102, 3645–3650 (2005).

7. Wiggins, P. A., van der Heijden, T., Moreno-Herrero, F., Spakowitz, A., Phillips, R., Widom, J., Dekker, C. & Nelson, P. C. High flexibility of DNA on short length scales probed by atomic force microscopy. Nat. Nanotechnol. 1, 137–141 (2006).

8. Du, Q., Smith, C., Shiffeldrim, N., Vologodskaia, M. & Vologodskii, A. Cyclization of short DNA fragments and bending fluctuations of the double helix. Proceedings of the National Academy of Sciences 102, 5397–5402 (2005).

9. Mathew-Fenn, R. S., Das, R. & Harbury, P. A. B. Remeasuring the Double Helix. Science 322, 446–449 (2008).

10. Mastroianni, A. J., Sivak, D. A., Geissler, P. L. & Alivisatos, A. P. Probing the Conformational Distributions of Subpersistence Length DNA. Biophysical Journal 97, 1408–1417 (2009).

11. Demurtas, D., Amzallag, A., Rawdon, E. J., Maddocks, J. H., Dubochet, J. & Stasiak, A. Bending modes of DNA directly addressed by cryo-electron microscopy of DNA minicircles. Nucl. Acids Res. (2009).

12. Forties, R. A., Bundschuh, R. & Poirier, M. G. The flexibility of locally melted DNA. Nucleic Acids Research 37, 4580–4586 (2009).

13. Qu, H., Wang, Y., Tseng, C.-Y. & Zocchi, G. Critical Torque for Kink Formation in Double-Stranded DNA. Phys. Rev. X 1, 021008 (2011).

14. Vafabakhsh, R. & Ha, T. Extreme Bendability of DNA Less than 100 Base Pairs Long Revealed by Single-Molecule Cyclization. Science 337, 1097–1101 (2012).

15. Vologodskii, A., Du, Q. & Frank-Kamenetskii, M. D. Bending of short DNA helices. Artificial DNA: PNA & XNA 4, 1–3 (2013).

16. Vologodskii, A. & D. Frank-Kamenetskii, M. Strong bending of the DNA double helix. Nucleic Acids Res 41, 6785–6792 (2013).

17. Fields, A. P., Meyer, E. A. & Cohen, A. E. Euler buckling and nonlinear kinking of double-stranded DNA. Nucleic Acids Research 41, 9881–9890 (2013).

18. Mazur, A. K. & Maaloum, M. DNA Flexibility on Short Length Scales Probed by Atomic Force Microscopy. Phys. Rev. Lett. 112, 068104 (2014).

19. Le, T. T. & Kim, H. D. Probing the elastic limit of DNA bending. Nucleic Acids Research 42, 10786–10794 (2014).

20. Harrison, R. M., Romano, F., Ouldridge, T. E., Louis, A. A. & Doye, J. P. K. Identifying Physical Causes of Apparent Enhanced Cyclization of Short DNA Molecules with a Coarse-Grained Model. J. Chem. Theory Comput. 15, 4660–4672 (2019).

21. Jeong, J. & Kim, H. D. Determinants of cyclization–decyclization kinetics of short DNA with sticky ends. Nucleic Acids Res 48, 5147–5156 (2020).

22. Yeou, S., Hwang, J., Yi, J., Kim, C., Keun Kim, S. & Ki Lee, N. Cytosine methylation regulates DNA bendability depending on the curvature. Chemical Science 13, 7516–7525 (2022).

23. Shimada, J. & Yamakawa, H. Ring-closure probabilities for twisted wormlike chains. Application to DNA. Macromolecules 17, 689–698 (1984).

24. Yan, J. & Marko, J. F. Localized Single-Stranded Bubble Mechanism for Cyclization of Short Double Helix DNA. Phys. Rev. Lett. 93, 108108 (2004).

25. Lankaš, F., Lavery, R. & Maddocks, J. H. Kinking Occurs during Molecular Dynamics Simulations of Small DNA Minicircles. Structure 14, 1527–1534 (2006).

26. Harris, S. A., Laughton, C. A. & Liverpool, T. B. Mapping the phase diagram of the writhe of DNA nanocircles using atomistic molecular dynamics simulations. Nucleic Acids Research 36, 21–29 (2008).

27. Mitchell, J. S., Laughton, C. A. & Harris, S. A. Atomistic simulations reveal bubbles, kinks and wrinkles in supercoiled DNA. Nucleic Acids Research 39, 3928–3938 (2011).

28. Irobalieva, R. N., Fogg, J. M., Catanese, D. J., Sutthibutpong, T., Chen, M., Barker, A. K., Ludtke, S. J., Harris, S. A., Schmid, M. F., Chiu, W. & Zechiedrich, L. Structural diversity of supercoiled DNA. Nat Commun 6, 8440 (2015).

29. Shore, D., Langowski, J. & Baldwin, R. L. DNA flexibility studied by covalent closure of short fragments into circles. Proceedings of the National Academy of Sciences 78, 4833–4837 (1981).

30. Shore, D. & Baldwin, R. L. Energetics of DNA twisting: II. Topoisomer analysis. Journal of Molecular Biology 170, 983–1007 (1983).

31. Horowitz, D. S. & Wang, J. C. Torsional rigidity of DNA and length dependence of the free energy of DNA supercoiling. J Mol Biol 173, 75–91 (1984).

32. Taylor, W. H. & Hagerman, P. J. Application of the method of phage T4 DNA ligase-catalyzed ring-closure to the study of DNA structure: II. NaCl-dependence of DNA flexibility and helical repeat. Journal of Molecular Biology 212, 363–376 (1990).

33. Spakowitz, A. J. Wormlike chain statistics with twist and fixed ends. EPL 73, 684 (2006).

34. Wang, J. C. Helical repeat of DNA in solution. Proceedings of the National Academy of Sciences 76, 200–203 (1979).

35. Hayes, J. J., Tullius, T. D. & Wolffe, A. P. The structure of DNA in a nucleosome. Proceedings of the National Academy of Sciences 87, 7405–7409 (1990).

36. Marko, J. F. & Siggia, E. D. Bending and twisting elasticity of DNA. Macromolecules 27, 981–988 (1994).

37. Du, Q., Kotlyar, A. & Vologodskii, A. Kinking the double helix by bending deformation. Nucleic Acids Res. 36, 1120 (2008).

38. Protozanova, E., Yakovchuk, P. & Frank-Kamenetskii, M. D. Stacked-Unstacked Equilibrium at the Nick Site of DNA. Journal of Molecular Biology 342, 775–785 (2004).

39. Snodin, B. E. K., Romano, F., Rovigatti, L., Ouldridge, T. E., Louis, A. A. & Doye, J. P. K. Direct Simulation of the Self-Assembly of a Small DNA Origami. ACS Nano 10, 1724–1737 (2016).

40. Sengar, A., Ouldridge, T. E., Henrich, O., Rovigatti, L. & Sulc, P. A primer on the oxDNA model of DNA: When to use it, how to simulate it and how to interpret the results. Front. Mol. Biosci. 8, 693710 (2021).

41. Agarwal, N. P., Matthies, M., Joffroy, B. & Schmidt, T. L. Structural Transformation of Wireframe DNA Origami via DNA Polymerase Assisted Gap-Filling. ACS Nano 12, 2546–2553 (2018).

42. Smith, S., Finzi, L. & Bustamante, C. Direct mechanical measurements of the elasticity of single DNA molecules by using magnetic beads. Science 258, 1122–1126 (1992).

43. Bustamante, C., Marko, J. F., Siggia, E. D. & Smith, S. Entropic Elasticity of λ-Phage DNA. Science 265, 1599–1600 (1994).

44. Strick, T. R., Allemand, J.-F., Bensimon, D., Bensimon, A. & Croquette, V. The Elasticity of a Single Supercoiled DNA Molecule. Science 271, 1835–1837 (1996).

45. Gore, J., Bryant, Z., Nöllmann, M., Le, M. U., Cozzarelli, N. R. & Bustamante, C. DNA overwinds when stretched. Nature 442, 836–839 (2006).

46. Pascal, J. M., O’Brien, P. J., Tomkinson, A. E. & Ellenberger, T. Human DNA ligase I completely encircles and partially unwinds nicked DNA. Nature 432, 473–478 (2004).

47. Shi, K., Bohl, T. E., Park, J., Zasada, A., Malik, S., Banerjee, S., Tran, V., Li, N., Yin, Z., Kurniawan, F., Orellana, K. & Aihara, H. T4 DNA ligase structure reveals a prototypical ATP-dependent ligase with a unique mode of sliding clamp interaction. Nucleic Acids Research 46, 10474–10488 (2018).

48. Yuan, C., Lou, X. W., Rhoades, E., Chen, H. & Archer, L. A. T4 DNA ligase is more than an effective trap of cyclized dsDNA. Nucleic Acids Research 35, 5294–5302 (2007).

49. Mattiroli, F., Bhattacharyya, S., Dyer, P. N., White, A. E., Sandman, K., Burkhart, B. W., Byrne, K. R., Lee, T., Ahn, N. G., Santangelo, T. J., Reeve, J. N. & Luger, K. Structure of histone-based chromatin in Archaea. Science 357, 609–612 (2017).

50. Kriegel, F., Matek, C., Dršata, T., Kulenkampff, K., Tschirpke, S., Zacharias, M., Lankaš, F. & Lipfert, J. The temperature dependence of the helical twist of DNA. Nucleic Acids Research 46, 7998–8009 (2018).

51. Bates, A. D. & Maxwell, A. DNA gyrase can supercoil DNA circles as small as 174 base pairs. The EMBO Journal 8, 1861–1866 (1989).

52. Skoruppa, E., Laleman, M., Nomidis, S. K. & Carlon, E. DNA elasticity from coarse-grained simulations: The effect of groove asymmetry. J. Chem. Phys. 146, 214902 (2017).

53. Nomidis, S. K., Skoruppa, E., Carlon, E. & Marko, J. F. Twist-bend coupling and the statistical mechanics of the twistable wormlike-chain model of DNA: Perturbation theory and beyond. Phys. Rev. E 99, 032414 (2019).

54. Mou, X., Liu, K., He, L. & Li, S. Mechanical response of double-stranded DNA: Bend, twist, and overwind. The Journal of Chemical Physics 161, 085102 (2024).

55. Skoruppa, E., Nomidis, S. K., Marko, J. F. & Carlon, E. Bend-Induced Twist Waves and the Structure of Nucleosomal DNA. Phys. Rev. Lett. 121, 088101 (2018).

56. Müller, J., Oehler, S. & Müller-Hill, B. Repression oflacPromoter as a Function of Distance, Phase and Quality of an AuxiliarylacOperator. Journal of Molecular Biology 257, 21–29 (1996).

57. Lee, D. H. & Schleif, R. F. In vivo DNA loops in araCBAD: size limits and helical repeat. Proc. Natl. Acad. Sci. U.S.A. 86, 476–480 (1989).

58. Haykinson, M. J. & Johnson, R. C. DNA looping and the helical repeat in vitro and in vivo: effect of HU protein and enhancer location on Hin invertasome assembly. The EMBO Journal 12, 2503–2512 (1993).

59. Li, G. & Widom, J. Nucleosomes facilitate their own invasion. Nat Struct Mol Biol 11, 763–769 (2004).

60. Spakowitz, A. J. & Wang, Z.-G. DNA Packaging in Bacteriophage: Is Twist Important? Biophysical Journal 88, 3912–3923 (2005).

61. Schmidt, T. L. & Heckel, A. Construction of a Structurally Defined Double-Stranded DNA Catenane. Nano Lett. 11, 1739–1742 (2011).

62. Ackermann, D., Schmidt, T. L., Hannam, J. S., Purohit, C. S., Heckel, A. & Famulok, M. A double-stranded DNA rotaxane. Nat. Nanotechnol. 5, 436–442 (2010).

63. Iric, K., Subramanian, M., Oertel, J., Agarwal, N. P., Matthies, M., Periole, X., Sakmar, T. P., Huber, T., Fahmy, K. & Schmidt, T. L. DNA-encircled lipid bilayers. Nanoscale 10, 18463–18467 (2018).

64. Dietz, H., Douglas, S. M. & Shih, W. M. Folding DNA into Twisted and Curved Nanoscale Shapes. Science 325, 725–730 (2009).

65. Han, D., Pal, S., Nangreave, J., Deng, Z., Liu, Y. & Yan, H. DNA Origami with Complex Curvatures in Three-Dimensional Space. Science 332, 342–346 (2011).

66. Joffroy, B., Uca, Y. O., Prešern, D., Doye, J. P. K. & Schmidt, T. L. Rolling circle amplification shows a sinusoidal template length-dependent amplification bias. Nucleic Acids Res 46, 538–545 (2018).

67. Schindelin, J., Arganda-Carreras, I., Frise, E., Kaynig, V., Longair, M., Pietzsch, T., Preibisch, S., Rueden, C., Saalfeld, S., Schmid, B., Tinevez, J.-Y., White, D. J., Hartenstein, V., Eliceiri, K., Tomancak, P. & Cardona, A. Fiji: an open-source platform for biological-image analysis. Nat Methods 9, 676–682 (2012).

68. Poppleton, E., Bohlin, J., Matthies, M., Sharma, S., Zhang, F. & Šulc, P. Design, optimization and analysis of large DNA and RNA nanostructures through interactive visualization, editing and molecular simulation. Nucleic Acids Research 48, e72 (2020).

69. Bohlin, J., Matthies, M., Poppleton, E., Procyk, J., Mallya, A., Yan, H. & Šulc, P. Design and simulation of DNA, RNA and hybrid protein–nucleic acid nanostructures with oxView. Nat Protoc 17, 1762–1788 (2022).

70. Walther, J. & Orozco, M. MC_DNA: A Web Server for the Detailed Study of the Structure and Dynamics of DNA and Chromatin Fibers.

71. BIOVIA, Dassault Systèmes, Visualizer. San Diego: Dassault Systèmes, 2021. Dassault Systèmes.

72. Hess, B., Bekker, H., Berendsen, H. J. & Fraaije, J. G. LINCS: A linear constraint solver for molecular simulations. Journal of computational chemistry 18, 1463–1472 (1997).

73. Darden, T., York, D. & Pedersen, L. Particle mesh Ewald: An N⋅ log (N) method for Ewald sums in large systems. The Journal of chemical physics 98, 10089–10092 (1993).

74. Bussi, G., Donadio, D. & Parrinello, M. Canonical sampling through velocity rescaling. The Journal of chemical physics 126, (2007).

75. Abraham, M., Alekseenko, A., Basov, V., Bergh, C., Briand, E., Brown, A., Doijade, M., Fiorin, G., Fleischmann, S., Gorelov, S., Gouaillardet, G., Gray, A., Irrgang, M. E., Jalalypour, F., Jordan, J., Kutzner, C., Lemkul, J. A., Lundborg, M., Merz, P., Miletic, V., Morozov, D., Nabet, J., Pall, S., Pasquadibisceglie, A., Pellegrino, M., Santuz, H., Schulz, R., Shugaeva, T., Shvetsov, A., Villa, A., Wingbermuehle, S., Hess, B. & Lindahl, E. GROMACS 2024.4 Manual. https://doi.org/10.5281/zenodo.14016613 (2024) doi:10.5281/zenodo.14016613.

76. Pérez, A., Marchán, I., Svozil, D., Sponer, J., Cheatham, T. E., Laughton, C. A. & Orozco, M. Refinement of the AMBER force field for nucleic acids: improving the description of α/γ conformers. Biophysical journal 92, 3817–3829 (2007).

77. Cornell, W. D., Cieplak, P., Bayly, C. I., Gould, I. R., Merz, K. M., Ferguson, D. M., Spellmeyer, D. C., Fox, T., Caldwell, J. W. & Kollman, P. A. A second generation force field for the simulation of proteins, nucleic acids, and organic molecules. Journal of the American Chemical Society 117, 5179–5197 (1995).

78. Yoo, J. & Aksimentiev, A. Improved parameterization of amine–carboxylate and amine–phosphate interactions for molecular dynamics simulations using the CHARMM and AMBER force fields. Journal of chemical theory and computation 12, 430–443 (2016).

79. Yoo, J. & Aksimentiev, A. New tricks for old dogs: improving the accuracy of biomolecular force fields by pair-specific corrections to non-bonded interactions. Physical Chemistry Chemical Physics 20, 8432–8449 (2018).

80. Yoo, J., Park, S., Maffeo, C., Ha, T. & Aksimentiev, A. DNA sequence and methylation prescribe the inside-out conformational dynamics and bending energetics of DNA minicircles. Nucleic Acids Research 49, 11459–11475 (2021).

81. Ohio Supercomputer Center. https://ror.org/01apna436 (1987).

82. The PyMOL Molecular Graphics System, Version 3.0 Schrödinger, LLC. Schrödinger, LLC.

83. Kumar, R. & Grubmüller, H. do_x3dna: A tool to analyze structural fluctuations of dsDNA or dsRNA from molecular dynamics simulations. Bioinformatics 31, (2015).

84. Lu, X. & Olson, W. K. 3DNA: a software package for the analysis, rebuilding and visualization of three-dimensional nucleic acid structures. Nucleic Acids Res 31, 5108–5121 (2003).

85. Jafilan, S., Klein, L., Hyun, C. & Florián, J. Intramolecular Base Stacking of Dinucleoside Monophosphate Anions in Aqueous Solution. J. Phys. Chem. B 116, 3613–3618 (2012).

86. Condon, D. E., Kennedy, S. D., Mort, B. C., Kierzek, R., Yildirim, I. & Turner, D. H. Stacking in RNA: NMR of Four Tetramers Benchmark Molecular Dynamics. J. Chem. Theory Comput. 11, 2729–2742 (2015).

87. Jacobson, H. & Stockmayer, W. H. Intramolecular Reaction in Polycondensations. I. The Theory of Linear Systems. The Journal of Chemical Physics 18, 1600–1606 (1950).

88. Nomidis, S. K., Kriegel, F., Vanderlinden, W., Lipfert, J. & Carlon, E. Twist-Bend Coupling and the Torsional Response of Double-Stranded DNA. Phys. Rev. Lett. 118, 217801 (2017).

89. Gao, X., Hong, Y., Ye, F., Inman, J. T. & Wang, M. D. Torsional Stiffness of Extended and Plectonemic DNA. Phys. Rev. Lett. 127, 028101 (2021).

90. Lipfert, J., Kerssemakers, J. W. J., Jager, T. & Dekker, N. H. Magnetic torque tweezers: measuring torsional stiffness in DNA and RecA-DNA filaments. Nat Methods 7, 977–980 (2010).

